# Targeting Outer Membrane β-barrel Proteins of *Burkholderia mallei* to design a Multi-epitope Vaccine Against Human Glanders

**DOI:** 10.64898/2026.05.25.727591

**Authors:** Jahnvi Kapoor, Amisha Panda, Sanjiv Kumar, Anannya Bandyopadhyay

## Abstract

*Burkholderia mallei*, a facultative intracellular Gram-negative pathogen, is the causative agent of glanders that primarily affects solipeds and is sporadically transmitted to humans. Current interventions mainly rely on antibiotics; however, increasing antimicrobial resistance and the lack of a licensed vaccine further complicate disease management. Surface exposed outer membrane β-barrel (OMBB) proteins serve as excellent targets for vaccine development. In the present study, a consensus-based computational framework was employed on the *B. mallei* turkey2 proteome that identified 59 OMBB proteins - including porins, TonB receptors, autotransporters, and efflux components. These OMBB proteins were leveraged to predict B- and T-cell epitopes which were manually curated, and mapped onto the corresponding protein models to identify surface-exposed epitopes with direct accessibility to the host immune cells. These epitopes were linked together to construct a multi-epitope vaccine (MEV) that was predicted to be antigenic, and soluble upon overexpression. The tertiary structure of the MEV was generated which was used for molecular docking with TLR4 and TLR2. Molecular dynamics simulation and flexibility analysis confirmed the structural stability of the MEV-TLR4/TLR2 complexes. In-silico immune simulation showed the capability of MEV to induce a strong immune response. Codon optimization and in-silico cloning were performed to evaluate its efficient expression in the *E. coli* host. The findings suggest that surface exposed OMBB proteins can serve as promising antigenic candidates for designing an MEV construct.

**Graphical Abstract:** 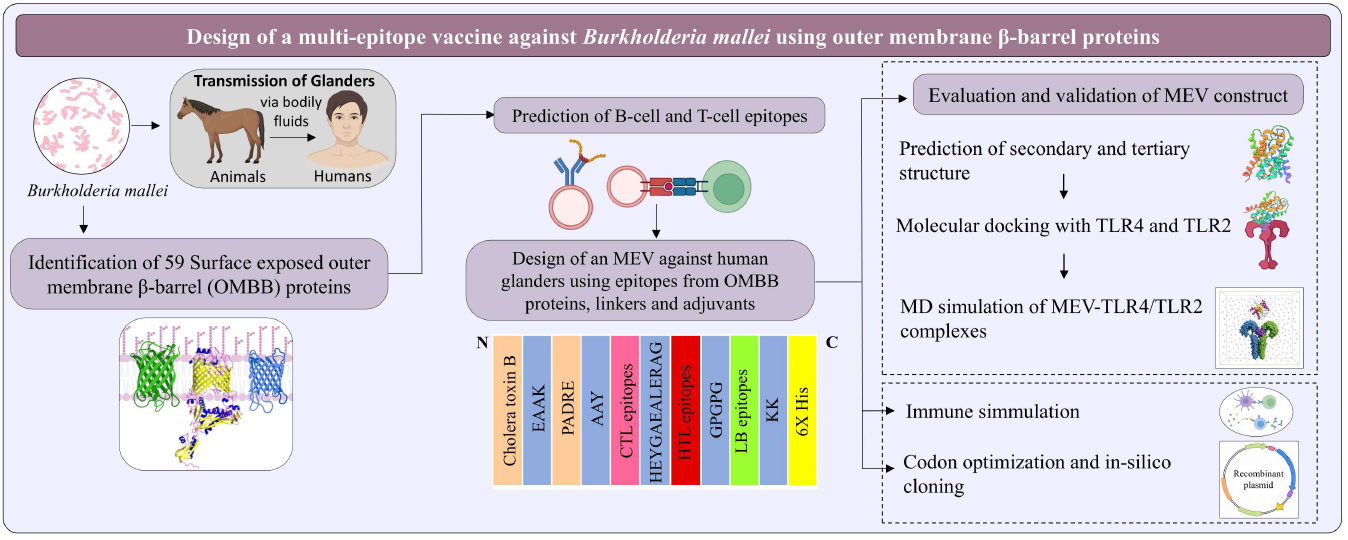

## 1. Introduction

Glanders is one of the oldest zoonotic diseases of equids caused by *Burkholderia mallei* (Bm) that can be transmitted to humans from solipeds^1^. The disease has been eradicated from the United States, while it is still prevalent in parts of Africa, Asia, the Middle East, Central America, South America, and endemic in Iraq, Pakistan, India, Mongolia, and parts of Brazil^2,3^. Glanders primarily affects equines, with reported seroprevalence of Bm in India ranging from 0.62% to 1.145% (2015-2018), while human infection occurs sporadically in laboratory workers and veterinary individuals^4^. Bm can be very hazardous in a laboratory setting; a pre-biosafety era report revealed one-half of the workers in a Bm research laboratory were infected within a year of exposure^5^. The disease is transmitted through contact with infected bodily fluids or via mucosal surfaces, such as the eyes, nose, and mouth. It is characterized by fever, malaise, rapid onset of pneumonia, pustules and abscesses, leading to death in 7-10 days without antibiotic treatment^6^. The etiological agent of glanders, Bm, is an obligate mammalian pathogen that can survive both in extracellular spaces and intracellularly within different host cell types^7^. The incubation period varies from ∼5 days to 12 weeks^8^. The bacteria enter via inhalation and localize in the upper and lower respiratory tract where they adhere to respiratory epithelial cells. They are then phagocytosed by alveolar macrophages via complement-mediated uptake. Presence of secretion systems promote intracellular survival and induce apoptosis. The infection spreads systemically through bloodstream, possibly via macrophage trafficking to lymph nodes^9^. Glanders diagnosis is challenging due to lack of specific serological tests. Bacterial culture, indirect haemagglutination test and qPCR are used for glanders diagnosis in humans^3^. Therapeutic interventions involve use of antibiotics, such as intravenous imipenem or ceftazidime followed by oral doxycycline and azithromycin for several months^10^. US Centres for Disease Control and Prevention has classified Bm as a bioterrorism agent due to its high pathogenicity (BSL-3), inherent resistance to many antibiotics, and ability to be dispersed via aerosols^11^. It has previously been used during World War II, where Japanese forces intentionally infected horses, civilians, and prisoners of war with Bm in Pingfang, China^12,13^. No vaccine is available for prevention of glanders in humans and animals. Thus, development of a vaccine is required to protect against potential use of the organism as a bioterrorism agent^10^.

*B. mallei* turkey2 is a Gram-negative, obligate aerobic coccobacillus (1.5-3.0 μ x 0.5-1.0 μ) belonging to the class β-proteobacteria. Bm belong to *Burkholderia pseudomallei* complex (Bpc) which consists of organisms whose genomes suggest a common ancestor similar to *B. pseudomallei* (Bp)^7^. Bm is a deletion-derived clone of Bp that has lost more than 1200 genes^14^. Bm has two circular chromosomes with a total genome size of about 5.6 Mb that encodes for 4641 proteins. The bacterium has a typical Gram-negative cell envelope comprising an outer membrane (OM), inner membrane (IM), and periplasmic space. The OM has lipopolysaccharides (LPS) with modified lipid A, supported by capsular polysaccharide, proteins, and phospholipids^15^. Diderm bacteria possess a diverse family of outer membrane β-barrel (OMBB) proteins^16,17^. β-barrels are cylindrical proteins with anti-parallel β-strands, typically comprising 8 to 36 β-strands^18^. They are involved in a variety of essential cellular functions, such as nutrient uptake, membrane biogenesis, OM assembly, adhesion, biofilm formation, efflux, and pilus formation^19^. In a previous study, ∼100 outer membrane proteins (OMPs) were identified in Bm expressed under different growth conditions using mass spectrometry and bioinformatics analysis. These OMPs included β-barrels, lipoproteins, Type VI secretion system constituent proteins, and other virulence-associated surface proteins^14^. Since Bm is considered as a potential biothreat pathogen, detailed knowledge of its OM and surface constituents is essential for the development of effective countermeasures. Although effective antibiotics are available for treating glanders, Bm exhibits both intrinsic and acquired mechanisms of antibiotic resistance. The presence of modified LPS confers intrinsic resistance to polymyxins, while efflux pumps contribute to resistance against tetracyclines, aminoglycosides, and fluoroquinolones^20^. Therefore, vaccination represents the most affordable and promising strategy to combat glanders. No licensed vaccine currently exists against Bm infection; however, multiple experimental vaccine candidates have shown promising protection in animal models. Many live attenuated vaccines (LAVs) have been created by deleting genes needed for metabolism, iron uptake, or secretion systems (e.g., ΔtonBΔhcp1 double mutant, ΔilvI, ΔpurM, ΔbatA)^21–24^. These vaccines were evaluated in mouse models providing protection and high sterilizing immunity; however, LAVs raise safety concerns, including the risk of tolerance induction, autoimmune reactions, and reversion to wild-type virulence^25,26^. Subunit vaccines, including purified proteins such as LolC^27^, OmpW^28^, OmpA-like proteins^29^, Hcp, BopA, and BimA^30^, have been evaluated along with adjuvants that were capable of inducing strong antibody responses but provided only partial protection. Advanced human vaccines against Bm, including nanoparticle-, and viral vector-based vaccines (e.g., AuNP-OmpW-LPS, Parainfluenza virus 5 carrying BatA protein etc.), have also shown promising immune responses and variable protection in animal models^31,32^. Despite significant progress in experimental vaccines against Bm, no vaccine has yet been commercially developed for human use. This highlights the need for novel strategies such as peptide based multi-epitope vaccines (MEVs). MEVs derived from multiple proteins offer broader antigen coverage, stronger immune responses, and reduced risk of immune escape compared to single-protein vaccines^33^. A previous study has proposed an MEV constructed using antigens from only two proteins, type VI secretion system tube protein (Hcp) and type IV pilus secretin protein, that were not exclusively surface exposed^34^. Since OMBB proteins are the most abundant proteins in the bacterial OM^17^, their surface exposed epitopes are readily accessible to the host immune system and are therefore more likely to induce stronger immune responses that can enhance the potential efficacy of the vaccine.

In the current study, we predicted 59 OMBB proteins in Bm turkey2 using a consensus-based computational approach^35,36^. AlphaFold 3 was utilized to generate structural models which were validated using four additional tools. Most of the OMBB proteins were annotated in NCBI except two hypothetical proteins. Sequence variations among 33 Bm strains were identified and mapped onto the structural models that identified variable surface exposed regions under selective pressure. To exploit the immunogenic potential of these proteins, B-cell, cytotoxic T-lymphocyte (CTL), and helper T-lymphocyte (HTL) epitopes were predicted, and linked together using appropriate linkers and adjuvants to design an MEV construct^37^. Population coverage analysis was performed for selected surface-exposed MHC-class I, and MHC-class II epitopes in glanders-endemic regions.

The physicochemical properties of the MEV were evaluated, and its secondary and tertiary structures were generated using multiple tools. Molecular docking was performed between the MEV, and human TLR4 and TLR2 to assess the binding interaction followed by molecular dynamics (MD) simulation to confirm the stability of the complex. Furthermore, immune simulation analysis was carried out to assess the potential of the MEV to elicit an immune response. Codon optimization and in-silico cloning were performed to demonstrate efficient recombinant protein expression of the MEV construct in *E. coli*. This study exploits the antigenic potential of surface exposed regions of OMBB proteins for the development of an effective vaccine and presents a potential vaccine candidate against Bm infection.

## 2. Materials and Methods

### 2.1 Prediction of outer membrane β-barrel (OMBB) proteins

*B. mallei* turkey2 is a well-annotated field isolate primarily isolated from horses that represents the natural genomic variation among circulating strains^38^. To predict OMBB proteins, amino acid sequences of 4641 proteins of *B. mallei* turkey2 (Assembly No. ASM234602v1) were downloaded from NCBI (http://www.ncbi.nlm.nih.gov/accessed) (accessed on 2 July, 2024)^39^. The computational framework designed to select OMPs is schematically represented in Figure 1. The workflow was adapted from the methodology described by Panda et al.^37^ and optimized for Bm. Ten computational tools were used to analyze the complete proteome of Bm. Pepstats tool (https://www.ebi.ac.uk/jdispatcher/seqstats/emboss_pepstats) of the EMBOSS package was used to determine the peptide length, molecular weight, charge, and isoelectric point (pI)^40^. Adhesins or adhesin-like proteins were predicted using SPAAN (https://sourceforge.net/projects/adhesin/files/SPAAN/spaan_64_bit.tar.gz/download) where the adhesin probability was calculated using a neural network model trained on known adhesins and non-adhesins^41^. SignalP 5.0 (https://services.healthtech.dtu.dk/services/SignalP-5.0/) was used to predict signal peptide and its cleavage position in the protein sequences based on a deep convolutional and recurrent neural network architecture^42^. Subcellular localization of the proteins was determined using CELLO v.2.5 (http://cello.life.nctu.edu.tw/)^43^ and PSORTb v3.0 (https://www.psort.org/psortb/) (all tools accessed on 4 July, 2024)^44^. CELLO v.2.5 uses support vector machines trained by multiple vectors based on n-peptide compositions^43^ while PSORTb v3.0 uses a probabilistic system for determining subcellular localization^44^.

**Figure 1.**
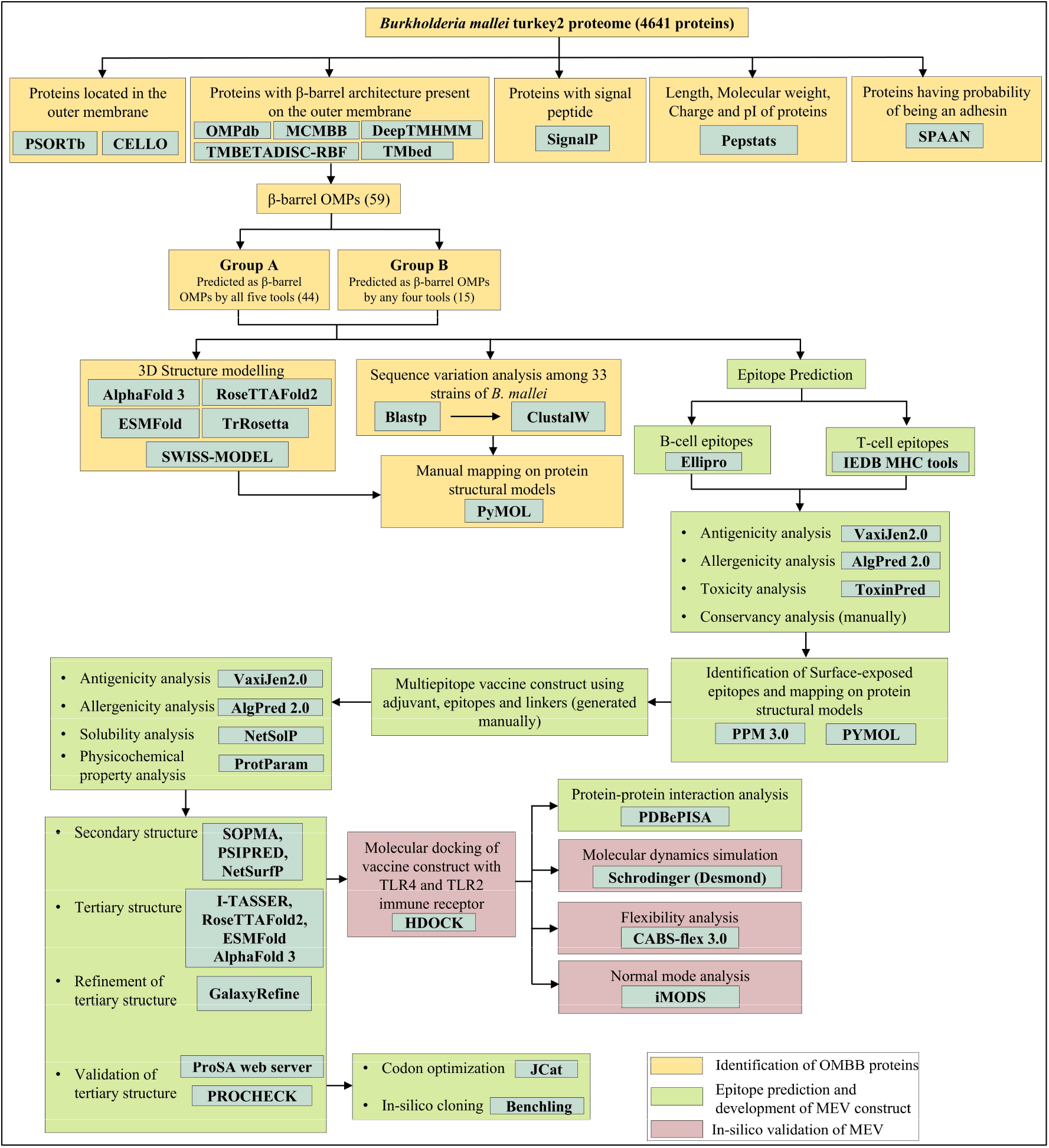
Computational framework for predicting OMBB proteins, and construction of a multi-epitope vaccine against Bm infection. Bm turkey2 proteome was analyzed in-silico using ten computational tools. OMBB proteins were subsequently categorized into two groups based on consensus predictions by five tools. Structural models were generated followed by sequence variation analysis among 33 Bm strains. B-cell and T-cell epitopes were predicted to construct an MEV using adjuvants, surface-exposed epitopes, and linkers. Tertiary structure of the MEV was predicted using four tools and best model was selected based on Ramachandran favoured residues. ProSA-web server and PROCHECK evaluated the overall quality of the tertiary structure. HDOCK was used for molecular docking of the MEV construct with TLR4 and TLR2 immune receptor. MD simulation was performed using Schrodinger. Fluctuation analysis and NMA were analyzed using CABS-flex and iMODS. PDBePISA was used to assess the interactions between MEV-TLR4/TLR2 complexes. Codon optimization and in-silico cloning was performed for optimal expression in host cell.

A consensus-based computational approach was employed to identify OMBB proteins using outputs from five OMBB prediction tools: one database - OMPdb (http://aias.biol.uoa.gr/OMPdb/)^45^, and four tools - MCMBB (http://athina.biol.uoa.gr/bioinformatics/mcmbb/)^46^, TMBETADISC-RBF (http://rbf.bioinfo.tw/~sachen/OMP.html)^47^, TMbed (https://github.com/BernhoferM/TMbed) (all tools accessed on 6 July, 2024)^48^, and DeepTMHMM (https://dtu.biolib.com/DeepTMHMM)^49^ (accessed on 13 April, 2025). OMPdb is a database of integral β-barrel OMPs from prokaryotes and eukaryotes^45^. MCMBB distinguishes β-barrel OMPs from globular proteins and α-helical membrane proteins^46^. TMBETADISC-RBF server predicts OMPs using radial basis function (RBF) network combined with position-specific scoring matrix (PSSM) profiles^47^. TMbed utilizes embeddings from protein language models (pLMs) to predict the propensity of each residue to form a transmembrane (TM) helix, β-strand, signal peptide, or other^48^. DeepTMHMM utilizes deep learning methods for the prediction of membrane topology of TM proteins^49^. Furthermore, proteins were classified based on the number of tools yielding positive results for a given protein sequence. This implied that higher confidence was assigned to those predicted as OMBB proteins by more tools. Proteins predicted as OMBBs by five tools were assigned to Group A, while those supported by four tools were assigned to Group B (Table S1). Group C, D, and E consisted of proteins predicted positive by any three, two, and one computational tool, respectively.

### 2.2 Structural modeling of predicted OMBB proteins

Structural models of the predicted proteins in Group A and Group B were retrieved from the AlphaFold database (https://alphafold.ebi.ac.uk/) (accessed on 10 August, 2025)^50^. Structure of proteins that were not available in the database were generated using AlphaFold 3 server (https://alphafoldserver.com/) (accessed on 11 August, 2025)^51^. To validate the structural models, we generated 3D structural models using additional four modeling tools: ESMFold (https://colab.research.google.com/github/sokrypton/ColabFold/blob/main/ESMFold.ipynb) (accessed on 18 August, 2025)^52^, SWISS-MODEL (https://swissmodel.expasy.org/) (accessed on 24 August, 2025)^53^, RoseTTAFold (https://robetta.bakerlab.org/) (accessed on 3 September, 2025)^54^ and TrRosetta (https://yanglab.qd.sdu.edu.cn/trRosetta/) (accessed on 4 September, 2025)^55^.

ESMFold predicts the protein structure based on embeddings from pLMs^52^, while SWISS-MODEL performs automated comparative modeling^53^. RoseTTAFold predicts 3D coordinates directly within the deep learning model using a three-track neural network^54^ while TrRosetta predicts protein structure using deep learning-based geometric constraints^55^. For each protein, the top ranked models were selected for further analysis based on highest confidence scores. The resulting atomic coordinate files were visualized using PyMOL^56^. Structural models were aligned using US-align (https://zhanggroup.org/US-align/) (accessed on 11 December, 2025)^57^ online web server to assess structural similarities and differences across models. US-align is a universal structure alignment tool designed to compare and superimpose 3D macromolecular structures. Alignments were visualized using PyMOL, and corresponding Root Mean Square Deviation (RMSD) values were recorded.

### 2.3 Amino acid sequence variation analysis across *B. mallei* strains

Amino acid sequences of predicted OMBB proteins were compared across 33 complete Bm proteomes to assess sequence variations (Table S4). Orthologous sequences for each protein were identified using BLASTP (E-value < 1.0E-03, bitscore > 100) (https://blast.ncbi.nlm.nih.gov/Blast.cgi?PROGRAM=blastp&PAGE_TYPE=BlastSearch&LINK_LOC=blasthome) (accessed on 16 December, 2025) and aligned using Clustal Omega^58^ for Multiple Sequence Alignment (MSA). MSA analysis identified amino acid variations among the orthologs, which were subsequently recorded in Table S5.

### 2.4 Prediction of B-cell and T-cell epitopes

ElliPro, a web server of the Immune Epitope Database (IEDB) (http://tools.iedb.org/ellipro/) (accessed on 23 December, 2025), was employed to predict both linear and conformational B-cell epitopes using the 3D structural model of the proteins^59^. The IEDB (Immune Epitope Database) web server predicts epitopes using structure- and sequence-based algorithms trained on experimentally validated epitope data^60^. Predictions of B-cell epitopes were carried out using default parameters. Epitopes longer than 50 amino acids were removed as they may introduce non-immunogenic flanking regions. MHC class-I (https://tools.iedb.org/mhci/) and MHC class-II (http://tools.iedb.org/mhcii/) prediction tools of the IEDB (accessed on 24 December, 2025)^61^ were used to predict cytotoxic T-lymphocyte (CTL) and helper T-lymphocyte (HTL) epitopes, respectively. CTL epitope prediction targeted 9-mer peptides with a percentile rank threshold of <0.5. The analysis included the following Human Leukocyte Antigen (HLA) alleles: HLA-A*01:01, HLA-A*02:01, HLA-A*11:01, HLA-A*24:02, HLA-B*07:02, HLA-B*08:01, HLA-B*15:01, HLA-B*44:02, HLA-C*04:01, HLA-C*07:02. HTL epitopes (15-mer peptides) were predicted with a percentile rank threshold <2 for the following HLA alleles: HLA-DRB1*01:01, HLA-DRB1*03:01, HLA-DRB1*04:01, HLA-DRB1*07:01, HLA-DRB1*08:02, HLA-DRB1*13:01, HLA-DRB1*13:02, HLA-DRB1*15:01.

### 2.5 Prediction of epitope homology with human proteome

Proteins lacking homology to human sequences are less likely to induce an autoimmune response. Given the fact, a comparison was made between *Homo sapiens* proteome (TaxID: 9606) and predicted peptides using the NCBI BLASTP database (https://blast.ncbi.nlm.nih.gov/Blast.cgi?PROGRAM=blastp&PAGE_TYPE=BlastSearch&LINK_LOC=blasthome) (accessed on 26 December, 2025)^62^. Peptides exhibiting an E-value exceeding 0.05 were considered non-homologous and suitable candidates for vaccine construction. Since BLASTP may have limited reliability for short peptides, k-mer analysis was performed using PepMatch^63^ as an additional validation step, and peptide sequences exhibiting zero mismatch (100% identity) with the human proteome were excluded.

### 2.6 Antigenicity, allergenicity and toxicity evaluation of predicted epitopes

After screening the predicted epitopes for homology with human proteome, the non-homologous epitopes were evaluated for antigenicity, allergenicity and toxicity. Antigenicity was assessed using the VaxiJen2.0 online web server (https://www.ddg-pharmfac.net/vaxijen/VaxiJen/VaxiJen.html) that predicts antigenicity using physicochemical properties of proteins with a threshold score of 0.4^64^. Allergenicity of the predicted epitopes was predicted using AlgPred 2.0 (https://webs.iiitd.edu.in/raghava/algpred2/), with default parameters^65^, that utilizes a combination of machine learning techniques, sequence-, motif-, and similarity-based approaches trained on an experimentally validated allergen dataset^65^. Toxicity of the selected epitopes was determined using ToxinPred server (http://crdd.osdd.net/raghava/toxinpred/) (all tools accessed on 26 December, 2025)^66^. ToxinPred employs machine learning models trained on experimentally validated toxic and non-toxic peptide datasets^66^. Epitopes identified as antigenic, non-toxic, and non-allergenic were selected for further analysis.

### 2.7 Conservation analysis and surface exposure of predicted epitopes

MSA was performed as described in Section 2.3 and the resulting data was used to evaluate the conservation of selected epitopes across 33 Bm strains. Conserved epitopes across 33 Bm strains were selected for further analysis to ensure their broad applicability and potential effectiveness. These epitopes were mapped onto the structural models of the predicted OMBB proteins to determine their location using PyMOL. PPM 3.0 (accessed on 2 August, 2026) was subsequently employed to validate the membrane orientation of the corresponding OMBB proteins and determine whether the selected epitopes were exposed on the extracellular surface^67^. Epitopes located on the extracellular loop (ECL) region were considered for subsequent analysis.

### 2.8 Population coverage analysis of T-cell epitopes

The population coverage of the selected CTL and HTL epitopes was evaluated using the IEDB Population Coverage tool to estimate the proportion of the population expected to respond to the selected epitopes based on their associated HLA class I and class II alleles. The analysis was performed using the population dataset of glanders-endemic regions^34,68^. The estimated population coverage was calculated to assess the potential of the vaccine-induced immune response across different populations.

### 2.9 Design of multi-epitope vaccine (MEV) construct

Antigenic, non-allergenic, non-toxic, conserved, and surface-exposed epitopes were selected for the construction of MEV. Three epitopes from each - CTL, HTL, and linear B-cell epitopes - were shortlisted based on their highest antigenicity scores. To construct the vaccine sequence, epitopes were joined together with appropriate peptide linkers^69,70^. CTL epitopes were connected using AAY linkers, while HTL epitopes were connected with GPGPG linkers. The last CTL epitope was connected to the first HTL epitope through HEYGAEALERAG linker^71^. B-cell epitopes were linked using KK linkers. To enhance immunogenicity, two adjuvants- Cholera toxin B subunit (CTB) (accession no.: P01556) and PADRE sequence were inserted at the N-terminus with the help of EAAAK linkers. Finally, a complete vaccine sequence was obtained by adding a hexa-histidine tag at the C-terminus through a KK linker for purification^72^.

### 2.10 Evaluation of multi-epitope vaccine construct

The MEV was evaluated using four computational tools - ProtParam (http://web.expasy.org/protparam/) was used to analyze physicochemical properties^73^, VaxiJen 2.0 for antigenicity assessment^64^, NetSolP 1.0 (https://services.healthtech.dtu.dk/services/NetSolP-1.0/) for solubility prediction^74^ and AlgPred 2.0 for allergenicity evaluation^65^ (all tools accessed on 3 August, 2026). NetSolP employs a sequence-based prediction method to predict the propensity of a protein to be soluble upon overexpression^74^. A previous study has demonstrated that *B. pseudomallei* OmpA proteins are immunogenic in mice and are potential vaccine candidates^29^. Thus, Omp7 (OmpA domain containing protein, Accession no. ACN64871.1) of *B. pseudomallei* was used as positive control.

### 2.11 Structure prediction and validation of the MEV construct

SOPMA (Self-Optimized Prediction method With Alignment) secondary structure prediction tool (http://npsa-pbil.ibcp.fr/cgibin/npsa_automat.pl?page=/NPSA/npsa_sopma.html)^75^, PSIPRED 4.0 (https://bioinf.cs.ucl.ac.uk/psipred/)^76^, and NetSurfP 2.0 (https://services.healthtech.dtu.dk/services/NetSurfP-2.0/)^77^ (all tools accessed on 4 August, 2026) were used to predict the secondary structures of MEV construct. SOPMA uses MSA–based consensus scoring, PSIPRED utilizes PSI-BLAST-derived evolutionary profiles, and NetSurfP uses machine-learning models to assess the structure of the target protein and predict the formation of α-helices, beta-turns, and random coils. Intrinsically disordered regions (IDRs) of the MEV construct were predicted using the PrDOS web server (http://prdos.hgc.jp/cgi-bin/top.cgi) that predicts residue-wise disorder propensity using a support vector machine (SVM)-based approach^78^. Tertiary structure of MEV was generated using four modeling tools - I-TASSER tool (https://zhanggroup.org/I-TASSER/), AlphaFold 3, ESMFold, and RoseTTAFold2 (all tools accessed on 5 August, 2026)^79^, followed by refinement of the tertiary structure using GalaxyRefine web server (http://galaxy.seoklab.org/) (accessed on 6 August, 2026) to improve model quality^80^. I-TASSER (Iterative Threading ASSEmbly Refinement) predicts a protein’s 3D structure by threading the sequence onto known templates, and iteratively assembles the fragments^79^.

GalaxyRefine refines loop or terminus regions in the initial protein 3D model by performing ab initio calculations^80^. The model with the highest proportion of residues in the favored region of the Ramachandran (R) plot was refined again to obtain the best model. The quality of the MEV tertiary structure model was assessed using the PROCHECK (https://saves.mbi.ucla.edu/)^81^ and the ProSA-web server (https://prosa.services.came.sbg.ac.at/prosa.php) (both tools accessed on 6 August, 2026)^82^. PROCHECK performs R-plot analysis, while ProSA (Protein Structure Analysis) web server calculates a Z-score by comparing the input protein model against known experimentally determined structures in the Protein Data Bank (PDB)^82^. The R-plot and the Z-score analysis provide insights into the stereochemical quality and overall stability of the predicted structure, respectively.

### 2.12 Molecular docking of the MEV construct with human TLR4 and TLR2 immune receptors

Molecular docking of MEV construct was performed with human TLR4 and TLR2 to evaluate the interaction between the MEV construct and host immune receptors. TLR4 and TLR2 plays a key role in recognizing *Burkholderia* and initiating immune responses^83^. The atomic coordinate files of human TLR4 (PDB ID: 3FXI) and TLR2 (PDB ID: 6NIG) were obtained from the RCSB PDB. Docking was performed using the HDOCK server (http://hdock.phys.hust.edu.cn/) (accessed on 7 August, 2026). The HDOCK server utilizes a hybrid algorithm to perform protein-protein and protein-DNA/RNA docking that combines template-based modeling and template-free docking methods. The structural models of docked complexes were validated by calculating interaction energetics using the PRODIGY (PROtein binDIng enerGY prediction) server^84^ (https://rascar.science.uu.nl/prodigy/), which provides Gibbs free energy (ΔG, kcal mol^-1^) or dissociation constant (K_d_, M) for intermolecular interactions typically within a 5.5 Å distance cutoff. The resulting MEV-TLR4/TLR2 complexes were evaluated for interacting amino acid residues using PDBePISA (https://www.ebi.ac.uk/pdbe/pisa/) (accessed on 8 August, 2026)^85^ and the docked complexes were visualized in PyMOL.

### 2.13 Protein flexibility assessment of MEV-TLR4/TLR2 complexes

CABS-flex v3.0 (https://lcbio.pl/cabsflex3/) (accessed on 8 August, 2026) was used to perform rapid protein flexibility analysis using the coarse-grained CABS model^86^. The MEV-TLR4/TLR2 coordinate files were submitted to the server for a simulation equivalent to 10ns with default settings. The output generated ten modeled clusters and a root mean square fluctuation (RMSF) profile for per-residue fluctuations. It also provided RMSD values for medoid (representative structure of a cluster) compared to input reference structure. The iMODS server (https://imods.iqf.csic.es/) (accessed on 8 August, 2026) was employed for normal mode analysis (NMA)-based simulation, assuming that the lowest-frequency modes represent the largest, functionally relevant motions^87^. Docked PDB structures were submitted with default parameters, and stability was evaluated based on deformability, B-factor, eigenvalues, variance, covariance map, and elastic network analyses.

### 2.14 MD simulation of MEV-TLR4/TLR2 complexes

MD simulation of the docked MEV-TLR4/TLR2 complexes were performed using the Desmond module in the Schrodinger suite 2025-3^88^ to assess their structural stability, conformational flexibility, and dynamic interactions under near-physiological conditions. The system was prepared using the System Builder, where each complex was solvated with TIP3P water molecules enclosed within an orthorhombic simulation box. The OPLS4 force field was applied at physiological pH 7.4 with Na^+^ & Cl^−^ ions and a salt concentration of 0.15M. MD simulation was performed for 100 ns under NPT conditions at a bar pressure of 1.01325 and a constant temperature of 300K. The resulting trajectories were analyzed using Simulation Interactions Diagram (SID) to obtain RMSD plot for each protein-protein complex.

### 2.15 Immune simulation of the MEV construct

The online web server C-ImmSim (https://kraken.iac.rm.cnr.it/C-IMMSIM/) (accessed on 8 September, 2026) was employed to study the extent and type of immune responses elicited by the MEV construct in humans^89^. The server simulates the immune system using position-specific scoring matrices (PSSMs) and machine-learning approaches and evaluates the interactions between the antigen and various immune cells. Simultaneously, it simulates three different anatomical regions in mammals: bone marrow, thymus, and lymph nodes^90^. The vaccine was administered in three doses at three different time steps: 1, 84, and 168, respectively, with each time step equivalent to 8 hours. The simulation parameters were set as follows: random seed 12345, simulation volume of 50, and simulation steps of 1050. Six human HLA alleles were selected for HLA-antigen interaction including HLA-A*01:01, HLA-A*02:01, and HLA-B*07:02, HLA-B*08:01, HLA-DRB1*01:01, and HLA-DRB1*03:01.

### 2.16 Codon optimization and in-silico cloning

Codon optimization of the designed MEV construct was performed to ensure efficient heterologous expression in *E. coli* K-12 using Java Codon Adaptation Tool (JCat) (http://www.jcat.de/) (accessed on 9 August, 2026)^91^. This improves the codon usage according to the codon preference of *E. coli* while maintaining the amino acid sequence of the vaccine construct. To facilitate efficient transcription and translation, rho-independent transcription terminators, prokaryotic ribosome binding sites, and restriction enzyme (BamHI and EcoRI) cleavage sites were avoided. The codon adaptation index (CAI) and GC content of the optimized nucleotide sequence were evaluated.

For in-silico cloning, BamHI and EcoRI restriction enzyme recognition sites were incorporated at the 5′ and 3’ termini of the optimized nucleotide sequence, respectively. Subsequently, the optimized MEV gene was amplified using suitable primers and inserted into the pET-28a(+) expression vector between the BamHI and EcoRI restriction sites using the molecular cloning tool available in Benchling (https://www.benchling.com/)^92^. The recombinant plasmid construct was assessed to confirm the correct orientation and in-frame insertion of the MEV sequence for potential recombinant protein expression in *E. coli*.

## 3. Results

### 3.1 Prediction of outer membrane β-barrel (OMBB) proteins

Although several OMPs have previously been identified in *Burkholderia* species, a comprehensive proteome-wide approach for the identification of OMBB proteins has not been reported in Bm. Therefore, we applied a consensus-based computational approach to identify the complete repertoire of OMBB proteins in Bm turkey2 genome, which encodes 4,641 proteins (Figure 1). The framework utilized ten computational tools: Pepstats, SPAAN, SignalP, CELLO, PSORTb, OMPdb, MCMBB, DeepTMHMM, TMBETADISC-RBF, and TMbed. Predictions from all the tools were integrated for each protein sequence (Table S1). Outputs from five OMBB-specific prediction tools - OMPdb, MCMBB, DeepTMHMM, TMBETADISC-RBF, and TMbed - were considered for OMBB identification (Table 1). Through consensus-based screening criteria, 4641 proteins were categorized into five groups (A-E), where Group A included proteins identified as OMBB by all five prediction tools, Group B contained proteins predicted positive by any four tools, Group C had proteins predicted positive by any three tools and so on. For high confidence in OMBB prediction, only Group A and B proteins were considered for further analysis. Manual curation based on structural models generated using AlphaFold 3 confirmed 59 OMBB proteins in both the groups where Group A included all 44 OMBB proteins, while Group B initially contained 19 proteins, of which only 15 were β-barrels. Literature-based evidences revealed that these 59 OMBB proteins comprised three OMPs previously identified experimentally in Bm ATCC 23344, five proteins with known β-barrel architectures, four conserved immunogenic proteins reported in other *Burkholderia* species, 20 with experimentally characterized functional homologs in other Gram-negative bacteria including *Burkholderia*, 25 proteins only annotated in NCBI/UniProt, and two novel OMBB proteins identified in this study (Table S2). Structural models of Group A and Group B proteins were generated using four additional modeling tools: ESMFold, SWISS-MODEL, RoseTTAFold, and TrRosetta. To assess the structural consistency, structural models generated using five tools were aligned using the US-align server, and the resulting RMSD values were found to be low ranging from 1.28 to 4.98 Å (Table S3). Group C, D, and E consisted of 22, 150, and 1321 proteins, respectively, based on consensus-based screening. Proteins in Groups C-E were excluded from further analysis as they were predicted positive by less than three computational tools, resulting in lower β-barrel prediction confidence compared to Groups A and B.

**Table 1:**
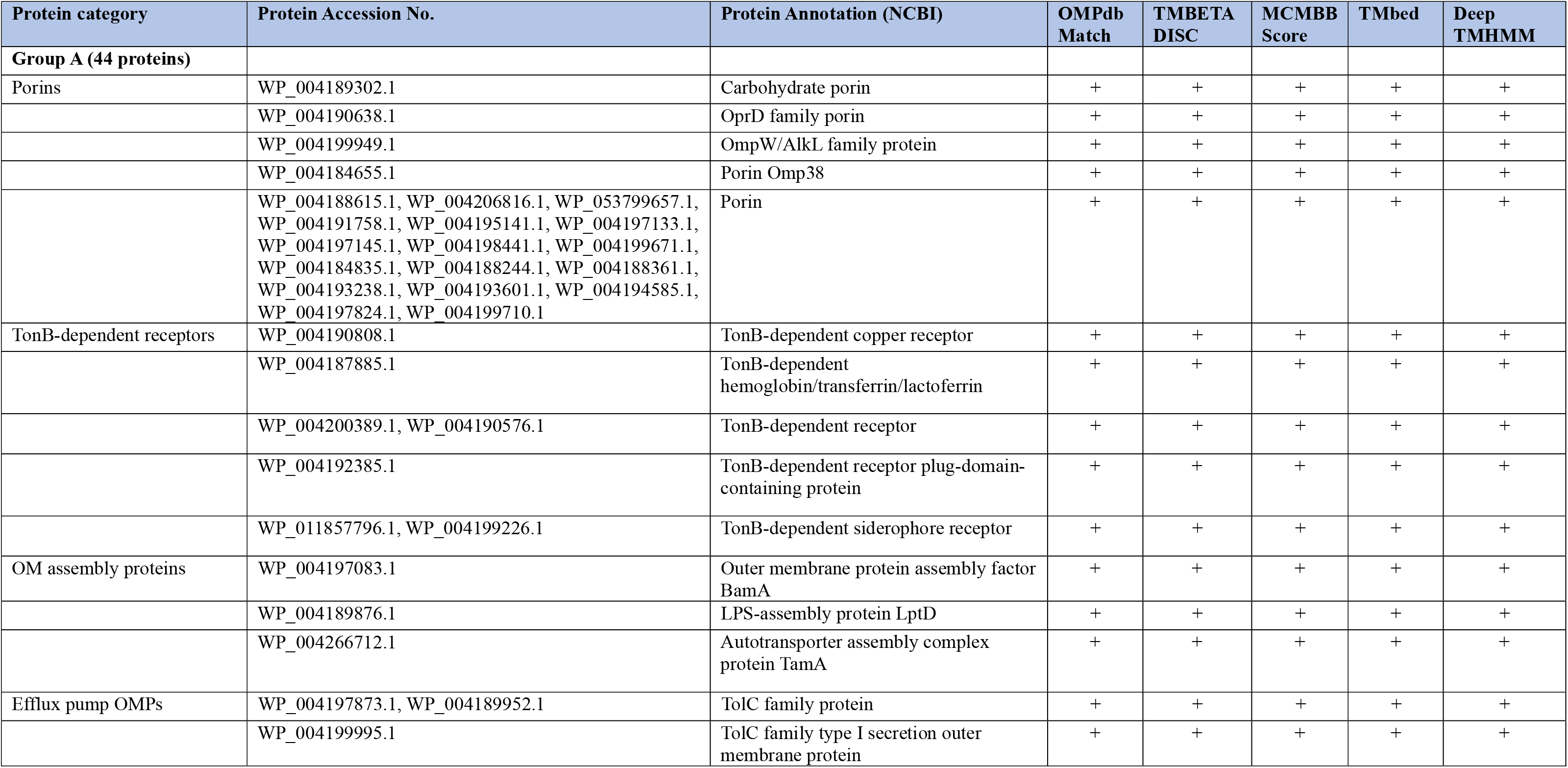
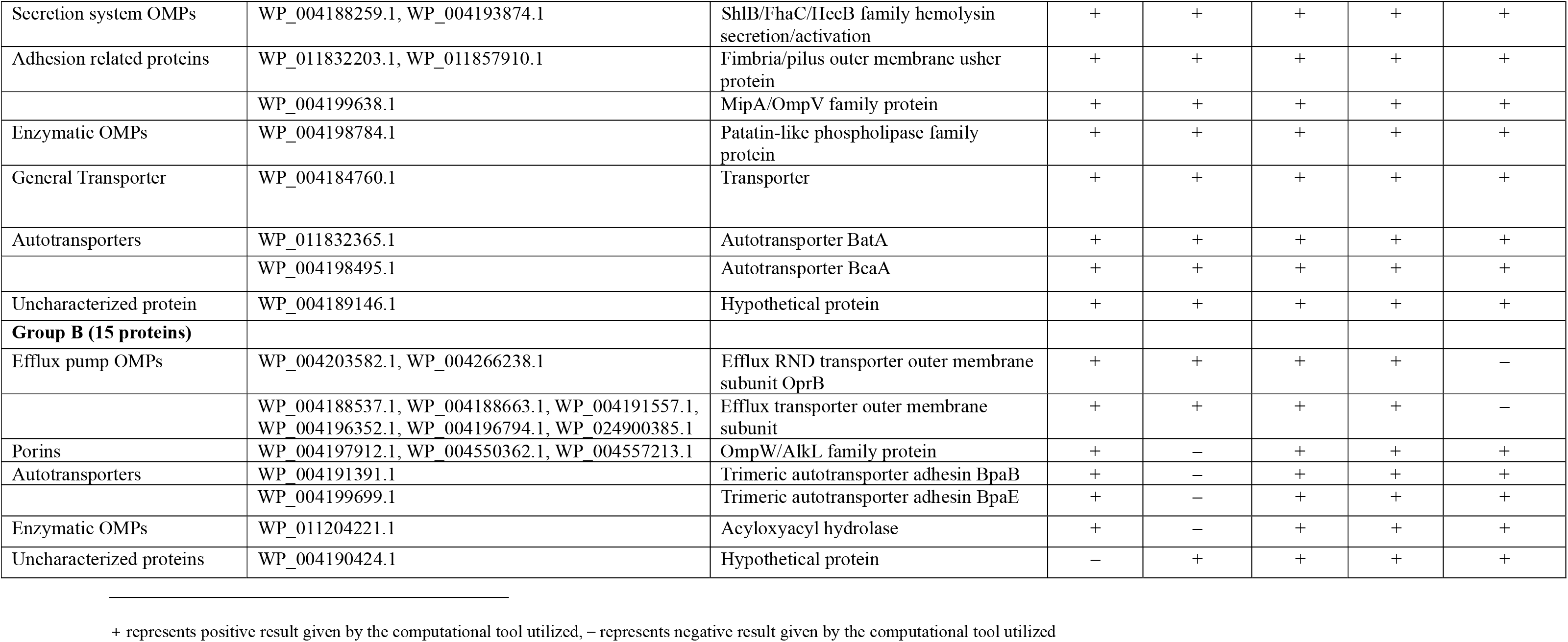
List of OMBB proteins predicted in *B. mallei* turkey2 using consensus-based computational approach.

These 59 OMBB proteins were divided into ten categories - porins, TonB-dependent receptors, OM assembly proteins, efflux pump OMPs, secretion system OMPs, adhesion related proteins, enzymatic OMPs, general transporter, autotransporters, and uncharacterized proteins - based on their NCBI annotation (Table 1).

#### 3.1.1 Porins

Out of 59 proteins, Group A has 21 OMBB proteins which are annotated as porin family proteins in NCBI and UniProt (Table 1, Figure 2), while Group B has three proteins - WP_004197912.1, WP_004550362.1, and WP_004557213.1 - annotated as OmpW family proteins (Figure S1). Group A includes carbohydrate porin, OprD family porin, OmpW/AlkL family protein, porin Omp38, and general porins. In 2011, Schell et. al identified only 13 porins in *B. mallei* ATCC 23344 using LC-MS/MS proteome analysis where WP_004189302.1 and WP_004184655.1 were detected under all seven growth conditions^14^. We predicted WP_004189302.1, OprB family porin, to have a 16-stranded β-barrel with molecular weight of 53.07 kDa. In other Gram-negative bacteria, OprB is involved in carbohydrate-specific transport of glucose, mannitol, fructose and glycerol^93^. Further, WP_004184655.1, an Omp38 family porin, is known to form a 16 stranded β-barrel that allows passive diffusion of small hydrophilic molecules and contributes to intrinsic antibiotic resistance in *B. pseudomallei*^94^. WP_004190638.1 is annotated as OprD family porin that was predicted to form an 18-stranded β-barrel. OprD family porins are substrate-specific OMPs that facilitate the uptake of basic amino acids (such as arginine) and structurally related antibiotics (like carbapenems) in *Pseudomonas aeruginosa*^95^. Structural modeling of WP_004199949.1 revealed an eight-stranded β-barrel belonging to OmpW family which is involved in bacterial attachment to host epithelial cells. It has been identified as a protective vaccine antigen in various *Burkholderia* species, including *B. cenocepacia*, *B. multivorans*, *B. thailandensis*, and *B. pseudomallei*^28,96^. Rest of the 17 proteins in Group A, including WP_004188615.1, WP_004206816.1, WP_053799657.1, WP_004191758.1, WP_004195141.1, WP_004197133.1, WP_004197145.1, WP_004198441.1, WP_004199671.1, WP_004184835.1, WP_004188244.1, WP_004188361.1, WP_004193238.1, WP_004193601.1, WP_004194585.1, WP_004197824.1, and WP_004199710.1, were annotated as general porins. We predicted these porins to have a 16-stranded β-barrel architecture using five structural modeling tools. Three OmpW family proteins in Group B form eight-stranded β-barrel structure generated using five modeling tools. WP_004550362.1 is known to be a highly conserved protein in *Burkholderia* species and serve as cross protective antigen against the bacterial infection^28^.

**Figure 2.**
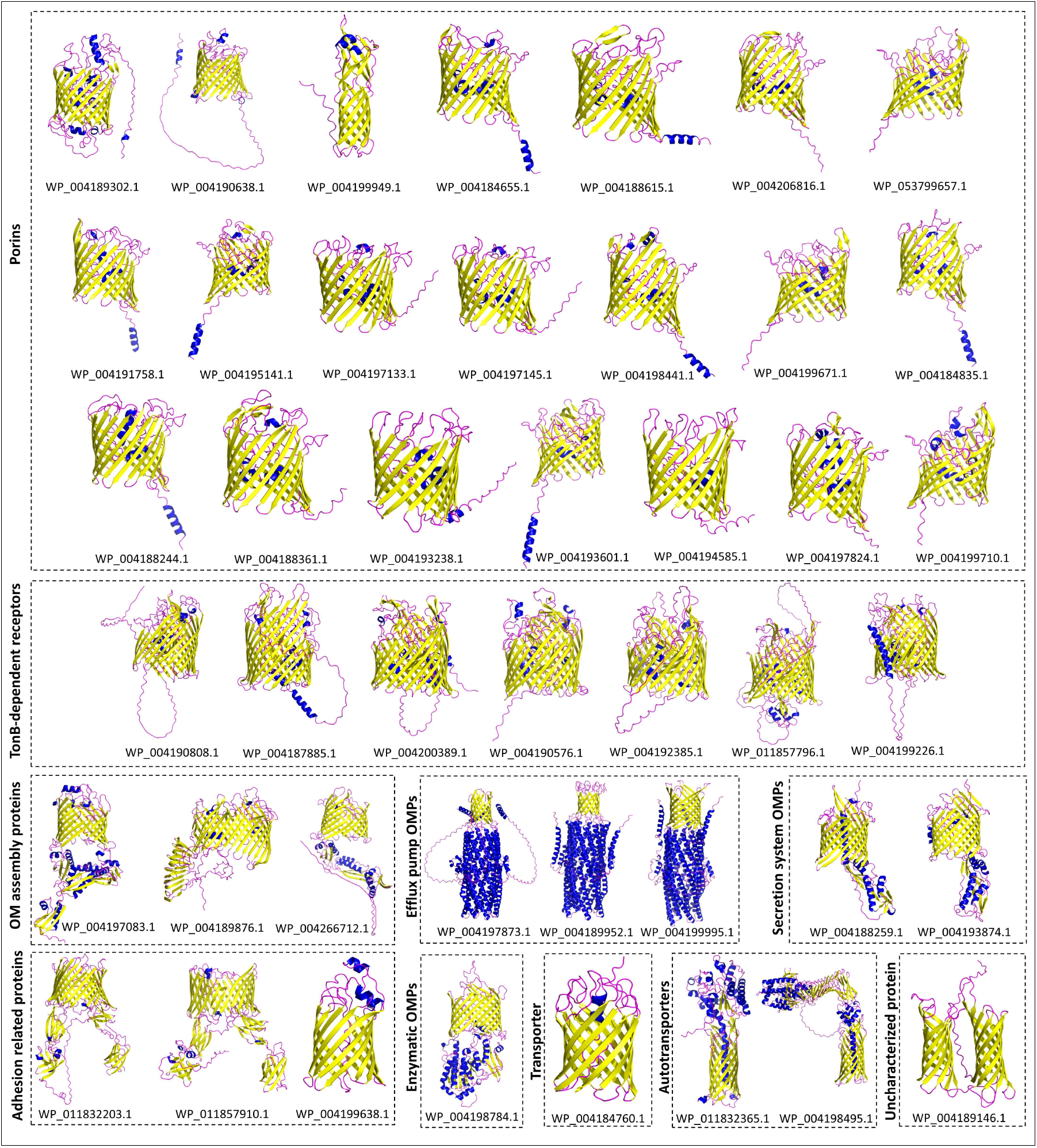
**Structural models of OMBB proteins in Group A generated using AlphaFold 3**. 44 proteins were predicted as β-barrels by all the five OMBB prediction tools – OMPdb, MCMBB, TMBETADISC-RBF, TMbed, and DeepTMHMM.

#### 3.1.2 TonB-dependent receptors

We identified seven proteins - WP_004190808.1, WP_004187885.1, WP_004200389.1, WP_004190576.1, WP_004192385.1, WP_011857796.1, and WP_004199226.1 - annotated as TonB-dependent proteins (TBDP) in NCBI which were classified in Group A (Table 1). In a previous study, by Schell et. al, only three TBDP were identified in Bm ATCC 23344 including WP_004190808.1^14^. Structural models of these seven proteins, generated using five tools, revealed a 22-stranded β-barrel containing a plug domain (Figure 2). In TBDPs, plug domain undergoes conformational changes to regulate the opening and closing of the pore^97^. Presence of multiple TonB receptors is required for the transport of various molecules like iron, copper, hemoglobin, transferrin, lactoferrin, and vitamin B12 in Bm^14,98^.

#### 3.1.3 OM assembly proteins

Three OM assembly proteins include WP_004197083.1, WP_004189876.1, and WP_004266712.1 were identified in Group A (Table 1). WP_004197083.1 is annotated as outer membrane protein assembly factor BamA. BamA is a member of Omp85 family, previously identified by Schell et. al^14^. We generated the structural model of BamA having a 16-stranded β-barrel architecture and five polypeptide transport-associated (POTRA) domains towards periplasmic side (Figure 2). It is an essential component of *<u>β</u>*-barrel <u>a</u>ssembly <u>m</u>achine (BAM) complex. The β-barrel contains a lateral opening between strands β1 and β16 that facilitates the insertion of OMPs into the OM of Gram-negative bacteria^99^. BamA is a conserved immunogenic protein, showing >86% identity across *Burkholderia* species^100^. WP_004189876.1 is annotated as lipopolysaccharide (LPS) transport protein D (LptD) that forms the OM component of Lpt complex. Structural model of WP_004189876.1, generated using five tools, has a 26-stranded β-barrel with a distinctive periplasmic β-jelly roll domain. *B. cenocepacia* and other related species are highly resistant to antimicrobial peptides, primarily depending on composition of LPS, which is transported to OM via Lpt complex^101^. Another OM assembly protein identified in this study is WP_004266712.1 which is annotated as Translocation and Assembly Module subunit A (TamA). TamA is an Omp85 family protein having a 16-stranded β-barrel with three POTRA domains. In Gram-negative bacteria, it is involved in folding and insertion of specific autotransporters, adhesins, and secretion systems into the OM^102^.

#### 3.1.4 Efflux pump OMPs

Three proteins - WP_004197873.1, WP_004189952.1, and WP_004199995.1 - were annotated as TolC family proteins in Group A. Additionally, eight efflux proteins were present in Group B including WP_004203582.1, WP_004266238.1, WP_004188537.1, WP_004188663.1, WP_004191557.1, WP_004196352.1, WP_004196794.1, and WP_024900385.1 (Table 1). These TolC efflux family proteins were predicted to form a trimeric 12-stranded α/β-barrel where each subunit contains four β-strands, spanning the OM of Bm (Figures S1, and S2). TolC is a key component of tripartite systems such as Resistance-Nodulation-Division (RND) efflux pumps in Gram-negative bacteria^103^. A previous study has identified 14 RND homologs in *B. cenocepacia* which are responsible for intrinsic resistance against antibiotics such as β-lactams, fluoroquinolones, and aminoglycosides^104^. RND efflux pumps have also been identified in *B. thailandensis* that contribute to control the permeability barrier of the OM^105^. Presence of these many efflux proteins in Bm might contribute to the intrinsic resistance observed in the bacterium^20^.

#### 3.1.5 Secretion system OMPs

WP_004188259.1, and WP_004193874.1, classified into Group A, are annotated as ShlB/FhaC/HecB family hemolysin secretion/activation proteins. These proteins form 16-stranded β-barrels as predicted by the five modeling tools. ShlB proteins are components of Type V Secretion System (T5SS) which are known to secrete and activate the hemolysin ShlA, a pore-forming toxin involved in bacterial virulence in *Paraburkholderia kururiensis*^106,107^.

#### 3.1.6 Adhesion related proteins

Three OMBB proteins in Group A - WP_011832203.1, WP_011857910.1, and WP_004199638.1 - were predicted to be adhesion related proteins based on their NCBI and UniProt annotation. WP_011832203.1, and WP_011857910.1 are annotated as fimbria/pilus outer membrane usher proteins. In 2024, Schully et. al identified Yersinia-like fimbrial (YLF) gene cluster in *B. pseudomallei* isolates from Ghana^108^. The gene cluster encodes proteins involved in the formation of fimbriae (pili), including an OM usher protein that facilitates their assembly^108^. These proteins are known to be a part of chaperone-usher pili system in Gram-negative bacteria^109^. Structural modeling revealed that both the proteins have a 24 stranded β-barrel domain, a plug domain, an N-terminal domain, and two C-terminal domains, consistent with the general structure of usher proteins^109^. WP_004199638.1 is an MipA/OmpV family protein forming a 12-stranded β-barrel.

OmpV family protein helps in adhesion and invasion of *Salmonella typhimurium* to intestinal epithelial cells and thus plays a vital role in the pathogenesis^110^. MipA in *B. thailandensis* is identified as part of the BesRS regulon, which is activated in response to cell envelope stress such as exposure to antibiotics like polymyxin B leading to antibiotic resistance phenotypes^111^.

#### 3.1.7 Enzymatic OMPs

Two proteins - WP_004198784.1 (Group A), and WP_011204221.1 (Group B) - were predicted as enzymatic OMPs based on their NCBI annotation. WP_004198784.1 is annotated as patatin-like phospholipase family protein. Hanson et. al predicted the structure of phospholipase A homolog (BpPlA) in *B. pseudomallei.* The protein has three key domains - N-terminal patatin-like (PL) domain, POTRA domain (periplasmic scaffold), and C-terminal 16 stranded β-barrel (OM anchor)^112^. Structural model of WP_004198784.1 consists 16-stranded β-barrel similar to phospholipase protein homolog in *B. pseudomallei* (Figure 2). It plays an important role in hydrolysis of membrane phospholipids like phosphatidylethanolamine, phosphatidylglycerol, phosphatidylserine, and cardiolipin in multiple bacterial species. It is a membrane-targeting enzyme that becomes active only after interaction with ubiquitin^113^. WP_011204221.1 is annotated as acyloxyacyl hydrolase in NCBI and Lipid A deacylase in UniProt. We identified WP_011204221.1 to form an eight-stranded β-barrel having a PagL domain searched in conserved domain database (CDD). PagL is a lipid A 3-O-deacylase that removes an acyl chain from lipid A and converts penta-acylated lipid A to tetra-acylated lipid A in *B. pseudomallei*. Deacylated lipid A reduces TLR4 immune recognition thus helps the bacteria to evade innate immunity^114^.

#### 3.1.8 Transporter

Another protein, WP_004184760.1, in Group A is annotated as a transporter protein in both NCBI and UniProt. It was predicted to have a 12-stranded β-barrel structure with molecular weight of 64.32 kDa (Figure 2). The specific substrate transported by this protein remains unknown. However, conserved domain analysis indicated that the protein contains the Pfam13557 domain, which is associated with phenol degradation and has also been reported in *Acinetobacter baumannii* as mentioned in InterPro^115^.

#### 3.1.9 Autotransporters

Among the 59 identified OMBB proteins, four were annotated as autotransporters in the NCBI and UniProt, where two proteins classified in Group A (WP_011832365.1, and WP_004198495.1) and two in Group B (WP_004191391.1, and WP_004199699.1) (Table 1). Autotransporter (AT) proteins are one of the largest classes of virulence factors in Gram-negative bacteria^116^. BatA (WP_011832365.1) consists of 610 amino acid residues with a molecular weight of 64.32 kDa.

BatA is a classical AT known to have a 12-stranded β-barrel domain and an extracellular passenger domain (Figure 2). In 2018, Lafontaine et. al showed that BatA is a highly conserved protein that aids in intracellular survival of Bm^24,31^. The presence of a surface-exposed N-terminal passenger domain makes BatA a promising vaccine candidate. Delivery of BatA using a recombinant Parainfluenza virus 5 (PIV5) in mice models can elicit strong immune responses and protect against Bm infection^31^. Another classical AT, BcaA (WP_004198495.1), comprises 1131 amino acids and has an estimated molecular weight of 116.05 kDa. BcaA model, generated using five modeling tools, contains a 12-stranded β-barrel domain anchored to OM and a larger passenger domain compared to BatA (Figure 2). The N-terminal passenger domain harbors a serine protease (subtilisin-like) domain^117^. Experimental findings determined that BcaA acts as an “invasin”, as mutants lacking BcaA in *B. pseudomallei* have significantly reduced ability to invade epithelial cells^117^. Further, two trimeric autotransporter adhesins (TAAs) included BpaB (WP_004191391.1), and BpaE (WP_004199699.1) (Figure S1). *bpaB* and *bpaE* genes have been identified in *B. pseudomallei* located on chromosome I and II respectively. These proteins are known to have a 12-stranded β-barrel embedded in OM and a YadA-like passenger domain towards extracellular region. These autotransporters contribute to host cell adhesion, and invasion, leading to *B. pseudomallei* infection^118^.

#### 3.1.10 Uncharacterized protein

Two proteins, one from Group A (WP_004189146.1) and other from Group B (WP_004190424.1), are annotated as hypothetical proteins in NCBI. WP_004189146.1 consists of 340 amino acid residues with molecular weight of 37.28 kDa. The structural model generated using five modeling tools revealed that it is a double barrel protein with eight β-strands in each barrel (Figure 2). Double β-barrel structures have previously been reported in *Burkholderia oklahomensis* that consist of two homologous domains, each forming a β-barrel composed of five antiparallel β-strands^119^.

WP_004190424.1 is annotated as hypothetical protein in NCBI and as porin in UniProt. It was predicted as an OMBB protein having an 18-stranded β-barrel architecture (Figure S1). It is 432 amino acid long with molecular weight of 46.8 kDa. Sequence variation analysis of both the proteins revealed that they are highly conserved protein with no amino acid variation among 33 strains of Bm, thus can serve as protective antigens for vaccine development.

### 3.2 Amino acid sequence variation analysis

Amino acid sequence variation analysis was performed across 33 Bm strains for the identified OMBB proteins (Table S4). These variations were mapped onto the corresponding structural models and visualized in PyMOL. These variations were observed to be distributed throughout the protein models including extracellular (ECL), transmembrane (TM), and intracellular (ICL) regions. Amino acid sequence variations within the ECL region indicate that these regions are under continuous selection pressure at the host-pathogen interface. Variations in these regions may help the pathogen to evade immune recognition and enhance bacterial adaptability by modifying these interactions within the host environment. Out of 59 OMBB proteins, 29 proteins were found to be highly conserved with no variation, while 11 proteins (WP_004188244.1, WP_004190808.1, WP_004194585.1, WP_004198495.1, WP_004198784.1, WP_004206816.1, WP_011832203.1, WP_011857796.1, WP_004188537.1, WP_004188663.1, and WP_004191391.1) were found to have more than 100 amino acid variations each (Table S5).

### 3.3 Prediction and evaluation of B-cell epitopes

ElliPro predicted linear B-cell epitopes (LBEs) from all the 59 OMBB proteins. A total of 656 LBE were initially predicted by ElliPro with a score (protrusion index) above 0.5 and selected for further analysis. Epitopes with >50 amino acid residues were discarded as they extend beyond surface-exposed loops into buried transmembrane regions. Remaining LBEs were then screened for homology with human proteome that retained only 173 non-homologous LBEs. Further, 25 non-homologous LBEs were found to be antigenic, non-allergenic, non-toxic, and conserved across 33 Bm strains. These LBEs were mapped onto the corresponding protein structural models embedded in a bacterial OM using PPM 3.0 server to identify surface exposed epitopes. Eight LBEs were located in the ECL region, and top three epitopes based on their ElliPro scores were considered for inclusion in the MEV design (Table 2). These three LBEs belonged to 3 OMBB proteins: WP_004187885.1, WP_004190576.1, and WP_004184760.1 (Figure 3). ElliPro also predicted 294 conformational B-cell epitopes, out of which top 20 epitopes are listed in Table S6 based on their protrusion indexes. Since the antigenicity of discontinuous conformational epitopes depends on their native protein fold, which is difficult to preserve in a synthetic construct, they were not considered for MEV design^120^.

**Figure 3.**
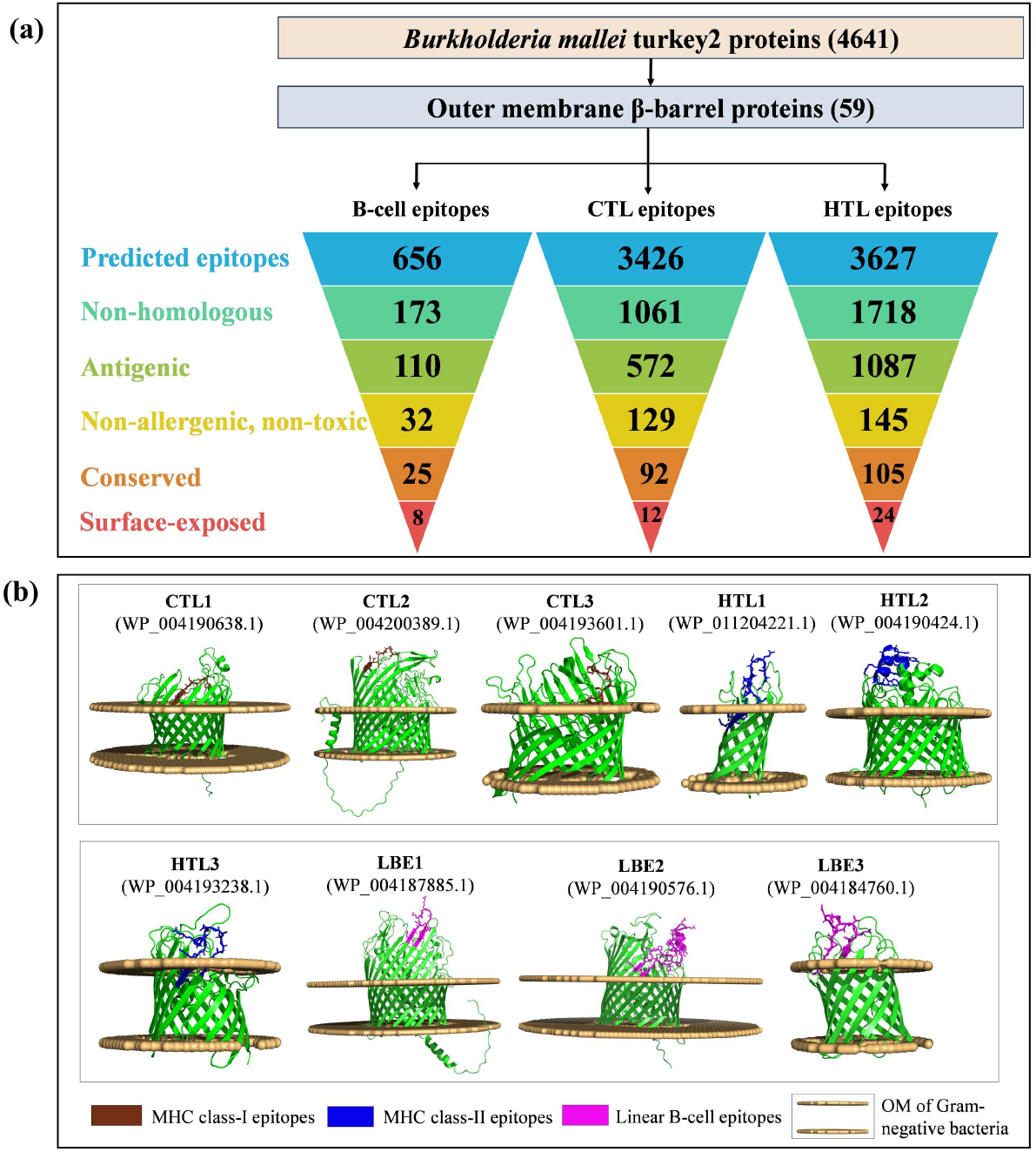
Predicted surface-exposed MHC class-I (CTL), MHC class-II (HTL), and linear B-cell epitopes (LBE). **(a)** Epitopes predicted from IEDB tools were filtered based on non-homology with human proteome, antigenicity, allergenicity, non-toxicity, conservancy, and surface accessibility. **(b)** Three surface-exposed epitopes with the highest prediction scores from each MHC class-I (magenta), MHC class-II (brown), and LBE (blue) were selected and mapped onto the structural models of their respective proteins embedded in bacterial outer membrane.

**Table 2:**
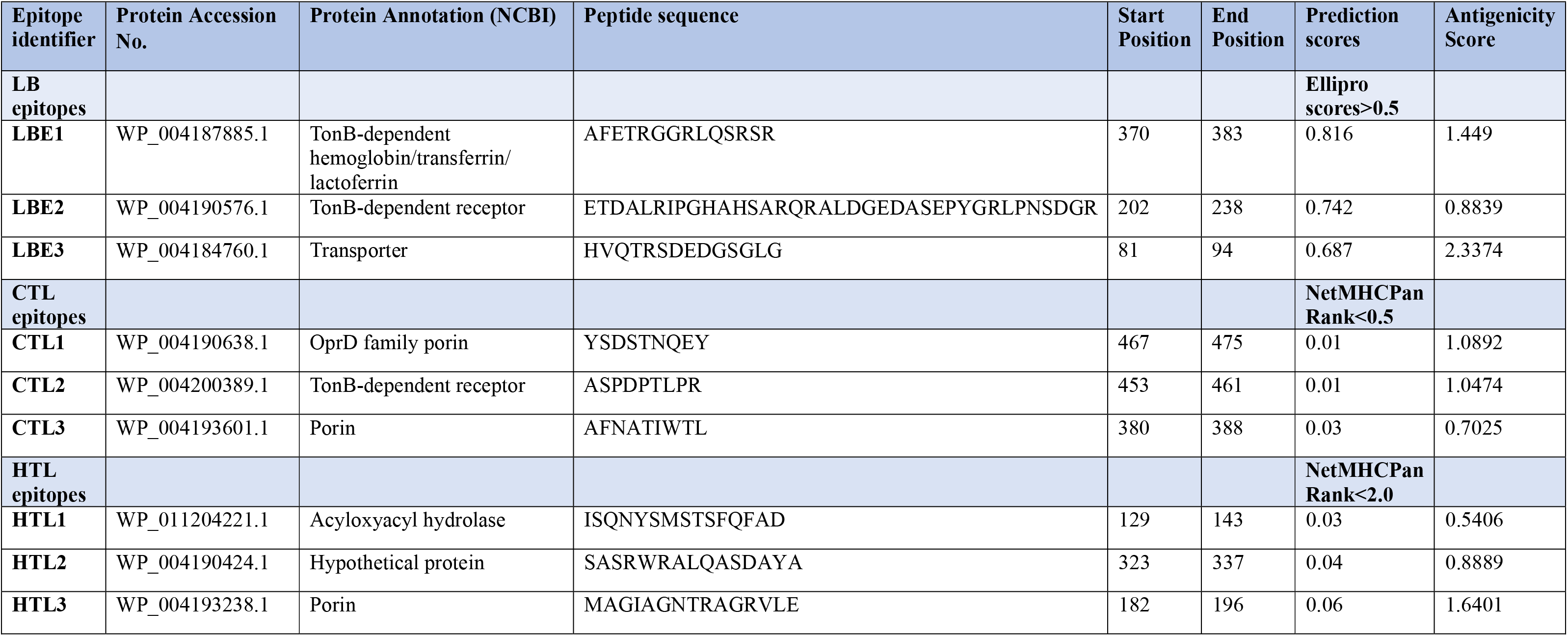
Selected surface-exposed Linear B-cell, CTL and HTL epitopes from the identified OMBBs for the construction of MEV.

### 3.4 Prediction and evaluation of T-cell epitopes

A total of 3426 CTL epitopes (9-mer) and 3627 HTL epitopes (15-mer) were predicted from 59 OMBB proteins using the IEDB MHC I (NetMHCIpan 4.1) and MHC II (NetMHCIIpan 4.0) prediction tools, respectively. Among these, 92 CTL and 105 HTL epitopes were identified as antigenic, non-allergenic, non-toxic, non-homologous to the human proteome, and conserved across 33 Bm strains. These epitopes were manually mapped onto the corresponding protein models that were embedded in a bacterial OM using PPM 3.0 server. Among these, 12 CTL and 24 HTL epitopes were located within the ECL region. Finally, three epitopes from each T-cell category were selected based on their NetMHCpan percentile score for inclusion into the MEV construct (Table 2). The selected six epitopes (CTL, and HTL) belonged to 6 OMBB proteins - WP_004190638.1, WP_004200389.1, WP_004193601.1, WP_011204221.1, WP_004190424.1, and WP_004193238.1 (Figure 3). Population coverage analysis demonstrated broad coverage of the selected CTL and HTL epitopes across the evaluated populations. The estimated global population coverage was 97.56%, with the highest in Austria (99.16%) and lowest in the United Kingdom (44.65%). Overall, the selected epitopes were suggested to provide broad HLA-mediated immune recognition across diverse populations (Figure S2).

### 3.5 Design of the MEV construct and evaluation of physicochemical properties

The proposed amino acid sequence of the MEV construct is ‘MIKLKFGVFFTVLLSSAYAHGTPQNITDLCAEYHNTQIYTLNDKIFSYTESLAGKREMAIITFKNGAIFQVEVPGSQHIDSQKKAIERMKDTLRIAYLTEAKVEKLCVWNNKTPHAIAAISMANEAAAKAKFVAAWTLKAAAAAYYSDSTNQEYAAYASPDPTLPRAAYAFNATIWTLHEYGAEALERAGISQNYSMSTSFQFADGPGPGSASRWRALQASDAYAGPGPGMAGIAGNTRAGRVLEKKAFETRGGRLQSRSRKKETDALRIPGHAHSARQRALDGEDASEPYGRLPNSDGRKKHVQTRSDEDGSGLGKKHHHHHH’ (Figure 4a).

**Figure 4.**
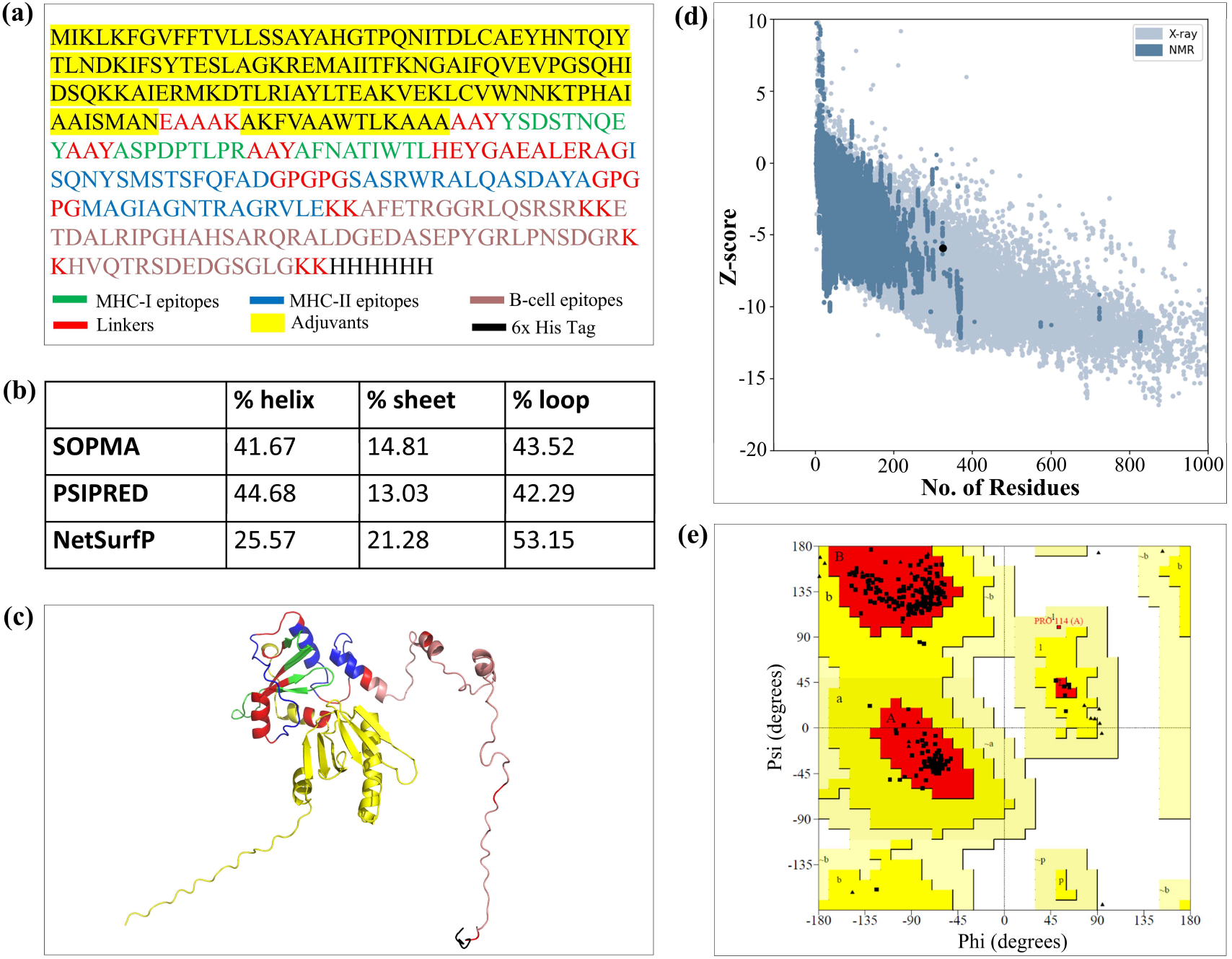
Structural analysis of the MEV construct. **(a)** The amino acid sequence of the MEV construct is shown consisting of adjuvants, linkers, 9 epitopes, and 6x His tag. **(b)** The assessment of MEV secondary structure using three computational tools. **(c)** The prediction of MEV tertiary structural model after refinement using GalaxyRefine server. (**d)** Z-score plot (−5.92) obtained from ProSA-web server. (**e)** Validation of MEV construct using PROCHECK. Ramachandran plot showing 96.8% of the amino acid residues falling in the most favored region.

CTL, HTL, and LBE were linked using flexible (GPGPG, AAY, and KK), and rigid (EAAAK) linkers. Flexible linkers are rich in Gly and Ser that enhance interdomain interactions, whereas rigid linkers maintain spacing and minimize unwanted interactions. CTL and HTL epitopes were fused via HEYGAEALERAG that enhance epitope presentation for the vaccine by creating a specific cleavage target for proteasomal and lysosomal degradation systems^121^. Adjuvants like CTB and PADRE peptides were added to enhance the immunogenicity. CTB is widely used for its ability to amplify immune responses^122^, and PADRE acts as a universal T-helper epitope that can induce robust, broad-spectrum MHC class-II responses and Th1 polarization^123^. The proposed MEV construct composed of 324 amino acid residues with molecular weight of 35.458 kDa. In the context of vaccine design, a molecular weight below 110 kDa is considered suitable for a vaccine construct^124^. Theoretical isoelectric point (pI) of the vaccine was calculated to be 9.35 indicating that the vaccine is basic in nature, which is acceptable, as most recombinant protein-based vaccines have pI values ranging from 5 to 10^125,126^. The predicted molecular formula of the MEV was C_1564_H_2433_N_459_O_471_S_8_. The instability index (II) was 36.17, classifying the MEV as a stable protein.

The hydrophilic nature of the protein was assessed based on the grand average of hydropathicity (GRAVY) score of −0.545, consistent with the inherent hydrophilic nature of the selected epitopes located within the ECL regions of OMBB proteins. In addition, MEV exhibited an antigenicity score of 0.7979 (greater than the threshold value of 0.4) and was found to be non-allergenic and non-toxic. NetSolP predicted a solubility score of 0.6212, suggesting that the protein remains soluble upon overexpression (Table 3). An OmpA family protein (Omp7) that was found to be immunogenic in mice models and considered as a potential vaccine candidate against *B. pseudomallei*^29^ was taken as positive control. Omp7 was also found to be stable, hydrophilic, non-allergenic, and non-toxic; however, it exhibited lower antigenicity (0.5146) but higher solubility (0.8624) compared to the MEV construct (Table 3).

**Table 3:** Physicochemical properties of the MEV construct and Omp7.

| Physicochemical Properties | Tool Prediction |  |
| --- | --- | --- |
|  | MEV construct | Omp7<sup>‡</sup> |
| Amino acid length | 324 residues | 177 residues |
| Molecular weight | 35.458 kDa | 19.608 kDa |
| Theoretical pI | 9.35 | 7.88 |
| Instability index (II) | 36.17 | 38.93 |
| GRAVY score | -0.545 | -0.503 |
| Antigenicity score | 0.7979 | 0.5146 |
| Allergenicity | Non-allergenic | Non-allergenic |
| Toxicity | Non-toxic | Non-toxic |
| Solubility | 0.6212 | 0.8624 |

<sup>‡</sup>Omp7 is an established vaccine candidate in *B. pseudomallei* that was taken as positive control.

### 3.6 Prediction of secondary and tertiary structure of the MEV construct

The secondary structure of MEV was predicted to have higher percentage of flexible loops followed by α-helixes and β-extended strands as depicted in Figures 4b, S3. The high proportion of flexible loops in the MEV suggests an enhanced probability of antigenic epitope formation^127^. Additionally, residues from 251-324 in the MEV were predicted to form an IDR as they are derived from flexible, surface exposed epitopes of the OMBB proteins. Even though the MEV construct lacks well-defined secondary structure, it remains highly soluble upon overexpression as it is composed of extracellular loop regions, which are inherently hydrophilic^128^. The tertiary structure of the MEV was modeled using four tools (Figure S4). As the MEV comprises epitopes derived from intrinsically flexible ECL regions and lacks a suitable experimental structural template, the predicted MEV structures exhibited a relatively low confidence scores- I-TASSER (TM score = 0.3, C-score = −3.7), RoseTTAFold2 (pLDDT = 0.48), ESMfold (pTM = 0.23), and AlphaFold 3 (pTM = 0.39, pLDDT = 55.23). These tertiary structures were optimized using GalaxyRefine server. We selected the best-refined model based on the highest percentage of Rama-favored residues, lowest clash-score, and highest GDT-HA (Global Distance Test - High Accuracy) score (Table S7).

AlphaFold 3 model exhibited highest proportion of Rama-favored residues compared to other models. Therefore, it was subjected to a second round of refinement and selected for further analysis. The tertiary structure of the MEV was visualized using PyMOL, where surface-exposed LBE, HTL, and CTL epitopes were mapped onto the tertiary structure, as shown in Figure 4c.

### 3.7 Model quality assessment of MEV construct

The quality of the selected MEV model was assessed using ProSA-web server, which yielded a Z-score of −5.92 (Figure 4d). To assess the accuracy of the predicted structure, we further evaluated the quality of the model using PROCHECK that generated a Ramachandran plot (Figure 4e).

Analysis showed that 96.8% of the amino acid residues were in most favoured regions, 3.2% in additionally allowed region, and no residue in disallowed region, confirming the structural reliability. Overall, the predicted tertiary structure of the MEV suggested a good vaccine design.

### 3.8 Molecular docking of the MEV construct with human TLR4 and TLR2 immune receptor

Molecular docking was performed to predict potential interactions between the vaccine construct (ligand molecule) and human immune receptors - TLR4 and TLR2, based on energy minimization and structural complementarity at the receptor’s active site. Several docking clusters were generated, and the top-ranked cluster was selected for analysis. For the MEV-TLR4 complex, the best docked complex exhibited a docking score of −286.34, and a confidence score of 0.9. The 3D structure of the docked complex was visualized in PyMOL (Figure 5). PRODIGY server predicted a negative ΔG value of −12.7 kcal mol^-1^ and Kd value of 4.9 × 10⁻^10^ M. For MEV-TLR2 complex, the best docking score was −309.08 with a confidence score of 0.9. It also exhibited a negative ΔG value of −18.2 kcal mol^-1^ and Kd value of 4.7 × 10⁻^14^ M. A docking score below −200 indicated strong binding interactions which were in concordance with Omp7-TLR4 (−13.2 kcal mol^-1^), and Omp7-TLR2 (−11.9 kcal mol^-1^). Negative ΔG and low Kd values revealed strong binding affinity of MEV for both TLR4 and TLR2. These results were comparable to the Omp7-TLR4 (ΔG: −13.2 kcal mol^-1^, Kd value: 2.2 × 10⁻^10^ M), and Omp7-TLR2 (ΔG: −11.9 kcal mol^-1^, Kd value: 1.9 × 10⁻^9^ M) docked complexes (Figure 5a).

**Figure 5.**
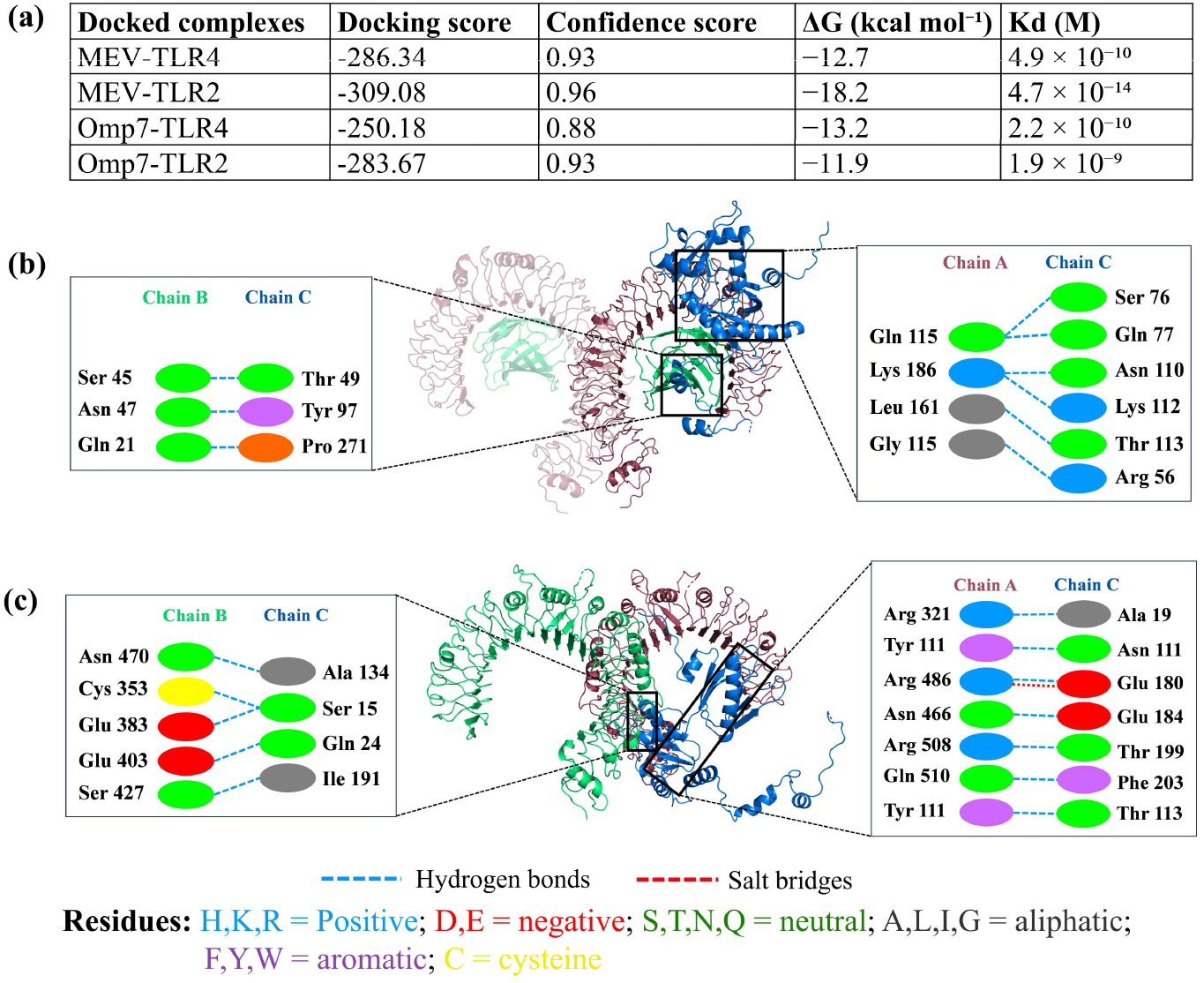
Interaction analysis of the docked MEV construct with human TLR4 and TLR2. **(a)** Binding energetics of docked MEV- and Omp7 - TLR complexes. **(b)** The docked complex of the MEV and TLR4 is shown. The binding interactions between the complex predicted formation of nine hydrogen bonds. **(c)** MEV-TLR2 complex formed 18 hydrogen bonds and one salt bridge at protein-protein interface.

Further, the binding interactions analyzed using PDBePISA revealed that the MEV forms nine hydrogen bonds with both TLR4 and its coreceptor MD-2 (Figure 5b). The MEV construct also interact with TLR2 via 12 hydrogen bonds and one salt bridge within the range of 5 Å interatomic distance (Figure 5c). Similar interactions were also observed in positive control where Omp7 formed 11 hydrogen bonds and one salt bridge with TLR4, and ten hydrogen bonds and three salt bridges with TLR2 (Figure S6).

### 3.9 Protein flexibility analysis and NMA of MEV-TLR4/TLR2 complexes

CABS-flex predicted RMSF values for each residue in the MEV-TLR4/TLR2 complexes over a time period equivalent to 10ns simulation. Higher RMSF values reflect greater flexibility, while lower values indicate restricted motion of the system. In MEV-TLR4 complex, RMSF values ranged from 0.1 to 5.1 (Figure 6a), whereas the MEV-TLR2 complex exhibited a broader range of 0.1 to 10.73 Å (Figure 6b). Most residues in the MEV-TLR4/TLR2 complexes displayed low RMSF values, supporting overall structural stability. However, localized increases in flexibility at the MEV-TLR2 docking interface suggest induced-fit conformational adjustments. Moreover, in MEV-TLR4 complex, the RMSD values for medoids (representative structure of each cluster) ranged from 3.7 Å - 4.2 Å compared to input reference structure. In the MEV-TLR2 complex, the corresponding RMSD values ranged from 5.5 to 6.9 Å. These relatively narrow RMSD ranges indicate that the complexes remained structurally stable throughout the simulation. The positive control also showed comparable RMSF profiles, with values ranging from 0.09 to 8.71 Å for the Omp7-TLR4 complex and 0.05 to 6.76 Å for the Omp7-TLR2 complex, indicating overall structural stability with localized flexible regions (Figure S7). These findings are consistent with the MEV-TLR4/TLR2 complexes that supports stable conformational behavior and residue-level fluctuations.

**Figure 6.**
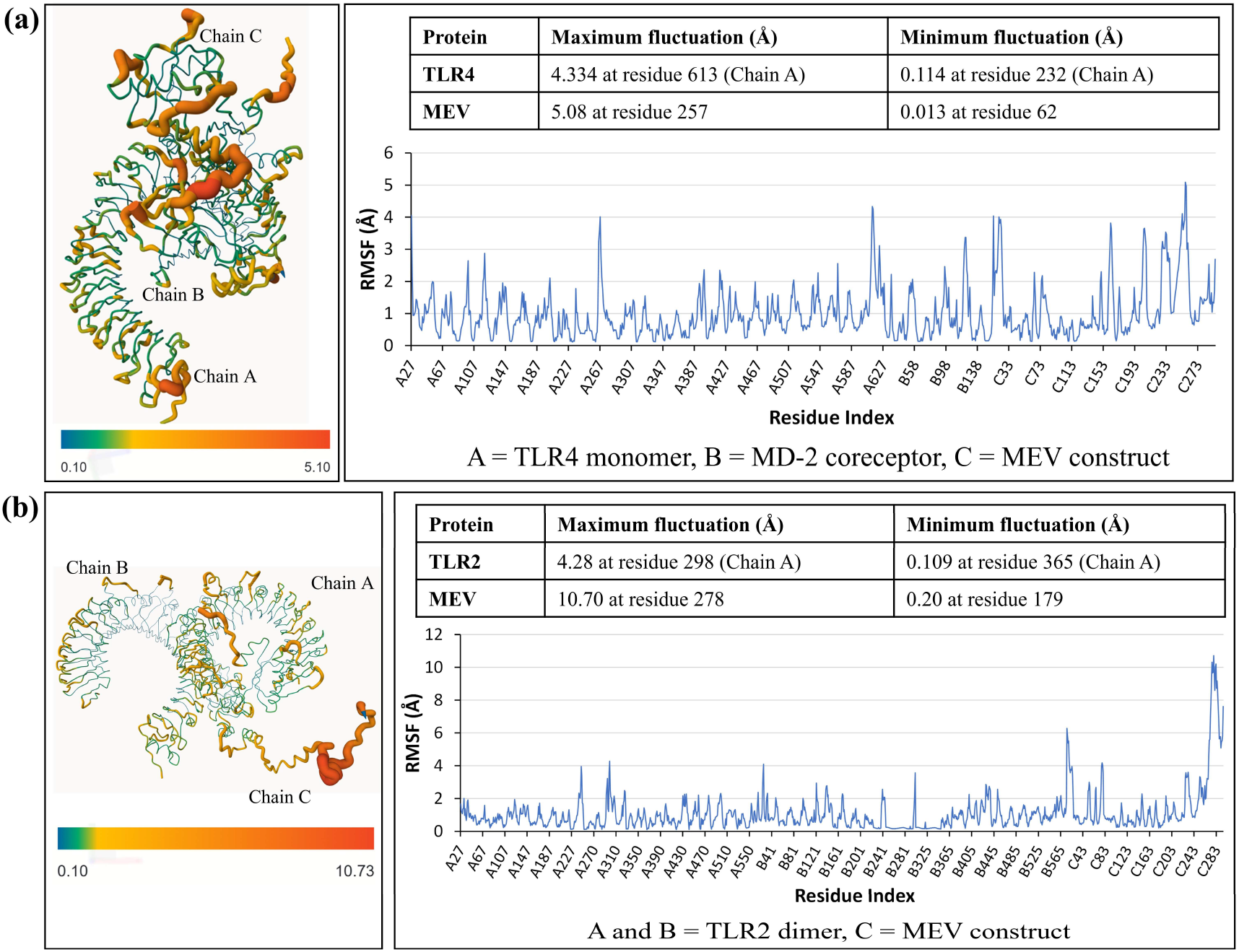
Results of RMSF analysis using CABS-flex. **(a)** RMSF-based model and graph showing flexibility distribution in the MEV–TLR4 complex. **(b)** RMSF profile of MEV–TLR2 complex throughout the simulation.

NMA was performed for further analysis to describe the flexible states accessible to a protein around an equilibrium position^129^. iMODS was used to perform NMA-based simulation, concentrating on low-frequency modes that depict large-scale, physiologically significant movements. Four output plots - eigenvalue spectra, B-factor comparison, covariance map, and elastic network model - were evaluated for both the complexes (Figures 7, S5). The eigenvalue for MEV-TLR4/TLR2 complexes (∼9.2 × 10⁻⁶ and ∼1.2 × 10⁻⁶) indicated high resistance to large-scale deformation responsible for mechanical stability (Figures 7a, 7e). The MEV-immune receptor complexes exhibited lower eigenvalues than the corresponding positive control complexes (Omp7-TLR4 = ∼1.9 × 10⁻^5^ and Omp7-TLR2 = ∼1.7 × 10⁻^5^) that indicates limited fluctuation of the MEV-receptor complexes (Figure S8). The B-factor profile from NMA showed relatively low B-factor fluctuations across most regions of both the complexes that indicates overall structural rigidity.

**Figure 7.**
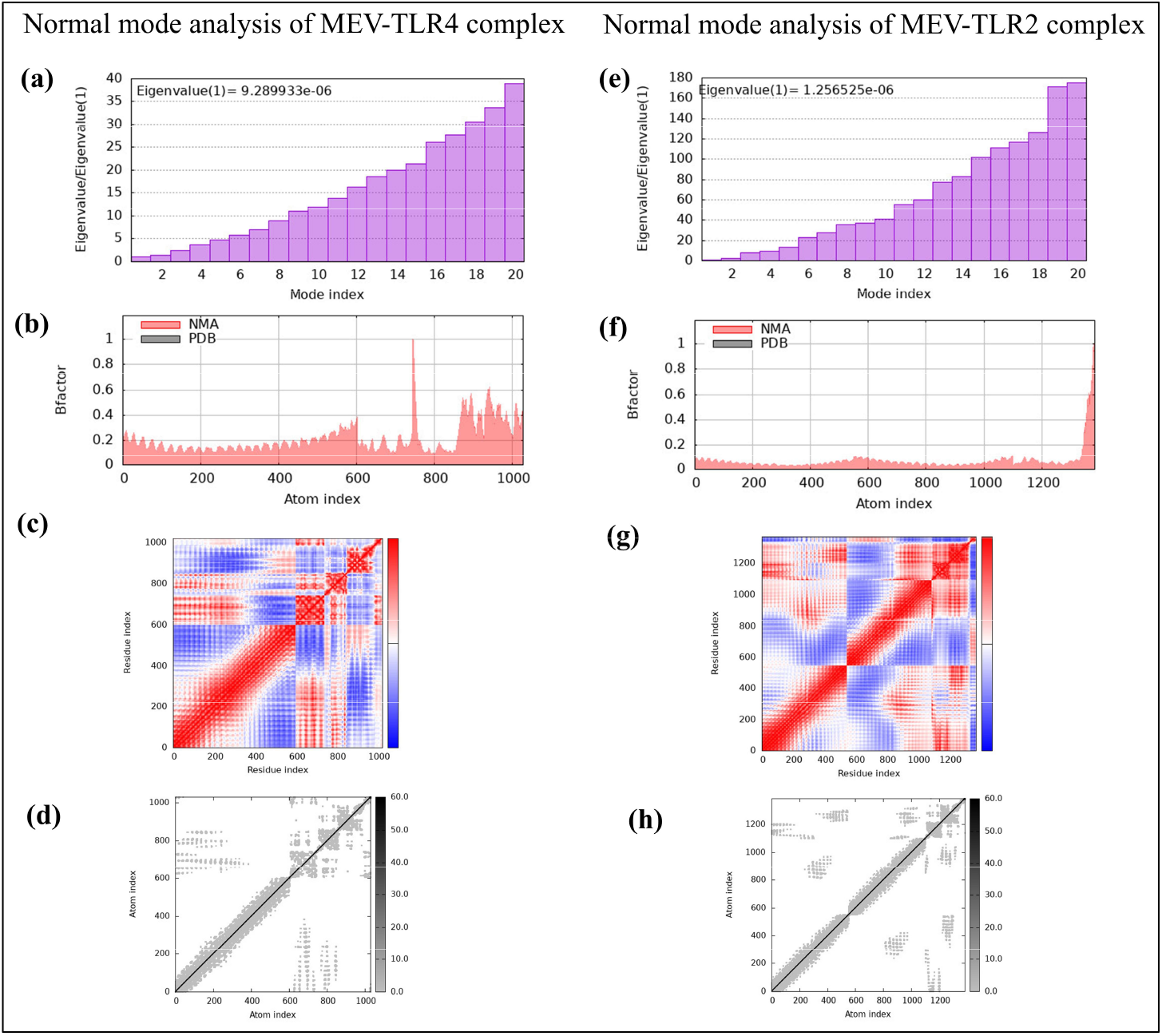
**Results of NMA of the MEV-TLR4/TLR2 complexes**. **(a, e)** Eigenvalue profiles of the first 20 normal modes. **(b, f)** NMA-derived and experimental PDB B-factor profiles. **(c, g)** Correlation matrices illustrating correlated and anticorrelated residue motions. **(d, h)** Elastic network representations showing residue-wise dynamic coupling in the MEV-TLR4 and MEV-TLR2 complexes, respectively.

High B-factor values near the binding interface in MEV-TLR4, and at C-terminal in MEV-TLR2 represents adaptive flexibility without destabilization (Figures 7b, 7f). The covariance map displayed a matrix where red regions in the matrix illustrate a decent correlation between residues, the white regions indicate uncorrelated motion, and the blue regions depict anti-correlated motion. The greater correlations indicate better functional stability. The complexes showed strong correlated motions along the diagonal, confirming coherent domain movement and structural stability (Figures 7c, 7g). The elastic network model showed dense interatomic contacts, reflecting a mechanically robust complex with global structural integrity (Figures 7d, 7h). Overall, the B-factor, covariance, and elastic network profiles of the MEV- immune receptor complexes were comparable to that of positive control (Figure S8).

### 3.10 MD simulation of MEV-TLR4/TLR2 complexes

The RMSD profiles of the MEV-TLR4/TLR2 complexes were analyzed over a period of 100 ns MD simulation. In the MEV-TLR4 complex, the TLR4 backbone RMSD initially increased from ∼1.2 Å to 4.0 Å during the first 25 ns, followed by fluctuations ranging from 3.0 Å to 4.5 Å for the remaining simulation. The MEV RMSD increased rapidly during the initial phase and subsequently fluctuated between ∼10 Å to 17 Å, with higher deviations at 35-55 ns and 80-90 ns, followed by decrease in RMSD by the end of the simulation. These larger fluctuations are consistent with the conformational flexibility of the MEV construct and IDR, rather than dissociation from TLR4 (Figure 8a). These results were comparable with positive control where RMSD values of TLR4 and Omp7 ranged from ∼1.15 Å to 4.6 Å and ∼2.16 Å to 9.5 Å, respectively (Figure S9a). For the MEV-TLR2 complex, the TLR2 RMSD increased rapidly from ∼2.5 Å to ∼3.5 Å within the first few ns and remained relatively stable throughout the simulation. The MEV RMSD increased during the initial phase and then remained comparatively stable around 12-14 Å (Figure 8b). In case of Omp7-TLR2 complex, the RMSD value for TLR2 fluctuated broadly ranging from ∼1.9 Å to 7.0 Å, while Omp7 remained stable between RMSD values of ∼1.7 Å to 4.7 Å (Figure S9b). The maintenance of relatively stable plateaus suggests that both MEV-TLR4 and MEV-TLR2 complexes retained their overall structural integrity during the simulations, and also exhibited RMSD profile comparable to Omp7-immune receptor complexes.

**Figure 8.**
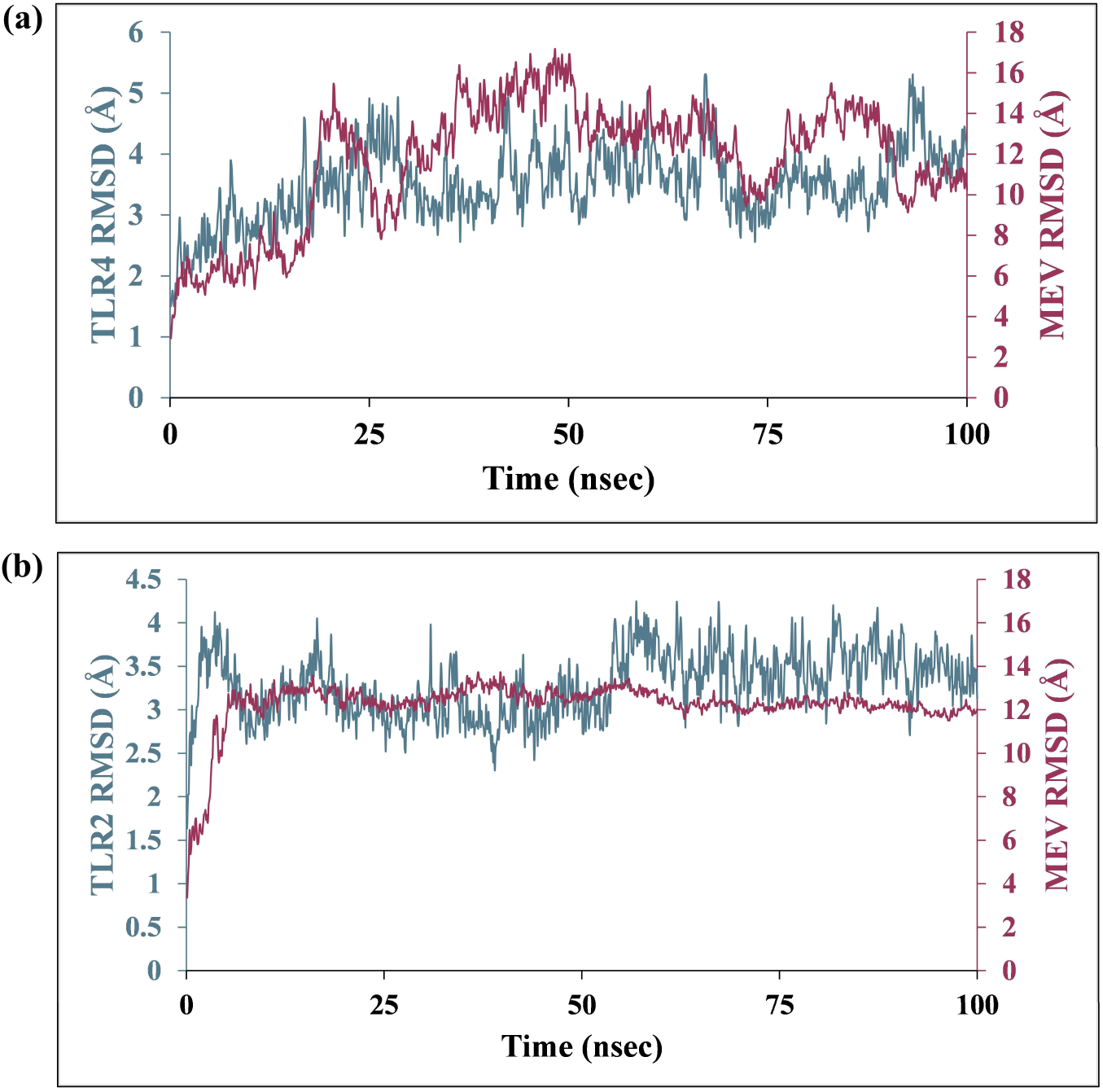
RMSD profiles of MEV-TLR4/TLR2 complexes over 100 ns MD simulation. **(a)** RMSD profile of TLR4/MD-2 and the MEV construct. **(b)** RMSD profile of TLR2 dimer and the MEV construct.

### 3.11 Immune simulation of the MEV construct

Immune simulation was performed to assess the immunological response of the human body to the MEV construct (Figure 9). Following the primary dose of the MEV, B-cell population increased upto ∼145 cells/mm³. Increase in memory B-cells over time suggests long-term immune memory (Figure 9a). There was also an increase in the population of HTLs (∼2,100 cells/mm³), while CTLs remained fluctuating around the baseline during vaccination (Figures 9b, and 9c). Increase in CD4⁺ T helper (TH) cells further stimulate B-cells and T cytotoxic (TC) cells. IgM levels rose initially but later declined due to class-switching to IgG1 and IgG2, indicating strong adaptive immunity (Figure 9d). During subsequent exposures, the levels of IgM + IgG, IgG1 + IgG2, IgG1, and IgG2 were markedly higher than in the primary response, reflecting a strong secondary immune response. Furthermore, cytokines IFN-γ and IL-2 were produced at substantially higher concentrations as compared to other cytokines that suggests TH1 mediated immunity (Figure 9e). Further, TH1 cells reached nearly 100% of the TH cell population post vaccination, indicating a TH1-biased immune profile, suitable for combating intracellular pathogens such as *Burkholderia* (Figure 9f). Immune simulation of OMP7 revealed a comparable immune response, with several notable differences from MEV (Figure S10). The B-cell population was slightly lower (∼135 cells/mm³) compared to the MEV. In contrast, the HTL population was marginally higher (∼2,300 cells/mm³); however, the IgM + IgG antibody response exhibited lower peak levels in case of Omp7. Overall, the MEV elicited a stronger immune response with higher antibody titres compared to Omp7.

**Figure 9.**
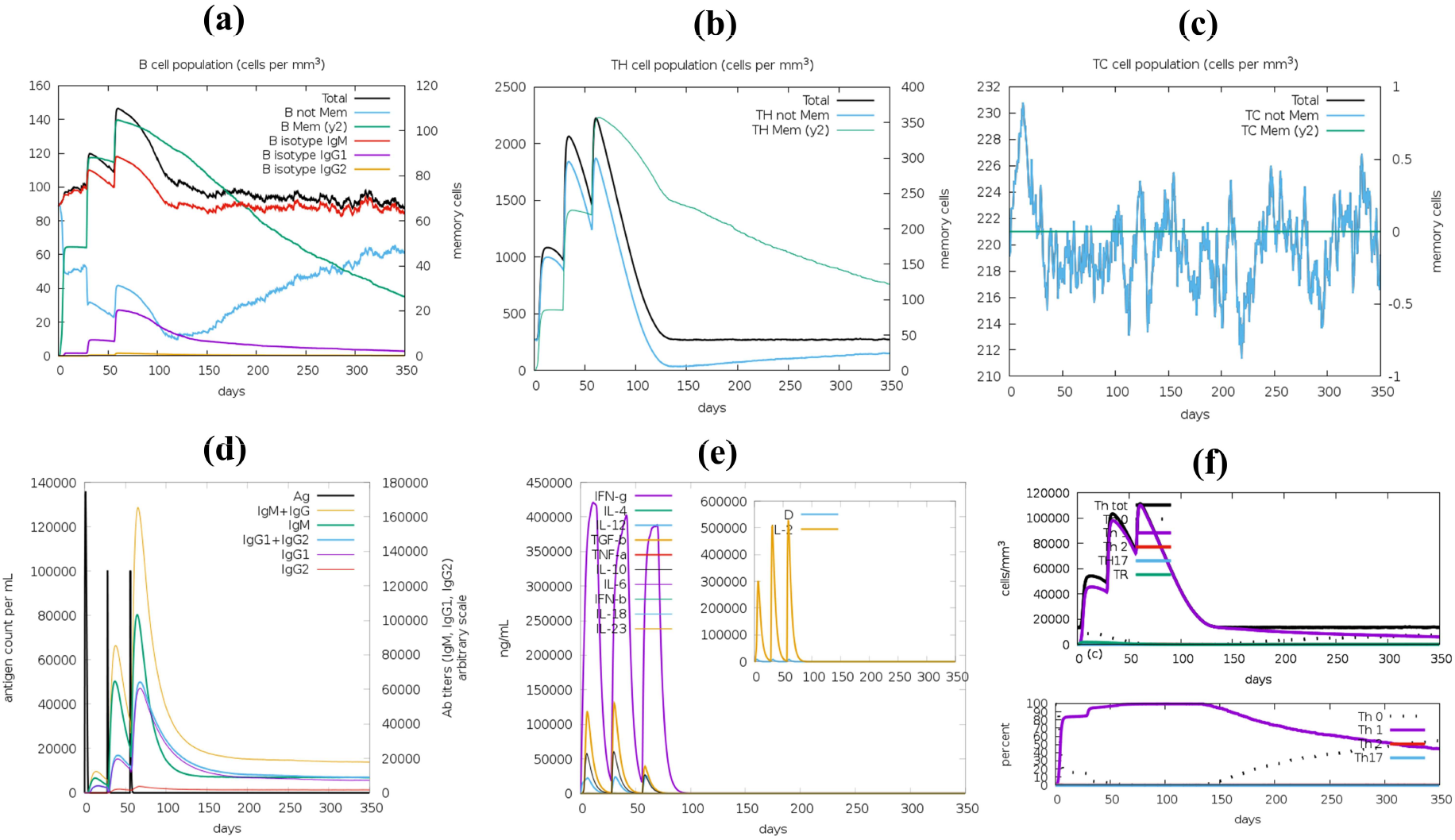
In-silico immune simulation of MEV construct using C-ImmSim server. **(a)** B-cell isotypes in various states. **(b)** Helper T-cell population. **(c)** Cytotoxic T-cell population. **(d)** Antigen and subtypes of immunoglobulin levels. **(e)** Concentration of cytokines and interleukins at three different stages (8 hrs, 28 days, and 56 days). **(f)** Helper T-cell isotypes in various states.

### 3.12 Codon optimization and in-silico cloning using Benchling

Sequence of the Bm MEV gene was optimized to improve its expression in the *E. coli* K-12 expression system. Size of MEV gene sequence was 972 bp with CAI of the optimized sequence estimated to be 0.94. This indicated a high level of compatibility with the codon usage bias of the host organism. The GC content was 52.36% falling within the optimal range (40-60%) for efficient transcription and translation. The optimized gene was subsequently cloned in-silico into the pET-28a(+) expression vector using Benchling. The recombinant construct exhibited the MEV gene sequence downstream of the T7 promoter with the C-terminal 6X His-tag exposed for purification using affinity chromatography (Figure 10).

**Figure 10.**
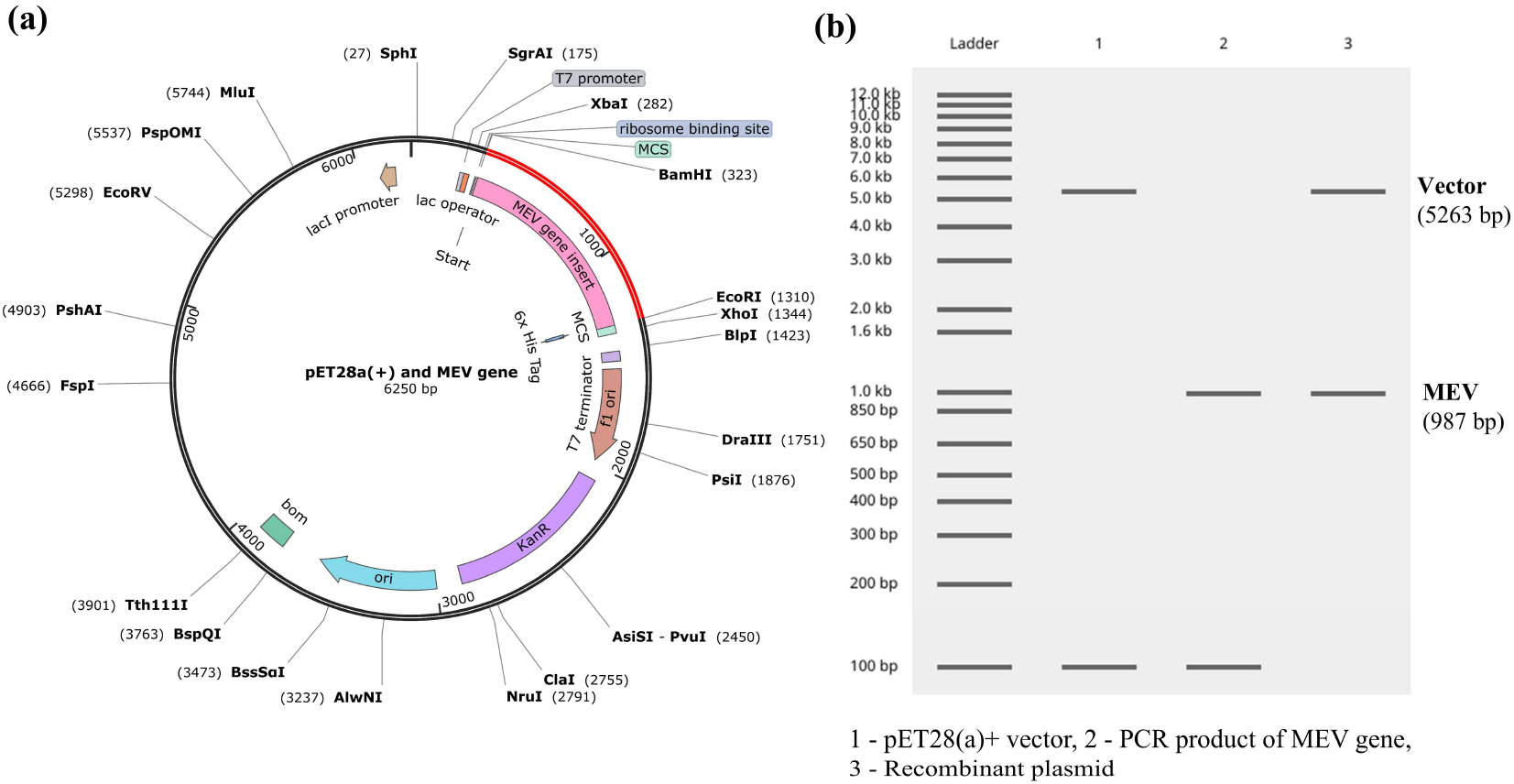
In-silico cloning of the codon-optimized MEV construct into the pET-28a(+) expression vector. **(a)** The optimized MEV sequence (red) was inserted into the MCS of the pET-28a(+) vector downstream of the T7 promoter. The recombinant plasmid contains a C-terminal 6×His tag for affinity purification and kanamycin resistance for selection. **(b)** The virtual restriction digestion analysis shows the expected DNA fragments, confirming the successful in-silico insertion of the vaccine construct into the expression vector.

## 4. Discussion

Glanders is one of the oldest known zoonotic diseases, described by Aristotle in 384-322 BC, caused by Bm^130^. Bm infection can cause acute septicaemia and pulmonary complications that can be fatal without treatment. Bm attracts attention as a potential biological warfare agent due to multidrug resistance, rapid transmission via aerosol, and absence of an approved vaccine against the bacteria^131^. This study systematically identified and characterized OMBB proteins in Bm and utilized them to design an MEV construct. The proteome wide approach divided all the proteins of Bm turkey2 into five groups (A-E) based on the number of computational tools that predicted them as β-barrels. Since Groups A and B proteins received positive predictions from a greater number computational tools, they were considered for further analyses to ensure a high level of confidence in the β-barrel predictions. Groups C-E exhibited lower consensus among prediction tools that limits the confidence of their classification as β-barrels. Therefore, these proteins were excluded from the present study but may represent potential OMBB repertoire that requires further investigation in future studies. Groups A and B consisted of 59 putative OMBB proteins belonging to functionally diverse categories such as porins, TonB-dependent receptors, efflux proteins, secretion system components, adhesins, and autotransporters. Structural validation using five modeling tools supported the prediction of β-barrel architecture, where structural alignment showed strong concordance with RMSD values under 5 Å, indicating minimal structural deviation and high model accuracy. These proteins comprised diverse β-barrel architectures ranging from 8 to 26 β-strands. Literature survey showed that most of the identified OMBB proteins had homologs in other *Burkholderia* species and were annotated in NCBI. The identification of previously characterized β-barrel proteins further supported the reliability of our prediction pipeline. Among the identified OMBB proteins were well-known proteins such as BamA, LptD, and TamA, that play essential roles in OM biogenesis. Bm also possessed a substantial number of porins, TonB-dependent receptors, and TolC homologs that represents the functional diversity of its OMBB repertoire. Notably, two novel OMBB proteins were identified in this study, one predicted to contain a double β-barrel architecture and the other was an 18-stranded β-barrel. Sequence variation analysis revealed variations in OMBB proteins across 33 Bm strains in ECL regions that suggested ongoing host-driven selection pressure.

Since OMBB proteins are in direct contact with the host cells, they were leveraged to identify antigenic, non-allergenic, non-toxic, conserved, and surface-exposed B-cell and T-cell epitopes which were non-homologous to human proteome with high population coverage in glanders-endemic regions. These epitopes belonged to 9 out of 59 OMBB proteins (Table 2, Figure 3). The selected epitopes were assembled into an MEV construct using multiple linkers, and adjuvants to enhance the stability, potency, and durability of the vaccine construct^132^. The MEV construct revealed favorable physicochemical properties, high antigenicity, and good solubility in *E. coli* for efficient heterologous expression comparable to the immunogenic protein Omp7 taken as positive control. Since receptor binding is as important as structural stability in a vaccine design, molecular docking of MEV was performed with human immune receptors. This demonstrated strong binding affinity of the MEV with human TLR4 and TLR2 showing multiple hydrogen bonds and salt bridges. TLR4 and TLR2 were mostly interacting with CTB adjuvant as it is structurally well-characterized, whereas vaccine epitopes are derived from loop regions that have greater conformational variability. This suggests that the adjuvant may contribute to TLR-mediated innate immune activation, while the epitopes can subsequently undergo antigen processing and presentation to induce immune responses. Furthermore, per-residue RMSF profiles, RMSD values over 100 ns timescale, and NMA, collectively confirmed the structural stability and adaptive flexibility of the docked complexes. These analyses were also in concordance with the immunogenic protein Omp7, thereby increasing confidence in the structural stability and reliability of the proposed MEV construct. Immune simulations also predicted a sustained immune response characterized by strong B- and T-cell activation, memory formation, and a TH1-biased cytokine profile. The in-silico cloning of the optimized vaccine construct into pET-28a(+) vector will facilitate its future cloning, recombinant protein expression, and experimental evaluation.

From a broader vaccine development perspective against Bm, epitope-based vaccines are safer alternatives to live attenuated vaccines^133^. A previous study proposed an MEV based on antigens derived from only two proteins with limited number of antigens that were not exclusively surface-exposed^34^. In contrast, this study integrates multiple OMBB-derived surface exposed epitopes to enhance a diverse immune response. The identification of conserved epitopes across 33 Bm strains facilitates cross-strain protection, while epitopes non-homologous to humans minimize the risk of unintended cross-reactivity with human proteins. Despite its promise, this study is limited by its purely in-silico approach which may not fully represent its biological complexity. Ultimately, this study lays the foundation for future work that requires experimental validation of the novel OMBB proteins, and the MEV construct through in vitro expression, purification, and immunogenicity testing in animal models, followed by challenge studies to assess protective efficacy.

## 5. Conclusion

This study presents a curative OMBB repertoire of Bm that was utilized to develop a potential MEV construct against human glanders. The computational approach identified diverse OMBB proteins including two novel OMBB proteins. These proteins were leveraged to predict surface-exposed epitopes for the development of a promising MEV candidate. The resulting MEV construct exhibited favourable physicochemical properties, stability, solubility, and high antigenicity, along with reliable structural quality. Molecular docking and MD simulations revealed stable interaction of MEV with immune receptors. Although these in-silico findings strongly support MEV’s potential, experimental validation through in vitro and in vivo studies will be essential to confirm its immunogenicity, safety, and protective efficacy.

## Supporting information

Supplementary material revised

## Acknowledgements

JK is a recipient of junior research fellowship from the University Grants Commission (Reference number: 231620067077), Government of India. AP is a recipient of senior research fellowship from the Department of Biotechnology (Grant number: DBT/2023-24/UOD/2326), Government of India.

## Author contributions statement

**Jahnvi Kapoor:** methodology, software, investigation, data analysis, data curation, writing-original draft preparation. **Amisha Panda:** data analysis, visualization, writing-reviewing and editing. **Sanjiv Kumar:** conceptualization, methodology, software, writing-reviewing and editing, supervision. **Anannya Bandyopadhyay**: methodology, data analysis, writing-original draft preparation, writing-reviewing and editing, supervision. All authors reviewed the final draft of the manuscript. All authors contributed to the manuscript revision, read and approved the submitted version.

## Conflict of interest

The authors declare no competing financial interest.

## Funding

This research received no specific grant from any funding agency in the public, commercial, or not-for-profit sectors.

## Data Availability Statement

The data supporting the findings of this study has been deposited in the Zenodo public repository accessible at https://zenodo.org/records/22742445

## Ethical Approval

Ethics approval statement is not applicable.

## Notes

### Competing Interest Statement

The authors have declared no competing interest.

### Summary of Updates

The revised submission emphasizes on the importance of outer membrane β-barrel (OMBB) proteins in Burkholderia mallei as potential targets for the development of a multi-epitope vaccine. A new MEV construct has been proposed, supported by additional evidences and comprehensive in silico analyses, which strengthen the design, and predicted immunogenicity of the vaccine candidate.

