## Supplementary material revised for "Targeting Outer Membrane β-barrel Proteins of *Burkholderia mallei* to design a Multi-epitope Vaccine Against Human Glanders"

**Supplementary Tables**

**Table S1:** Comprehensive results from computational tools employed in our study to predict outer membrane β-barrel proteins in *B. mallei*

| **Protein Accession No.** | **Protein Annotation (NCBI)** | **Protein Annotation (UniProt)** | **OMPdb Match** | **TMBETADISC** | **MCMBB Score** | **TMbed** | | **DeepTMHMM Prediction** | **Pepstats** | | | | **PSORTb** | | **SignalP** | | **SPAAN** | **CELLO** | |
| --- | --- | --- | --- | --- | --- | --- | --- | --- | --- | --- | --- | --- | --- | --- | --- | --- | --- | --- | --- |
|  |  |  |  |  |  | **tmbed_b** | **tmbed_B** |  | **AA Length** | **MolWt**  **(Da)** | **Charge** | **pI** | **Localization** | **psortb** | **Signal Peptide** | **CS Position** | **P_ad_ value** | **Localization** | **Score** |
| **Group A** |  |  |  |  |  |  |  |  |  |  |  |  |  |  |  |  |  |  |  |
| WP_004189302.1 | carbohydrate porin | Carbohydrate porin, OprB family | 500 | Outer Membrane Protein | 0.044 | 73 | 69 | BETA | 504 | 53079.42 | 1.5 | 6.9943 | OuterMembrane | 9.93 | SP(Sec/SPI) | CS pos: 33-34. ALA-QA. Pr: 0.4756 | 0.779522 | OuterMembrane | 2.604 |
| WP_004190638.1 | OprD family porin | Outer membrane porin, OprD family | 500 | Outer Membrane Protein | 0.022 | 81 | 76 | BETA | 486 | 52353.6 | 6.5 | 7.4483 | OuterMembrane | 9.93 | SP(Sec/SPI) | CS pos: 34-35. ARA-DD. Pr: 0.6924 | 0.781116 | OuterMembrane | 2.174 |
| WP_004199949.1 | OmpW/AlkL family protein | Outer membrane protein | 500 | Outer Membrane Protein | 0.037 | 39 | 36 | BETA | 243 | 25899.34 | 6 | 9.3622 | OuterMembrane | 9.93 | SP(Sec/SPI) | CS pos: 24-25. AHA-QS. Pr: 0.9025 | 0.811195 | Extracellular | 3.249 |
| WP_004184655.1 | porin Omp38 | Outer membrane porin OpcP | 500 | Outer Membrane Protein | 0.101 | 70 | 65 | BETA | 376 | 39396.13 | 9 | 9.5937 | OuterMembrane | 10 | SP(Sec/SPI) | CS pos: 20-21. AHA-QS. Pr: 0.9575 | 0.928625 | OuterMembrane | 2.681 |
| WP_004188615.1 | porin | Putative outer membrane porin | 500 | Outer Membrane Protein | 0.08 | 69 | 64 | BETA | 384 | 40221.47 | 4.5 | 8.7361 | OuterMembrane | 10 | SP(Sec/SPI) | CS pos: 22-23. AQA-QS. Pr: 0.9152 | 0.833356 | OuterMembrane | 3.496 |
| WP_004206816.1 | porin | Putative outer membrane porin | 500 | Outer Membrane Protein | 0.093 | 67 | 65 | BETA | 390 | 40632.81 | 5.5 | 8.774 | OuterMembrane | 10 | SP(Sec/SPI) | CS pos: 24-25. VHA-QS. Pr: 0.9664 | 0.872446 | OuterMembrane | 3.922 |
| WP_053799657.1 | porin | Porin | 500 | Outer Membrane Protein | 0.035 | 68 | 66 | BETA | 359 | 37935.5 | 7.5 | 8.7084 | OuterMembrane | 10 | LIPO(Sec/SPII) | CS pos: 18-19. AGA-CA. Pr: 0.9233 | 0.768831 | OuterMembrane | 2.538 |
| WP_004191758.1 | porin | Outer membrane porin, putative | 500 | Outer Membrane Protein | 0.055 | 68 | 65 | BETA | 362 | 38364.4 | 5.5 | 9.2182 | OuterMembrane | 9.95 | SP(Sec/SPI) | CS pos: 19-20. AHA-QS. Pr: 0.9517 | 0.945994 | Extracellular | 3.126 |
| WP_004195141.1 | porin | Putative outer membrane porin | 500 | Outer Membrane Protein | 0.084 | 69 | 63 | BETA | 386 | 40187.58 | 6.5 | 9.0575 | OuterMembrane | 10 | SP(Sec/SPI) | CS pos: 20-21. AHA-QS. Pr: 0.9677 | 0.800629 | OuterMembrane | 3.392 |
| WP_004197133.1 | porin | Porin | 500 | Outer Membrane Protein | 0.062 | 68 | 66 | BETA | 366 | 38631.23 | 5 | 9.0827 | OuterMembrane | 10 | SP(Sec/SPI) | CS pos: 19-20. AHA-SD. Pr: 0.9964 | 0.824755 | OuterMembrane | 4.563 |
| WP_004197145.1 | porin | Outer membrane porin | 500 | Outer Membrane Protein | 0.036 | 71 | 67 | BETA | 386 | 41575.04 | 3.5 | 7.2286 | OuterMembrane | 9.93 | TAT(Tat/SPI) | CS pos: 33-34. AHA-QS. Pr: 0.8995 | 0.822312 | OuterMembrane | 3.242 |
| WP_004198441.1 | porin | Putative outer membrane porin | 500 | Outer Membrane Protein | 0.072 | 69 | 65 | BETA | 384 | 40678.27 | 8.5 | 9.4769 | OuterMembrane | 10 | SP(Sec/SPI) | CS pos: 20-21. VFA-QS. Pr: 0.9595 | 0.709991 | OuterMembrane | 4.413 |
| WP_004199671.1 | porin | Porin | 500 | Outer Membrane Protein | 0.028 | 68 | 65 | BETA | 371 | 40102.39 | 4 | 7.5393 | OuterMembrane | 9.93 | LIPO(Sec/SPII) | CS pos: 20-21. ALA-CA. Pr: 0.8574 | 0.92273 | OuterMembrane | 2.519 |
| WP_004184835.1 | porin | Outer membrane porin protein | 500 | Outer Membrane Protein | 0.053 | 65 | 59 | BETA | 362 | 38232.8 | 8.5 | 9.4169 | OuterMembrane | 10 | SP(Sec/SPI) | CS pos: 20-21. AWA-QS. Pr: 0.9064 | 0.81101 | Extracellular | 2.468 |
| WP_004188244.1 | porin | Putative outer membrane porin | 500 | Outer Membrane Protein | 0.056 | 67 | 66 | BETA | 376 | 38969.84 | 11 | 9.8676 | OuterMembrane | 10 | SP(Sec/SPI) | CS pos: 22-23. AHA-QS. Pr: 0.9399 | 0.843456 | OuterMembrane | 2.963 |
| WP_004188361.1 | porin | Putative outer membrane porin | 500 | Outer Membrane Protein | 0.072 | 68 | 62 | BETA | 399 | 41354.82 | 9 | 9.4433 | OuterMembrane | 10 | SP(Sec/SPI) | CS pos: 20-21. AHA-QG. Pr: 0.9602 | 0.805893 | OuterMembrane | 3.403 |
| WP_004193238.1 | porin | Outer membrane porin, putative | 500 | Outer Membrane Protein | 0.037 | 67 | 65 | BETA | 355 | 37611.4 | 15 | 9.9265 | OuterMembrane | 9.93 | SP(Sec/SPI) | CS pos: 32-33. AHA-QS. Pr: 0.9824 | 0.661336 | OuterMembrane | 2.305 |
| WP_004193601.1 | porin | Outer membrane porin OpcP, putative | 500 | Outer Membrane Protein | 0.055 | 67 | 64 | BETA | 407 | 42509.38 | 16.5 | 10.2293 | OuterMembrane | 10 | TAT(Tat/SPI) | CS pos: 37-38. AHA-QS. Pr: 0.9437 | 0.686633 | OuterMembrane | 2.293 |
| WP_004194585.1 | porin | Porin | 500 | Outer Membrane Protein | 0.048 | 71 | 66 | BETA | 363 | 38759.34 | 12.5 | 9.685 | OuterMembrane | 10 | SP(Sec/SPI) | CS pos: 30-31. ALA-QS. Pr: 0.5598 | 0.691884 | OuterMembrane | 3.731 |
| WP_004197824.1 | porin | Outer membrane porin, putative | 500 | Outer Membrane Protein | 0.061 | 72 | 67 | BETA | 377 | 39832.71 | 6 | 9.3022 | OuterMembrane | 9.99 | SP(Sec/SPI) | CS pos: 25-26. AMA-QS. Pr: 0.9492 | 0.79052 | OuterMembrane | 3.935 |
| WP_004199710.1 | porin | Putative outer membrane porin | 500 | Outer Membrane Protein | 0.055 | 69 | 64 | BETA | 400 | 41615.48 | 8 | 8.8331 | OuterMembrane | 10 | LIPO(Sec/SPII) | CS pos: 19-20. APA-CA. Pr: 0.5425 | 0.648889 | OuterMembrane | 2.423 |
| WP_004190808.1 | TonB-dependent copper receptor | TonB-dependent copper receptor | 500 | Outer Membrane Protein | 0.027 | 99 | 96 | BETA | 745 | 80218.01 | 16 | 9.5573 | OuterMembrane | 10 | TAT(Tat/SPI) | CS pos: 43-44. AVA-QT. Pr: 0.7316 | 0.441756 | OuterMembrane | 2.8 |
| WP_004187885.1 | TonB-dependent hemoglobin/transferrin/  lactoferrin | TonB-dependent hemoglobin/transferrin/  lactoferrin | 500 | Outer Membrane Protein | 0.026 | 103 | 97 | BETA | 767 | 82520.92 | 14 | 9.3622 | OuterMembrane | 9.95 | SP(Sec/SPI) | CS pos: 23-24. ARA-AG. Pr: 0.4147 | 0.456666 | OuterMembrane | 4.319 |
| WP_004200389.1 | TonB-dependent receptor | TonB-dependent siderophore receptor family protein | 500 | Outer Membrane Protein | 0.038 | 96 | 90 | BETA | 756 | 80846.56 | 28.5 | 10.1584 | OuterMembrane | 10 | TAT(Tat/SPI) | CS pos: 36-37. AHA-CF. Pr: 0.2440 | 0.3817 | OuterMembrane | 3.097 |
| WP_004190576.1 | TonB-dependent receptor | – | 500 | Outer Membrane Protein | 0.047 | 101 | 94 | BETA | 681 | 72692.55 | 2.5 | 6.793 | OuterMembrane | 10 | OTHER |  | 0.410884 | OuterMembrane | 3.975 |
| WP_004192385.1 | TonB-dependent receptor plug domain-containing protein | Putative TonB-dependent vitamin B12 receptor BtuB | 500 | Outer Membrane Protein | 0.077 | 95 | 86 | BETA | 685 | 72562.49 | 7.5 | 8.7391 | OuterMembrane | 9.95 | SP(Sec/SPI) | CS pos: 22-23. ALA-QG. Pr: 0.5504 | 0.640831 | OuterMembrane | 3.789 |
| WP_011857796.1 | TonB-dependent siderophore receptor | – | 500 | Outer Membrane Protein | 0.027 | 97 | 91 | BETA | 881 | 94662.23 | 22 | 9.4723 | OuterMembrane | 10 | OTHER |  | 0.263948 | OuterMembrane | 3.769 |
| WP_004199226.1 | TonB-dependent siderophore receptor | – | 500 | Outer Membrane Protein | 0.042 | 96 | 88 | BETA | 736 | 80351.76 | 6.5 | 8.5764 | OuterMembrane | 10 | SP(Sec/SPI) | CS pos: 30-31. AQA-AQ. Pr: 0.4836 | 0.703949 | OuterMembrane | 4.53 |
| WP_004197083.1 | outer membrane protein assembly factor BamA | Outer membrane protein assembly factor BamA | 500 | Outer Membrane Protein | 0.034 | 74 | 68 | BETA | 769 | 84848.73 | 12 | 9.3612 | OuterMembrane | 10 | SP(Sec/SPI) | CS pos: 27-28. AHA-  TA. Pr: 0.8768 | 0.782303 | OuterMembrane | 4.801 |
| WP_004189876.1 | LPS-assembly protein LptD | LPS-assembly protein LptD | 500 | Outer Membrane Protein | 0.026 | 114 | 104 | BETA | 787 | 86452.62 | 5 | 7.4853 | OuterMembrane | 10 | SP(Sec/SPI) | CS pos: 39-40. SQA-QL. Pr: 0.4028 | 0.805196 | OuterMembrane | 4.095 |
| WP_004266712.1 | autotransporter assembly complex protein TamA | Outer membrane protein assembly factor | 500 | Outer Membrane Protein | 0.025 | 77 | 70 | BETA | 605 | 65967.24 | 17 | 9.7601 | OuterMembrane | 10 | SP(Sec/SPI) | CS pos: 50-51. AFA-KY. Pr: 0.6935 | 0.453939 | OuterMembrane | 3.617 |
| WP_004197873.1 | TolC family protein | Cobalt-zinc-cadmium resistance efflux protein | 500 | Outer Membrane Protein | 0.057 | 21 | 17 | BETA | 461 | 47477.6 | 9.5 | 10.2539 | OuterMembrane | 10 | SP(Sec/SPI) | CS pos: 23-24. AQA-QP. Pr: 0.5774 | 0.325526 | OuterMembrane | 4.235 |
| WP_004189952.1 | TolC family protein | TolC family protein | 500 | Outer Membrane Protein | 0.02 | 21 | 17 | BETA | 433 | 46817.26 | 8 | 10.0968 | Unknown | 2.5 | SP(Sec/SPI) | CS pos: 34-35. AFA-QS. Pr: 0.5356 | 0.204583 | OuterMembrane | 3.317 |
| WP_004199995.1 | TolC family type I secretion outer membrane protein | Protein CyaE | 500 | Outer Membrane Protein | 0.046 | 18 | 18 | BETA | 482 | 51643.11 | 7.5 | 9.3503 | OuterMembrane | 9.71 | SP(Sec/SPI) | CS pos: 26-27. VHA-QW. Pr: 0.9793 | 0.64796 | OuterMembrane | 4.646 |
| WP_004188259.1 | ShlB/FhaC/HecB family hemolysin secretion/activation | Hemolysin secretion/activation protein, | 500 | Outer Membrane Protein | 0.04 | 78 | 71 | BETA | 554 | 60198.69 | 12.5 | 9.5549 | OuterMembrane | 10 | OTHER |  | 0.295327 | OuterMembrane | 4.458 |
| WP_004193874.1 | ShlB/FhaC/HecB family hemolysin secretion/activation protein | ShlB/FhaC/HecB family hemolysin secretion/activation protein | 500 | Outer Membrane Protein | 0.04 | 75 | 73 | BETA | 551 | 59278.11 | 13 | 9.8778 | OuterMembrane | 10 | SP(Sec/SPI) | CS pos: 15-16. AHA-QS. Pr: 0.8368 | 0.654946 | OuterMembrane | 4.661 |
| WP_011832203.1 | fimbria/pilus outer membrane usher protein | Fimbrial usher family protein | 500 | Outer Membrane Protein | 0.05 | 106 | 98 | BETA | 803 | 86034.48 | 18 | 9.5205 | OuterMembrane | 10 | TAT(Tat/SPI) | CS pos: 34-35. ARA-GE. Pr: 0.8657 | 0.400181 | OuterMembrane | 3.982 |
| WP_011857910.1 | fimbria/pilus outer membrane usher protein | Fimbrial biogenesis outer membrane usher protein | 500 | Outer Membrane Protein | 0.052 | 106 | 96 | BETA | 766 | 80488.42 | 7.5 | 7.9724 | OuterMembrane | 9.92 | OTHER |  | 0.461161 | OuterMembrane | 3.975 |
| WP_004199638.1 | MipA/OmpV family protein | MipA family protein | 500 | Outer Membrane Protein | 0.062 | 60 | 46 | BETA | 248 | 26136.35 | 6.5 | 9.4915 | OuterMembrane | 9.49 | SP(Sec/SPI) | CS pos: 14-15. AQA-EN. Pr: 0.5416 | 0.632067 | OuterMembrane | 2.62 |
| WP_004198784.1 | patatin-like phospholipase family protein | Putative outer membrane protein, OMP85 family | 501 | Outer Membrane Protein | 0.028 | 72 | 70 | BETA | 728 | 78892.44 | 2.5 | 6.8949 | OuterMembrane | 9.83 | OTHER |  | 0.361495 | OuterMembrane | 3.797 |
| WP_004184760.1 | Transporter | Transporter | 500 | Outer Membrane Protein | 0.027 | 56 | 54 | BETA | 247 | 27064.5 | 2.5 | 7.4157 | Unknown | 2.5 | SP(Sec/SPI) | CS pos: 21-22. AHA-DH. Pr: 0.9736 | 0.248616 | OuterMembrane | 1.803 |
| WP_011832365.1 | autotransporter BatA | Outer membrane autotransporter domain protein | 4 | Outer Membrane Protein | 0.044 | 57 | 52 | BETA | 610 | 64322.86 | 12.5 | 9.472 | OuterMembrane | 9.83 | SP(Sec/SPI) | CS pos: 23-24. AWA-YT. Pr: 0.9213 | 0.40532 | OuterMembrane | 2.553 |
| WP_004198495.1 | autotransporter BcaA | Serine protease, subtilase family | 462 | Outer Membrane Protein | 0.043 | 52 | 52 | BETA | 1131 | 116053.8 | 9.5 | 7.3898 | OuterMembrane | 10 | TAT(Tat/SPI) | CS pos: 40-41. AQA-AP. Pr: 0.7440 | 0.800731 | Extracellular | 2.838 |
| WP_004189146.1 | hypothetical protein | Outer membrane protein beta-barrel | 3 | Outer Membrane Protein | 0.03 | 67 | 62 | BETA | 340 | 37282.94 | 16 | 10.0517 | OuterMembrane | 9.52 | TAT(Tat/SPI) | CS pos: 37-38. ARA-QE. Pr: 0.9365 | 0.382602 | OuterMembrane | 2.69 |
| **Group B** |  |  |  |  |  |  |  |  |  |  |  |  |  |  |  |  |  |  |  |
| WP_004203582.1 | efflux RND transporter outer membrane subunit OprB | Outer membrane efflux protein OprB | 500 | Outer Membrane Protein | 0.045 | 19 | 17 | SP | 514 | 54576.65 | 1.5 | 7.2431 | OuterMembrane | 10 | LIPO(Sec/SPII) | CS pos: 18-19. AAG-CT. Pr: 0.9948 | 0.426247 | OuterMembrane | 4.509 |
| WP_004266238.1 | efflux RND transporter outer membrane subunit OprB | RND efflux system, outer membrane lipoprotein, NodT family | 500 | Outer Membrane Protein | 0.053 | 18 | 16 | SP | 512 | 54509.46 | 2 | 7.9461 | OuterMembrane | 10 | LIPO(Sec/SPII) | CS pos: 18-19. AAG-CT. Pr: 0.9884 | 0.411043 | OuterMembrane | 4.609 |
| WP_004188537.1 | efflux transporter outer membrane subunit | Efflux transporter outer membrane subunit | 500 | Outer Membrane Protein | 0.004 | 17 | 15 | SP | 508 | 55253.59 | 12.5 | 9.7354 | OuterMembrane | 10 | LIPO(Sec/SPII) | CS pos: 38-39. VSG-CL. Pr: 0.6200 | 0.126893 | OuterMembrane | 3.769 |
| WP_004188663.1 | efflux transporter outer membrane subunit | Efflux transporter outer membrane subunit | 500 | Outer Membrane Protein | 0.029 | 17 | 19 | SP | 510 | 54642.4 | 3 | 8.6817 | OuterMembrane | 10 | LIPO(Sec/SPII) | CS pos: 29-30. LAA-CA. Pr: 0.9825 | 0.192304 | OuterMembrane | 4.621 |
| WP_004191557.1 | efflux transporter outer membrane subunit | Efflux transporter outer membrane subunit | 500 | Outer Membrane Protein | 0.037 | 19 | 18 | SP | 498 | 53134.84 | 5.5 | 9.0766 | OuterMembrane | 10 | LIPO(Sec/SPII) | CS pos: 24-25. LAG-CA. Pr: 0.9950 | 0.534695 | OuterMembrane | 3.791 |
| WP_004196352.1 | efflux transporter outer membrane subunit | Efflux transporter outer membrane subunit | 500 | Outer Membrane Protein | 0.031 | 21 | 19 | SP | 531 | 56037.11 | 17 | 10.6776 | OuterMembrane | 9.92 | LIPO(Sec/SPII) | CS pos: 35-36. LAG-CV. Pr: 0.9154 | 0.215177 | OuterMembrane | 3.475 |
| WP_004196794.1 | efflux transporter outer membrane subunit | Efflux transporter outer membrane subunit | 500 | Outer Membrane Protein | 0.045 | 17 | 17 | SP | 538 | 55958.87 | 3 | 7.9943 | OuterMembrane | 9.99 | LIPO(Sec/SPII) | CS pos: 27-28. LAG-CA. Pr: 0.9220 | 0.616324 | OuterMembrane | 3.811 |
| WP_024900385.1 | efflux transporter outer membrane subunit | – | 500 | Outer Membrane Protein | 0.039 | 20 | 18 | SP | 588 | 60181.46 | 5 | 7.3495 | OuterMembrane | 10 | LIPO(Sec/SPII) | CS pos: 18-19. MAG-CA. Pr: 0.9977 | 0.421678 | OuterMembrane | 3.245 |
| WP_004197912.1 | OmpW/AlkL family protein | Outer membrane protein, OmpW family | 500 | Non-Outer Membrane Protein | 0.038 | 39 | 35 | BETA | 276 | 28713.78 | 10 | 10.0378 | OuterMembrane | 9.93 | SP(Sec/SPI) | CS pos: 21-22. AHA-QS. Pr: 0.8895 | 0.762521 | Periplasmic | 1.987 |
| WP_004550362.1 | OmpW/AlkL family protein | OmpW family protein | 500 | Non-Outer Membrane Protein | 0.015 | 38 | 36 | BETA | 214 | 22763.27 | 5.5 | 8.4908 | OuterMembrane | 9.93 | SP(Sec/SPI) | CS pos: 28-29. SHA-AS. Pr: 0.9701 | 0.559411 | Extracellular | 1.266 |
| WP_004557213.1 | OmpW/AlkL family protein | OmpW family protein | 500 | Non-Outer Membrane Protein | 0.022 | 39 | 34 | BETA | 274 | 28729.8 | 8 | 9.7798 | OuterMembrane | 9.93 | SP(Sec/SPI) | CS pos: 19-20. AFA-QQ. Pr: 0.9791 | 0.854283 | Extracellular | 3.164 |
| WP_004191391.1 | trimeric autotransporter adhesin BpaB | Hemagglutinin family protein | 1235 | Non-Outer Membrane Protein | 0.086 | 17 | 15 | BETA | 1090 | 105486 | -10.5 | 4.6783 | Unknown | 6.26 | SP(Sec/SPI) | CS pos: 50-51. AWA-DT. Pr: 0.9898 | 0.886715 | Extracellular | 3.231 |
| WP_004199699.1 | trimeric autotransporter adhesin BpaE | Hemagglutinin family protein | 1158 | Non-Outer Membrane Protein | 0.084 | 17 | 15 | BETA | 820 | 78214.04 | -4.5 | 5.1527 | OuterMembrane | 9.95 | SP(Sec/SPI) | CS pos: 62-63. CMA-TD. Pr: 0.1245 | 0.901217 | Extracellular | 3.065 |
| WP_011204221.1 | acyloxyacyl hydrolase | Lipid A deacylase | 500 | Non-Outer Membrane Protein | 0.028 | 39 | 37 | BETA | 189 | 20851.43 | 5 | 9.2589 | Unknown | 2.5 | SP(Sec/SPI) | CS pos: 31-32. AFA-DR. Pr: 0.9568 | 0.702425 | Extracellular | 3.126 |
| WP_004190424.1 | hypothetical protein | Porin | – | Outer Membrane Protein | 0.021 | 80 | 71 | BETA | 432 | 46802.83 | 10 | 10.3427 | Periplasmic | 9.23 | SP(Sec/SPI) | CS pos: 23-24. AAA-TG. Pr: 0.5896 | 0.45235 | OuterMembrane | 2.83 |

**Table S2:** Existing literature-based evidences of the 59 OMBB proteins

| **S. No.** | **Status of prior characterization** | **No. of proteins** | **Protein accession no.** |
| --- | --- | --- | --- |
| 1. | OMPs previously identified experimentally in Bm ATCC 23344 | 3 | WP_004189302.1; WP_004184655.1; WP_004190808.1 |
| 2. | Known β-barrel architecture | 5 | WP_004190638.1; WP_004198784.1; WP_004198495.1; WP_004191391.1; WP_004199699.1 |
| 3. | Conserved immunogenic cross-protective antigens in other *Burkholderia* species | 4 | WP_004199949.1; WP_004550362.1; WP_004197083.1; WP_011832365.1 |
| 4. | Functional homologs experimentally characterized in other Gram-negative bacteria including *Burkholderia* species | 20 | WP_004189876.1; WP_004266712.1; WP_004197873.1; WP_004189952.1; WP_004199995.1; WP_004203582.1; WP_004266238.1; WP_004188537.1; WP_004188663.1; WP_004191557.1; WP_004196352.1; WP_004196794.1; WP_024900385.1; WP_004188259.1; WP_004193874.1; WP_011832203.1; WP_011857910.1; WP_004199638.1; WP_011204221.1; WP_004184760.1 |
| 5. | Annotated in NCBI/UniProt | 25 | WP_004188615.1; WP_004206816.1; WP_053799657.1; WP_004191758.1; WP_004195141.1; WP_004197133.1; WP_004197145.1; WP_004198441.1; WP_004199671.1; WP_004184835.1; WP_004188244.1; WP_004188361.1; WP_004193238.1; WP_004193601.1; WP_004194585.1; WP_004197824.1; WP_004199710.1; WP_004197912.1; WP_004557213.1; WP_004187885.1; WP_004200389.1; WP_004190576.1; WP_004192385.1; WP_011857796.1; WP_004199226.1 |
| 6. | Hypothetical / Novel OMBB proteins | 2 | WP_004189146.1; WP_004190424.1 |

**Table S3:** Structural alignment of 3D models of the identified proteins generated by five modelling tools

| **Protein Accession No.** | **Protein Annotation (NCBI)** | **Confidence Scores****^†^** | | | | | **RMSD (Å)** |
| --- | --- | --- | --- | --- | --- | --- | --- |
|  |  | **AlphaFold 3** | **RoseTTA-Fold** | **TrRosetta** | **ESM-Fold** | **SWISS-MODEL** | **US-align** |
| **Group A** |  |  |  |  |  |  |  |
| WP_004189302.1 | carbohydrate Porin | 0.83 | 0.807 | 0.884 | 0.875 | 0.86 | 2.09 |
| WP_004190638.1 | OprD family Porin | 0.81 | 0.763 | 0.892 | 0.838 | 0.88 | 2.23 |
| WP_004199949.1 | OmpW/AlkL family protein | 0.82 | 0.76 | 0.91 | 0.9 | 0.89 | 1.57 |
| WP_004184655.1 | Porin Omp38 | 0.88 | 0.83 | 0.935 | 0.913 | 0.86 | 2.3 |
| WP_004188615.1 | Porin | 0.9 | 0.853 | 0.935 | 0.905 | 0.93 | 2.34 |
| WP_004206816.1 | Porin | 0.89 | 0.71 | 0.933 | 0.897 | 0.93 | 2.01 |
| WP_053799657.1 | Porin | 0.87 | 0.822 | 0.931 | 0.903 | 0.92 | 2.71 |
| WP_004191758.1 | Porin | 0.9 | 0.841 | 0.92 | 0.9 | 0.93 | 2.34 |
| WP_004195141.1 | Porin | 0.91 | 0.833 | 0.931 | 0.913 | 0.94 | 2.24 |
| WP_004197133.1 | Porin | 0.9 | 0.859 | 0.936 | 0.913 | 0.93 | 2 |
| WP_004197145.1 | Porin | 0.9 | 0.839 | 0.891 | 0.844 | 0.91 | 2.85 |
| WP_004198441.1 | Porin | 0.92 | 0.871 | 0.936 | 0.916 | 0.95 | 2.11 |
| WP_004199671.1 | Porin | 0.89 | 0.855 | 0.924 | 0.896 | 0.93 | 2.29 |
| WP_004184835.1 | Porin | 0.89 | 0.836 | 0.934 | 0.908 | 0.93 | 2.17 |
| WP_004188244.1 | Porin | 0.9 | 0.874 | 0.906 | 0.912 | 0.94 | 2.5 |
| WP_004188361.1 | Porin | 0.9 | 0.859 | 0.924 | 0.903 | 0.94 | 2.13 |
| WP_004193238.1 | Porin | 0.86 | 0.826 | 0.914 | 0.9 | 0.91 | 2.06 |
| WP_004193601.1 | Porin | 0.86 | 0.837 | 0.919 | 0.878 | 0.91 | 2.43 |
| WP_004194585.1 | Porin | 0.87 | 0.82 | 0.931 | 0.913 | 0.93 | 2.16 |
| WP_004197824.1 | Porin | 0.87 | 0.829 | 0.916 | 0.887 | 0.9 | 2.45 |
| WP_004199710.1 | Porin | 0.88 | 0.823 | 0.915 | 0.92 | 0.92 | 2.46 |
| WP_004190808.1 | TonB-dependent copper receptor | 0.85 | 0.855 | 0.888 | 0.861 | 0.86 | 2.33 |
| WP_004187885.1 | TonB-dependent hemoglobin/transferrin/lactoferrin | 0.89 | 0.73 | 0.926 | 0.909 | 0.89 | 2.69 |
| WP_004200389.1 | TonB-dependent receptor | 0.86 | 0.85 | 0.899 | 0.895 | 0.87 | 2.2 |
| WP_004190576.1 | TonB-dependent receptor | 0.91 | 0.78 | 0.948 | 0.937 | 0.91 | 1.84 |
| WP_004192385.1 | TonB-dependent receptor plug domain-containing protein | 0.84 | 0.814 | 0.898 | 0.876 | 0.85 | 1.98 |
| WP_011857796.1 | TonB-dependent siderophore receptor | 0.78 | 0.866 | 0.851 | 0.815 | 0.84 | 3.4 |
| WP_004199226.1 | TonB-dependent siderophore receptor | 0.88 | 0.74 | 0.935 | 0.915 | 0.88 | 1.88 |
| WP_004197083.1 | outer membrane protein assembly factor BamA | 0.77 | 0.83 | 0.906 | 0.806 | 0.89 | 4.98 |
| WP_004189876.1 | LPS-assembly protein LptD | 0.83 | 0.73 | 0.908 | 0.84 | 0.86 | 3.09 |
| WP_004266712.1 | autotransporter assembly complex protein TamA | 0.84 | 0.78 | 0.862 | 0.843 | 0.9 | 3.65 |
| WP_004197873.1 | TolC family protein | 0.77 | 0.803 | 0.883 | 0.88 | 0.85 | 2.95 |
| WP_004189952.1 | TolC family protein | 0.85 | 0.842 | 0.88 | 0.909 | 0.91 | 2.99 |
| WP_004199995.1 | TolC family type I secretion outer membrane protein | 0.84 | 0.77 | 0.868 | 0.909 | 0.86 | 3.51 |
| WP_004188259.1 | ShlB/FhaC/HecB family hemolysin secretion/ activation | 0.89 | 0.78 | 0.946 | 0.919 | 0.89 | 2.42 |
| WP_004193874.1 | ShlB/FhaC/HecB family hemolysin secretion/ activation protein | 0.87 | 0.75 | 0.89 | 0.903 | 0.87 | 2.47 |
| WP_011832203.1 | fimbria/pilus outer membrane usher protein | 0.82 | 0.71 | 0.862 | 0.853 | 0.88 | 3.96 |
| WP_011857910.1 | fimbria/pilus outer membrane usher protein | 0.84 | 0.74 | 0.901 | 0.877 | 0.89 | 3.84 |
| WP_004199638.1 | MipA/OmpV family protein | 0.89 | 0.855 | 0.961 | 0.946 | 0.93 | 1.51 |
| WP_004198784.1 | patatin-like phospholipase family protein | 0.88 | 0.83 | 0.959 | 0.696 | 0.89 | 3.99 |
| WP_004184760.1 | transporter | 0.88 | 0.827 | 0.941 | 0.921 | 0.88 | 1.83 |
| WP_011832365.1 | autotransporter BatA | 0.89 | 0.799 | 0.934 | 0.902 | 0.9 | 2.6 |
| WP_004198495.1 | autotransporter BcaA | 0.76 | 0.874 | 0.746 | 0.87 | 0.91 | 4.32 |
| WP_004189146.1 | hypothetical protein | 0.81 | 0.792 | 0.783 | 0.602 | 0.79 | 3.21 |
| **Group B** |  |  |  |  |  |  |  |
| WP_004203582.1 | efflux RND transporter outer membrane subunit OprB | 0.88 | 0.79 | 0.886 | 0.897 | 0.9 | 3.43 |
| WP_004266238.1 | efflux RND transporter outer membrane subunit OprB | 0.89 | 0.8 | 0.874 | 0.898 | 0.91 | 3.48 |
| WP_004188537.1 | efflux transporter outer membrane subunit | 0.86 | 0.79 | 0.88 | 0.905 | 0.88 | 3.01 |
| WP_004188663.1 | efflux transporter outer membrane subunit | 0.86 | 0.79 | 0.879 | 0.908 | 0.86 | 3.68 |
| WP_004191557.1 | efflux transporter outer membrane subunit | 0.88 | 0.79 | 0.882 | 0.893 | 0.91 | 3.49 |
| WP_004196352.1 | efflux transporter outer membrane subunit | 0.82 | 0.76 | 0.855 | 0.887 | 0.85 | 3.56 |
| WP_004196794.1 | efflux transporter outer membrane subunit | 0.83 | 0.79 | 0.851 | 0.866 | 0.83 | 3.73 |
| WP_024900385.1 | efflux transporter outer membrane subunit | 0.76 | 0.77 | 0.794 | 0.784 | 0.79 | 3.61 |
| WP_004197912.1 | OmpW/AlkL family protein | 0.85 | 0.85 | 0.942 | 0.847 | 0.91 | 1.78 |
| WP_004550362.1 | OmpW/AlkL family protein | 0.79 | 0.82 | 0.908 | 0.856 | 0.87 | 1.39 |
| WP_004557213.1 | OmpW/AlkL family protein | 0.82 | 0.85 | 0.944 | 0.885 | 0.9 | 1.4 |
| WP_004191391.1 | trimeric autotransporter adhesin BpaB | 0.6 | 0.814 | 0.275 | 0.465 | 0.81 | 2.18 |
| WP_004199699.1 | trimeric autotransporter adhesin BpaE | 0.61 | 0.73 | 0.578 | 0.469 | 0.81 | 1.28 |
| WP_011204221.1 | acyloxyacyl hydrolase | 0.8 | 0.82 | 0.874 | 0.853 | 0.9 | 2.28 |
| WP_004190424.1 | hypothetical protein | 0.84 | 0.814 | 0.869 | 0.77 | 0.87 | 2.31 |

† AlphaFold’s ranking score combines ptm, iptm, disorder, clashes into a single value from -100 to 1.5; higher scores indicate better model quality. RoseTTAFold confidence score with values above 0.7-0.8 suggests high model accuracy. TrRosetta’s TM-score evaluates global structural similarity between predicted and native structures; scores above 0.5 imply correct topology. ESMFold pLDDT score ranges from 0 to 1 where higher pLDDT scores (closer to 1.0) indicate higher confidence in the accuracy of the predicted structure. SwissModel’s GMQE estimates overall model quality based on alignment and template structure; values above 0.6-0.7 indicate good reliability.

**Table S4:** 33 strains of *B. mallei* with complete genome searched for predicted OMBB’s variations

| **Assembly Accession** | **Strain Name** | **Genome Size (Mb)** | **CDS**^†^ | **BioSample ID** | **Host** | **Location** | **Submitter** |
| --- | --- | --- | --- | --- | --- | --- | --- |
| GCF_000011705.1 | ATCC 23344 | 5.836 | 4,964 | [SAMN02603987](https://www.ncbi.nlm.nih.gov/biosample/SAMN02603987/) | NA | NA | TIGR |
| GCF_000015465.1 | SAVP1 | 5.232 | 4,468 | [SAMN02604034](https://www.ncbi.nlm.nih.gov/biosample/SAMN02604034/) | NA | NA | TIGR |
| GCF_000015605.1 | NCTC 10229 | 5.742 | 4,887 | [SAMN02604032](https://www.ncbi.nlm.nih.gov/biosample/SAMN02604032/) | NA | NA | TIGR |
| GCF_000015625.1 | NCTC 10247 | 5.848 | 4,973 | [SAMN02604033](https://www.ncbi.nlm.nih.gov/biosample/SAMN02604033/) | NA | NA | TIGR |
| GCF_000755785.1 | FMH 23344 | 5.836 | 4,934 | [SAMN02945023](https://www.ncbi.nlm.nih.gov/biosample/SAMN02945023/) | Homo sapiens | Myanmar | Los Alamos National Laboratory |
| GCF_000755845.1 | 6 | 5.648 | 4,755 | [SAMN02837932](https://www.ncbi.nlm.nih.gov/biosample/SAMN02837932/) | Human | NA | Los Alamos National Laboratory |
| GCF_000755865.1 | 23344 | 5.625 | 4,781 | [SAMN02821273](https://www.ncbi.nlm.nih.gov/biosample/SAMN02821273/) | Knee fluid | Myanmar | Los Alamos National Laboratory |
| GCF_000755885.1 | BMQ | 5.63 | 4,807 | [SAMN02839409](https://www.ncbi.nlm.nih.gov/biosample/SAMN02839409/) | horse | India | Los Alamos National Laboratory |
| GCF_000756025.1 | 2E+09 | 5.875 | 4,979 | [SAMN02849482](https://www.ncbi.nlm.nih.gov/biosample/SAMN02849482/) | NA | Hungary | Los Alamos National Laboratory |
| GCF_000762285.1 | NCTC 10247 | 5.828 | 4,921 | [SAMN02798191](https://www.ncbi.nlm.nih.gov/biosample/SAMN02798191/) | NA | Turkey: Ankara | Los Alamos National Laboratory |
| GCF_000959165.1 | 2E+09 | 5.742 | 4,863 | [SAMN03010440](https://www.ncbi.nlm.nih.gov/biosample/SAMN03010440/) | Guinea pig | Hungary | Los Alamos National Laboratory |
| GCF_000959405.1 | 11 | 5.913 | 5,018 | [SAMN03079578](https://www.ncbi.nlm.nih.gov/biosample/SAMN03079578/) | Human | Turkey | Los Alamos National Laboratory |
| GCF_000959465.1 | India86-567-2 | 5.686 | 4,894 | [SAMN03107083](https://www.ncbi.nlm.nih.gov/biosample/SAMN03107083/) | Mule | India | Los Alamos National Laboratory |
| GCF_000959485.1 | 2E+09 | 5.409 | 4,662 | [SAMN03120828](https://www.ncbi.nlm.nih.gov/biosample/SAMN03120828/) | NA | United Kingdom | Los Alamos National Laboratory |
| GCF_000959625.1 | 2E+09 | 5.78 | 4,915 | [SAMN03222861](https://www.ncbi.nlm.nih.gov/biosample/SAMN03222861/) | NA | NA | Los Alamos National Laboratory |
| GCF_001729545.1 | Bahrain1 | 5.78 | 4,907 | [SAMN05607081](https://www.ncbi.nlm.nih.gov/biosample/SAMN05607081/) | Equus caballus | Bahrain | Friedrich Loeffler Institut, Federal |
| GCF_002345985.1 | Turkey1 | 5.592 | 4,764 | [SAMN03121648](https://www.ncbi.nlm.nih.gov/biosample/SAMN03121648/) | NA | Turkey | USAMRIID |
| GCF_002346005.1 | Turkey10 | 5.729 | 4,922 | [SAMN03121657](https://www.ncbi.nlm.nih.gov/biosample/SAMN03121657/) | NA | Turkey | USAMRIID |
| GCF_002346025.1 | Turkey2 | 5.592 | 4,767 | [SAMN03121649](https://www.ncbi.nlm.nih.gov/biosample/SAMN03121649/) | NA | Turkey | USAMRIID |
| GCF_002346045.1 | FMH | 5.835 | 4,960 | [SAMN03174435](https://www.ncbi.nlm.nih.gov/biosample/SAMN03174435/) | Homo sapiens | USA:Maryland | USAMRIID |
| GCF_002346065.1 | Turkey3 | 5.663 | 4,871 | [SAMN03121650](https://www.ncbi.nlm.nih.gov/biosample/SAMN03121650/) | NA | Turkey | USAMRIID |
| GCF_002346085.1 | Turkey4 | 5.733 | 4,947 | [SAMN03121651](https://www.ncbi.nlm.nih.gov/biosample/SAMN03121651/) | NA | Turkey | USAMRIID |
| GCF_002346105.1 | Turkey5 | 5.698 | 4,899 | [SAMN03121652](https://www.ncbi.nlm.nih.gov/biosample/SAMN03121652/) | NA | Turkey | USAMRIID |
| GCF_002346125.1 | Turkey6 | 5.748 | 4,907 | [SAMN03121653](https://www.ncbi.nlm.nih.gov/biosample/SAMN03121653/) | NA | Turkey | USAMRIID |
| GCF_002346145.1 | Turkey7 | 5.725 | 4,901 | [SAMN03121654](https://www.ncbi.nlm.nih.gov/biosample/SAMN03121654/) | NA | Turkey | USAMRIID |
| GCF_002346165.1 | Turkey8 | 5.694 | 4,860 | [SAMN03121655](https://www.ncbi.nlm.nih.gov/biosample/SAMN03121655/) | NA | Turkey | USAMRIID |
| GCF_002346185.1 | Turkey9 | 5.774 | 4,917 | [SAMN03121656](https://www.ncbi.nlm.nih.gov/biosample/SAMN03121656/) | NA | Turkey | USAMRIID |
| GCF_002346205.1 | JHU | 5.737 | 4,926 | [SAMN03174429](https://www.ncbi.nlm.nih.gov/biosample/SAMN03174429/) | Homo sapiens | USA:Maryland | USAMRIID |
| GCF_033870355.1 | Zagreb | 5.822 | 4,950 | [SAMN34156473](https://www.ncbi.nlm.nih.gov/biosample/SAMN34156473/) | Equus caballus | Yugoslavia | Canadian Food Inspection Agen |
| GCF_033870375.1 | Mukteswar | 5.755 | 4,893 | [SAMN34156472](https://www.ncbi.nlm.nih.gov/biosample/SAMN34156472/) | Equus caballus | India | Canadian Food Inspection Agen |
| GCF_033870395.1 | Bogor | 5.823 | 4,955 | [SAMN34156471](https://www.ncbi.nlm.nih.gov/biosample/SAMN34156471/) | Equus caballus | Indonesia | Canadian Food Inspection Agen |
| GCF_033956065.1 | ATCC 23344 | 5.835 | 4,965 | [SAMN34156470](https://www.ncbi.nlm.nih.gov/biosample/SAMN34156470/) | Homo sapiens | Myanmar | Canadian Food Inspection Agen |
| GCF_939576165.1 | 34 | 5.647 | 4,806 | [SAMEA14091970](https://www.ncbi.nlm.nih.gov/biosample/SAMEA14091970/) | NA | NA | FRIEDRICH-LOEFFLER-INSTITUT |

†CDS: Coding Sequences
NA: Information not available on NCBI

**Table S5:** Amino acid sequence variations in *B. mallei* turkey2 OMBBs across 33 *B. mallei* ­strains

| **Protein Accession No.** | **Protein Annotation (NCBI)** | **Total no. of variations** | **Amino acid residues** |
| --- | --- | --- | --- |
| **Group A** |  |  |  |
| WP_004189302.1 | Carbohydrate Porin | – | – |
| WP_004190638.1 | OprD family Porin | 6 | M1-S6 |
| WP_004199949.1 | OmpW/AlkL family protein | 4 | M1-K4 |
| WP_004184655.1 | Porin Omp38 | 2 | Y302, T303 |
| WP_004188615.1 | Porin | – | – |
| WP_004206816.1 | Porin | 176 | M1-T4, H6, A9-F16, L18-H23, I33, V34, A36, G37, A39, N43-Q45, A47-S59, S64, R71, G78, A81, L83, L92-T94, Q96-N98, N100, E102, K106, I108, A112-W116, S118, V119, V128, G134, L135, E137, G143, V144, L151, Y154, S161, S163, L171, S172, G183, V185, A188, T189, S191, R193, Y195, F197, S200, A202, L206-G209, L213, A215-R226, T229, T234, L235, N237-V239, Q242, F244, A245, Q251, V253-G256, L261, S263, A264, F266, A269, T273, R276, R277, S279, L280, A282, D285, A287, F289, S291, S293, A294, F296, N297, F300, S301, R305, A306, V308, N311, S314-T316, P318-S320, H322, N327, L328, G331, L340, A342-G345, K348, H352-D362, V367, A372, S374, D377, L381-V384, I386 |
| WP_053799657.1 | Porin | 1 | E64 |
| WP_004191758.1 | Porin | – | – |
| WP_004195141.1 | Porin | 1 | R113 |
| WP_004197133.1 | Porin | – | – |
| WP_004197145.1 | Porin | – | – |
| WP_004198441.1 | Porin | – | – |
| WP_004199671.1 | Porin | – | – |
| WP_004184835.1 | Porin | – | – |
| WP_004188244.1 | Porin | 274 | M1-K5, I7-A9, L12, V14-A17, L19, H21, A31, L32, V36-Q39, T41-T49, N52, T53, A55-V57, F59, D61, G63-S66, L68, I71, K72, T74, Y81-N84, Q88-F91, G93-T95, K97, G99-P104, A106-A109, N111-I113, L116, V118, S119, P121, F122, T124-A127, Q130-V132, M134-A137, A139-T141, V143-L157, L159-G164, P166-N168, N171-I174, A176-D179, D181-L183, Y185-P188, F190, A193-E198, A200, P201, G203, G207-V217, K219, A221, N223-A240, G242-V256, A258, Q259, K261-S268, S270, G272, G274, N276-L285, F288-F290, L292, G293, R295-S297, F299, R301-V306, Y308-S319, S321-V323, G325, D327, E329, H333, T335, Q339, R342-P358, A360-L370, R373-S375 |
| WP_004188361.1 | Porin | – | – |
| WP_004193238.1 | Porin | – | – |
| WP_004193601.1 | Porin | – | – |
| WP_004194585.1 | Porin | 238 | K3-A8, T10-A18, C20-L23, S25-H27, L29, Y37, I39, M40, A42-E45, V47-G55, A57, R59, K61, S62, N64-T67, W70, R73, V75, V85, R87, I92, D93, A95, N96, D100, G102-S105, A108, R110, T112, K116, G117, W119, E121, L122, N127, F128, P130-Y132, Y134-F138, M141-S148, A150-D161, F164-S168, A170, R172, D174, A176, Y177, R181, F182, G187, N190, V191, S194, K196-K200, D202, A204, G206, E208, S209, F212-I240, G243, S245, D247, G249-L251, T253, M254, R258-S273, M275-L278, G280, Y282-F284, F288, S289, L294, H296-G302, D304, D306-T308, S311-Q315, V323, L324, A326-G329, A331-N338, S340-T351, L353, T356, V357, M359-R362 |
| WP_004197824.1 | Porin | – | – |
| WP_004199710.1 | Porin | – | – |
| WP_004190808.1 | TonB-dependent copper receptor | 588 | M1-E47, V49-G51, T54, L55, S58-A88, D91, T94, P96, T97, E99-V101, A103, E105-A128, Y130, K132-I134, F137-S142, G144-N146, D148, V150, L151, F155, S157, L159-I161, A163, M166-G170, C172-M176, A178-E186, Y188-L193, K195, Q198-V200, P204, S205, S207, A208, T210, L212-T217, F220-P223, M225-G234, F236, R238-D244, T246, A247, T249, P250, F252, Y253, V256-N259, Q264-N270, R272-N282, D284-L287, W289, T290, D292-T295, L297-T303, D305, G306, A308, R309, A311, R313-D316, A318-F320, R322, T324-G326, K328-F346, N348-D355, T358-P362, P364-V376, R378-G382, A386, T388, L389, L391-A394, K396-T399, D402-S405, R407-Y419, D421-N425, Q427-A435, E438-T440, Y442, S444, S447-G451, D456-T467, G469-T478, D480-L482, V486-S489, F491, R493-D497, A499, S500, P502-T504, Y506-Q513, F515, D517-W519, F522-R526, N529, S531-K539, E541-L546, I548-Q551, K553, D555-V561, A563-G566, V568-D575, A577, P580-N589, N591, Q593-M595, G597, V599-S602, P605-F611, G613, L615, A616, A618-G620, V623-S625, A627, Q631-P633, E636, R638-E642, T644, R645, W648, S649, G651-P659, Y663-A679, F681-V683, S685-Q689, N691-S699, V702, D703, V705, D707-H713, N715-G718, A720-G723, P725-P733, T736-S742, K744, L745 |
| WP_004187885.1 | TonB-dependent hemoglobin/transferrin/lactoferrin | 4 | A41-S43, S87 |
| WP_004200389.1 | TonB-dependent receptor | 2 | A120, H260 |
| WP_004190576.1 | TonB-dependent receptor | 1 | M1 |
| WP_004192385.1 | TonB-dependent receptor plug domain-containing protein | 1 | L162 |
| WP_011857796.1 | TonB-dependent siderophore receptor | 624 | P71-R76, V79, R81, V82, A84, A85, L87-S89, C91-P134, E136-S143, L146-L177, L180, P183-G187, D189, A191, L192, D195, T196, T200-A202, R204-P216, E218-A220, L222-S224, A226, L228, A229, V231, A234-I237, A239, Q240, V242-D253, L255-S259, I261-K276, G280-R283, S286-R289, G291-G297, S299-T304, V307, E308, K311, A314, L316, I320, M321, G325-N328, V330, T331, Q333-A341, S343, A344, S347-Y349, G351-E357, T359, S362-A365, E368-A372, L375-V396, A401, Y403-V410, S412-E414, R416-D429, R431-P435, A437, I438, A440-L444, E446, P447, N449-M451, G453-D463, Q465, A467, D469, K471-Y483, A485-V493, L496-G498, L500, S503-A519, V521-V525, L527-R531, D533-Q535, V538, G540-M550, R552-N563, T565, G567-V569, P571-S576, D579, S580, Q582-D584, H587-S590, F592, F593, S596, H598-E601, M604, V606-R610, V612, Y614, L617-P623, Q625-P637, A639, I641, V642, K644-A648, S650, L651, G653, T656, Q657, L659, K660, T662-A669, Y672-A679, E681, E682, A684-W686, L688, A690-M694, G697, A699-L702, F704-D706, D708-H711, L713, V714, Q716-D718, A720, N722-S729, R731, A732, I737, D740-S742, R744-N750, S754, A756, I758, A760, T762-L773, N775, A777-T780, A784, V786-V789, T791-I800, A803, G804, V807, G808, R810, P811, D813-A815, F818, L820, A822, A824-A826, F829-T831, D833-Q842, Q844, V847, K848, T854, Y856-V867, D869, A870, Q872, S874-F881 |
| WP_004199226.1 | TonB-dependent siderophore receptor | – | – |
| WP_004197083.1 | outer membrane protein assembly factor BamA | – | – |
| WP_004189876.1 | LPS-assembly protein LptD | – | – |
| WP_004266712.1 | autotransporter assembly complex protein TamA | 2 | F390, S507 |
| WP_004197873.1 | TolC family protein | – | – |
| WP_004189952.1 | TolC family protein | – | – |
| WP_004199995.1 | TolC family type I secretion outer membrane protein | – | – |
| WP_004188259.1 | ShlB/FhaC/HecB family hemolysin secretion/activation | – | – |
| WP_004193874.1 | ShlB/FhaC/HecB family hemolysin secretion/activation protein | – | – |
| WP_011832203.1 | fimbria/pilus outer membrane usher protein | 546 | M1-F26, A28, G30, H31, R33-T37, T39-L48, P50-G53, A55-T59, E63-V66, A68-S70, G72, R73, P76-R80, I83, Y84, R87-G89, A92-S95, T98-D103, V106, D107, S109, R110, V114-E117, E119, S120, E122, R124-K126, T128, V129, P131-Q137, L139-R143, Y145-T148, A151-F154, L156-F158, V162, N165-A176, T178, Q180, L182, R185, W186, T188-N191, Y195-V206, S208, N209, L212, F217, R219, Y220, Q223-R225, R227, Y229-A231, V234, T236-A238, S240, S242-V245, L247-D255, K257, V258, I262, Y265, L267, Q269-S271, Q273, A275, T278, A279, F283, I284, S287-T291, Q293, N295, P296, T300, M301, N303, F306, N308, E312, T318, L321, Q324, V325, T328, I329, F331, V333-L337, Q339, K340, S343, D344, L347, S348, A351, M352, D355, Y356, I358, S362, G364-A368, G370, H374, L376-Y379, L382, G384, V386, G388-G395, L397, F399-I403, M405, F406, L409-A412, T414, R417-S422, R424, Y426-F428, S431-Q435, V439-L441, R443-R451, S454-Y456, L458-V467, S470, Q472-G489, F491-V493, G495-T501, A504-L506, T509-R515, T517-A520, V522-E528, G530-Q536, I538, P540, E543-V546, T548-R562, Q564-S566, S568-D572, L575, N578-G583, S586, H587, Q589, D592, T594-Q603, G605, Y607-E620, Q622, S624, V627, A631, V632, P634-N636, V638, A641, V643, I645, D646, Q648-G651, R656, Y657, Q660, V662, K664, G667, G668, H670, V673, W675-S678, Y680-E685, P688, D690, S693-V695, A697-E701, R703, A705-G710, A712, V714, T715, P717, R719, I721-C723, Q725, A727, V729, A732, R734, V736-I738, S740, R741, L743, E745, E749-L752, W755, Q756, E758, Y760-E762, S765, L767, R771-D776, R778, T779, R781-T783, A785-V796, A800, G802, E803 |
| WP_011857910.1 | fimbria/pilus outer membrane usher protein | 4 | D218, S275, L290, Q517 |
| WP_004199638.1 | MipA/OmpV family protein | 1 | A197 |
| WP_004198784.1 | patatin-like phospholipase family protein | 206 | M1, D3-I12, L14, V15, S17, G20, Y24, L27, V32, N36, R37, D41-I43, A45, M48, A50, G54, T58, T61, Q63-L74, D76-Y98, D100-G105, D107, K109-V115, V118, N121, R122, Q124-A128, W130-T136, Q138, F140-I145, F147-I150, Q155, T156, K159-H164, S166-P168, I171, M175-L177, G179, L180, S182, A184, E185, D187, G188, A190, L191, G198-L200, V202-A204, A207, V212, G219, P221-S230, A232-Q236, M238-G240, L242-N257, I259-Q262, D264, G266-T271, Q274-A276, Q278, A281, A285, A287-L291, R293, A295, Y297-E305, Y307, R308 |
| WP_004184760.1 | transporter | – | – |
| WP_011832365.1 | autotransporter BatA | 1 | S492 |
| WP_004198495.1 | autotransporter BcaA | 520 | A471-N473, D475-Y479, V483, R485-Q490, R492, E493, Q495, S497-G502, G504-T507, A509, A511-Y514, P517-D521, G523-D528, S530, A532, A535-N538, A540-A550, I552, V554, G557-A562, T564-A566, L568-G580, T582-V585, V587-S589, A592-N600, A602-T650, T652, A654, N656-S664, T667-G672, V674-G679, I681-V684, V686-P688, G690-R700, Q702, D704-L708, L710, A711, E713, T715-P718, T720-G729, R731, E733, A737, V741-S749, Y751, F753-T764, S766-A780, T782-R786, V788, A789, A791-P797, A800-F807, S810-I825, P827, A829, A831, L833, V834, E836-V840, E842, T843, E846, R847, W849, T850, R852, A854, G856-G859, A862-L864, G866-G873, D875, V876, G878-T880, S882-G885, L887, A888, A892, A893, L895-V898, G902-N912, R914-H926, A929, A931-A937, G939, I942, G943, A945-A947, H949, G951-V953, V957-E966, T968-E980, R984, D988, A990-T992, E994, F996-G998, A1000, V1002, H1003, K1005-Q1007, T1009, T1012, G1014-Q1022, N1024-D1026, T1028, S1030-L1032, V1034, G1036-G1041, T1043-L1046, T1049-Q1051, S1053-D1062, Q1064, S1066, L1069-G1074, D1076, F1078-S1081, P1084-G1104, L1106-S1111, S1113, A1115-S1126, H1128-K1130 |
| WP_004189146.1 | hypothetical protein | – | – |
| **Group B** |  |  |  |
| WP_004203582.1 | efflux RND transporter outer membrane subunit OprB | 2 | C231, A501 |
| WP_004266238.1 | efflux RND transporter outer membrane subunit OprB | 21 | R2-V10, A12, F14, F15, A319, S502-S504, R506, T507, A509-G512 |
| WP_004188537.1 | efflux transporter outer membrane subunit | 301 | M1-A28, L30, L32-S37, L40, S41, D45, A47, K49-P51, P54, A55, Y57, Y60, E63-G65, G67-L74, A78, Y79, A87, T91, A94, I101-R105, Q108, A109, A112, G114, R116, D119-Y121, I124-F133, E135, G137, L139-S143, I145, G147-L149, S151-Q158, I161, F163, W164, R168, A172, A173, D176-L179, S181-S184, R18s6, V188-V190, I193-Q195, A199, A202, C204-Y206, E208, R209, A211, L212, N214, E215, I217, A218, R220, D222, L224-E233, A235, I236, K238, T242, S244-Q250, Q252-Q260, D263-H267, D270, V273, T275-T281, S284, D286-P292, L294-P296, E301, A304, N305, V309-A311, Y314, Q315, H320, R333, A335, S339, I340, G343, E346, H348, N349, A352, S353, N359, I361, L364-V366, D370, A371, R373, G376, K379, N382-L389, Q391, T395, T398, D402, A404, A406, H411-D415, V417-T426, A428, E429, A431, R432, A434, K435, D439, S440, A442-F445, E447, D450, R453, L456, N457, V463-T465, R467, L469-R473, A475, A478, R483-D485, P487, R489-A493, R495-S497, E499-A502, Q504, G505, L507, S508 |
| WP_004188663.1 | efflux transporter outer membrane subunit | 316 | M1-I10, V13, K15, I16, A19-A25, A29, R37, V41, T43, F47, A52-T61, A64, E66, A70, H72, E75, R78-G81, P83, V84, D86, E89-L93, A95, N98-A102, R105-E107, A112-A115, S118-W120, Q123, V124, V126-T131, E133-A138, Q140-Q144, S146-P148, N150-V160, Y162, A164, F167, R169, G171, N173, E175, S177, R178, D180, Q183-Q185, L187-Q192, A194, L195, D198, V199, Q201, N202, E205, R208, L209, S211, D212, D214, Y216-R218, G221-A226, K228, V230, R232-F234, E236, D238-L242, S245, R246, K248-E250, A252, T253, D257, V259-R264, A267, S268, L272, I274, L276, K278-D282, F285-T288, I290-V297, A299-L301, L305, I312, A318-M320, A324, R325, L328, K330, S331, Y333, K336, D338-F343, Y345-N352, L355-S358, T360-L363, F366-A370, T372-I375, G378, R380, S382, G384-Q387, K391, E394, E395, N398, Q401, Q402, V405, R408, E409, D416, L417, L419, D421-Q423, R425, A426, S428, D429, N432-T443, Q446, E447, A449, S451, D454, I456-Q471, T473, T475-A477, T480, N482, I484, R485, G492-P502, K504, D506, V507, R510 |
| WP_004191557.1 | efflux transporter outer membrane subunit | – | – |
| WP_004196352.1 | efflux transporter outer membrane subunit | 20 | M1-L15, L162, R308-A311 |
| WP_004196794.1 | efflux transporter outer membrane subunit | 1 | R349 |
| WP_024900385.1 | efflux transporter outer membrane subunit | 12 | S499-A510 |
| WP_004197912.1 | OmpW/AlkL family protein | 1 | Q129 |
| WP_004550362.1 | OmpW/AlkL family protein | – | – |
| WP_004557213.1 | OmpW/AlkL family protein | – | – |
| WP_004191391.1 | trimeric autotransporter adhesin BpaB | 320 | P648, Q650, S652-S723, T725-V739, A741, I743-N757, T759, K760, D762-A768, S770, V771, P775-T793, Q795, G797-T803, N805-T815, I817, T819-A824, G826-I831, N833-S842, T844-A846, N848-S850, Y853-Q857, I859-L862, D864-M866, L869, N871-N875, L877-V924, T926, T928-M931, V933, S935, G936, V938-T940, S944, T949-G955, S957, V959, A964, P965, A967, V971-N973, L986, M989, G991, L993-F1000, L1004, A1006, V1007, D1010-V1014, A1017, G1020, I1022, E1033, K1036, T1039-V1042, I1044, G1047, T1048, R1050, Y1052, Q1053, A1059, T1064, I1067, G1074, S1077, T1080-A1082, A1086, M1088 |
| WP_004199699.1 | trimeric autotransporter adhesin BpaE | – | – |
| WP_011204221.1 | acyloxyacyl hydrolase | – | – |
| WP_004190424.1 | hypothetical protein | – | – |

– indicates no amino acid variation

**Table S6:** Top 20 conformational B-cell epitopes predicted by ElliPro

| **Protein Accession No.** | **Residues** | **Number of Residues** | **Score^†^** |
| --- | --- | --- | --- |
| WP_004197083.1 | M1, L2, F3, K4, P5, H6 | 6 | 0.995 |
| WP_004199671.1 | M1, G2, F3 | 3 | 0.995 |
| WP_053799657.1 | M1, D2, K3 | 3 | 0.994 |
| WP_004193238.1 | M1, T2, T3, L4 | 4 | 0.993 |
| WP_011857796.1 | M1, D2, S3, I4, C5, N6, R7 | 7 | 0.993 |
| WP_004184835.1 | V2, K3, H4 | 3 | 0.992 |
| WP_004199949.1 | M1, K2, Q3 | 3 | 0.992 |
| WP_004193601.1 | M1, S2, A3, R4, A5, A6 | 6 | 0.991 |
| WP_004197824.1 | M1, K2, Q3, T4, T5, K6 | 6 | 0.991 |
| WP_011832365.1 | T2, I3, G4, S5, K6, T7, K8 | 7 | 0.99 |
| WP_004199995.1 | M1, I2, A3, M4 | 4 | 0.989 |
| WP_004206816.1 | Q3, T4, K5 | 3 | 0.989 |
| WP_004199671.1 | Q4, S5, K7 | 3 | 0.986 |
| WP_004199794.1 | K2, A3, K4 | 3 | 0.986 |
| WP_004200389.1 | M1, S2, R3, A4, P5, H6, A7, P8, S9, R10 | 10 | 0.985 |
| WP_011204221.1 | N2, D3, K4 | 3 | 0.984 |
| WP_011857796.1 | R8, P9, R10, R11 | 4 | 0.982 |
| WP_004189952.1 | R6, A7, P8, H9 | 4 | 0.982 |
| WP_004190808.1 | M1, T2, S3, T4, F5, L6, R7, H8, A9, P10, A11, A12, R13 | 13 | 0.981 |
| WP_004190576.1 | M1, S2, L3, L4, L5, A6, A7, S8, L9, A10, H11, G12, E13, T14, G15, A16, P17, P18, A19, E20 | 20 | 0.98 |

†Score for conformational epitopes reflects the protrusion index (PI) of predicted antibody-binding residues on a protein’s 3D structure. It ranges from 0 to 1, where a higher score indicates greater surface accessibility.

**Table S7:** Tertiary structure assessment of MEV models after refinement

| **Modeling Tools^≠^** | **Rama favoured residues (%)** | **Clash score** | **GDT-HA** |
| --- | --- | --- | --- |
| I-TASSER | 86.3 | 12.1 | 0.90 |
| ESMFold | 98.1 | 5.9 | 0.87 |
| AlphaFold 3 (Round 1) | 99.4 | 6.3 | 0.94 |
| AlphaFold 3 (Round 2) | 99.4 | 5.5 | 0.97 |

**≠** RoseTTAFold2 structure could not be refined because of backbone discontinuities in the predicted model.

**Supplementary Figures**

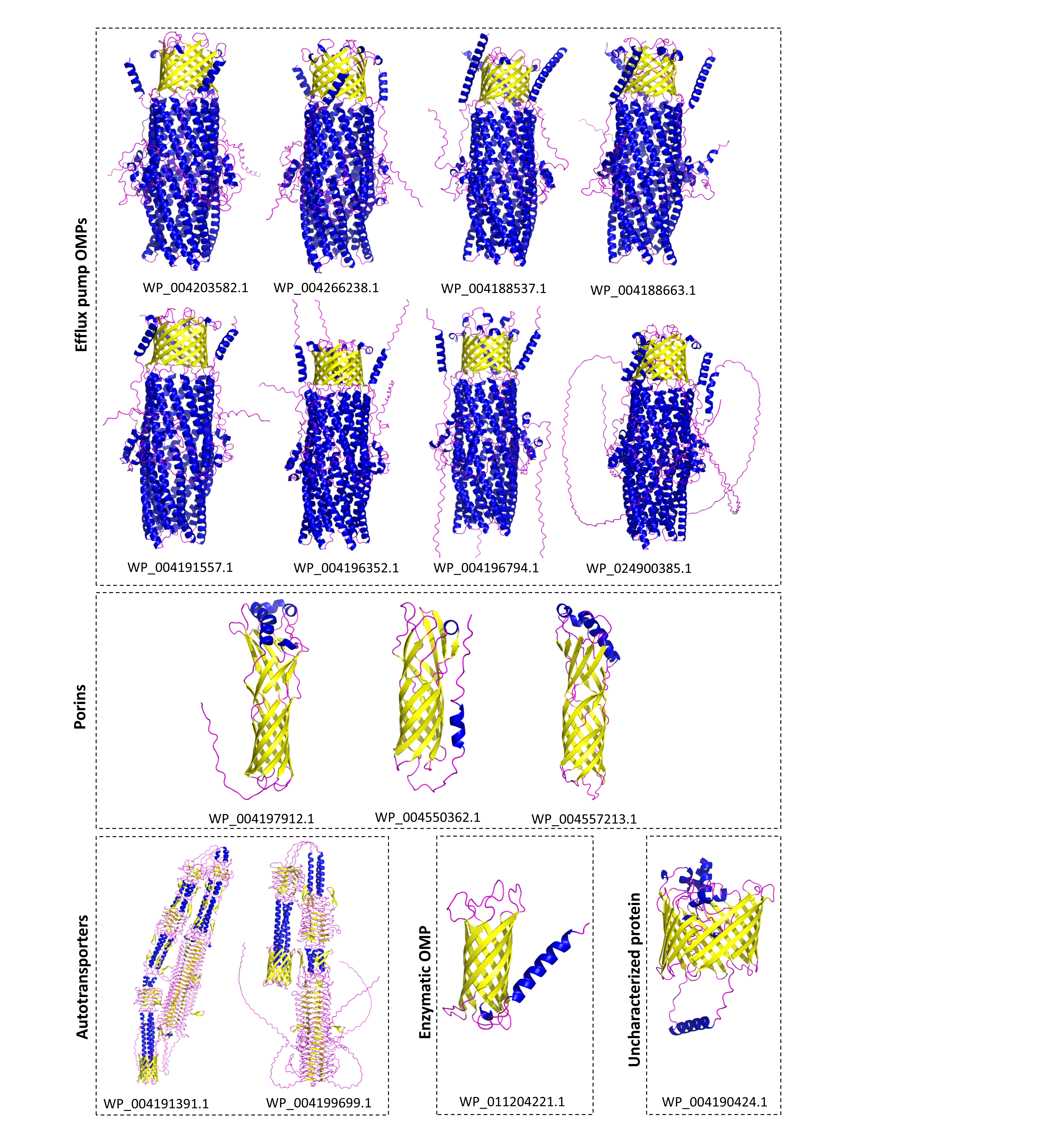

**Figure S1.** Structural models of 15 OMBB proteins in Group-B (predicted as β-barrels by any four OMBB prediction tools) generated using AlphaFold 3.

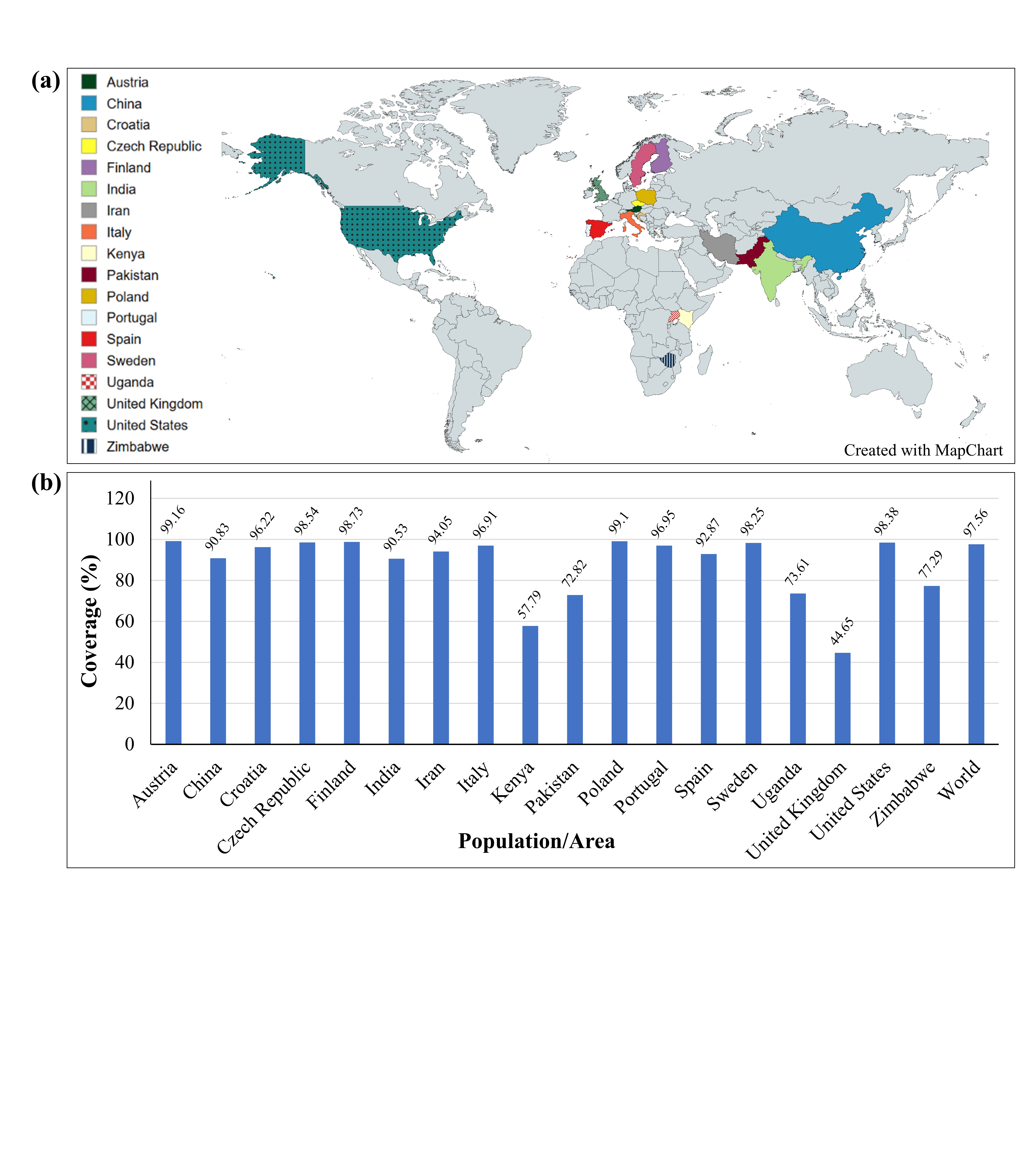

**Figure S2. Population coverage analysis of nine epitopes selected for MEV design in glanders-endemic regions. (a)** Geographical distribution of glanders across the 18 countries included in the population coverage analysis. **(b)** Predicted % of population coverage of the selected epitopes across 18 countries and the world.

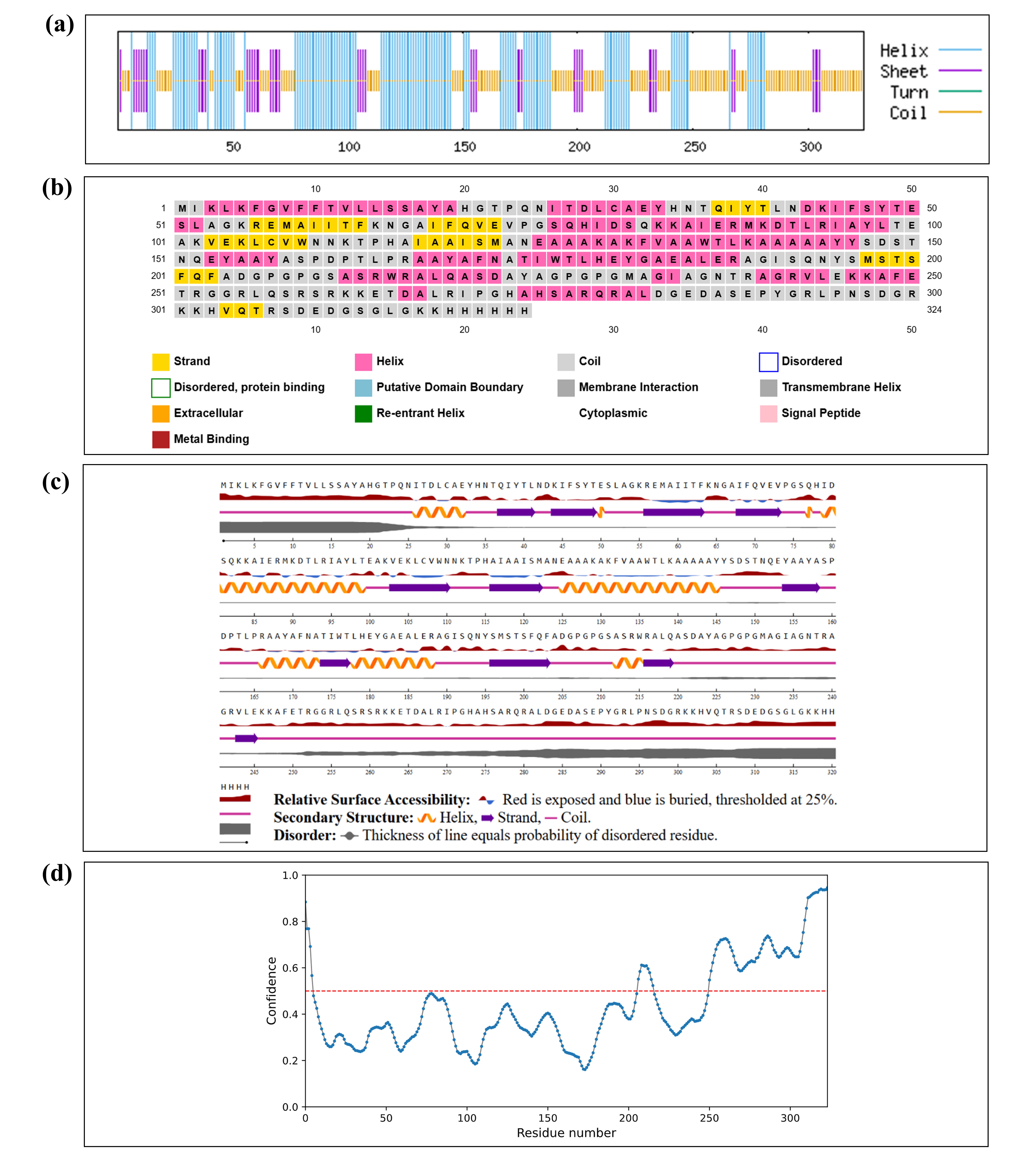

**Figure S3. Secondary structure analysis of MEV construct.** Secondary structure profile predicted by **(a)** SOPMA, **(b)** PSIPRED 4.0, and **(c)** NetSurfP. **(d)** Amino acid residues forming IDRs as predicted by PrDOS.

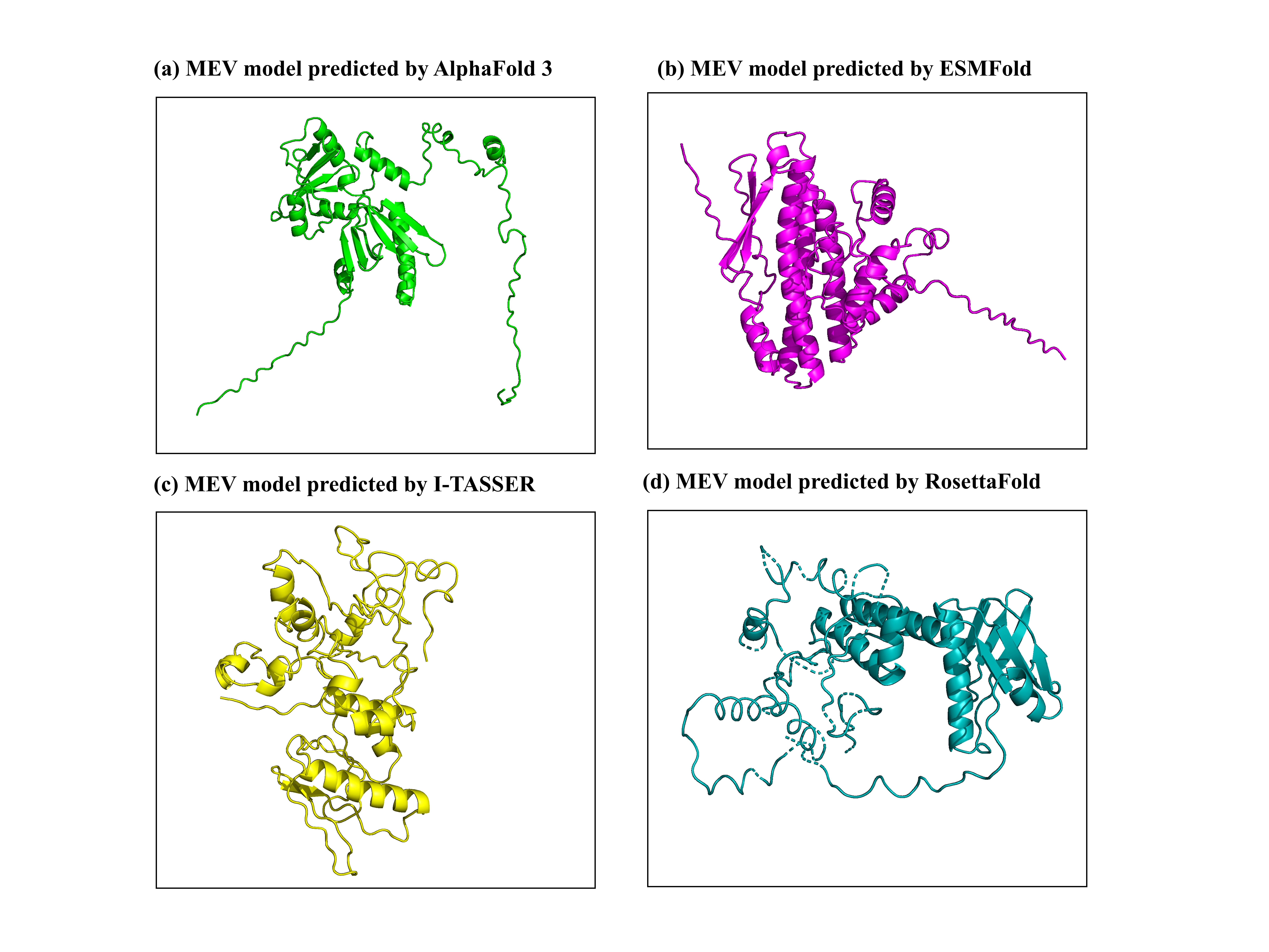

**Figure S4. Tertiary structure of the MEV construct predicted by four modeling tools. (a)** AlphaFold 3, **(b)** ESMFold, **(c)** I-TASSER, and **(d)** RosettaFold.

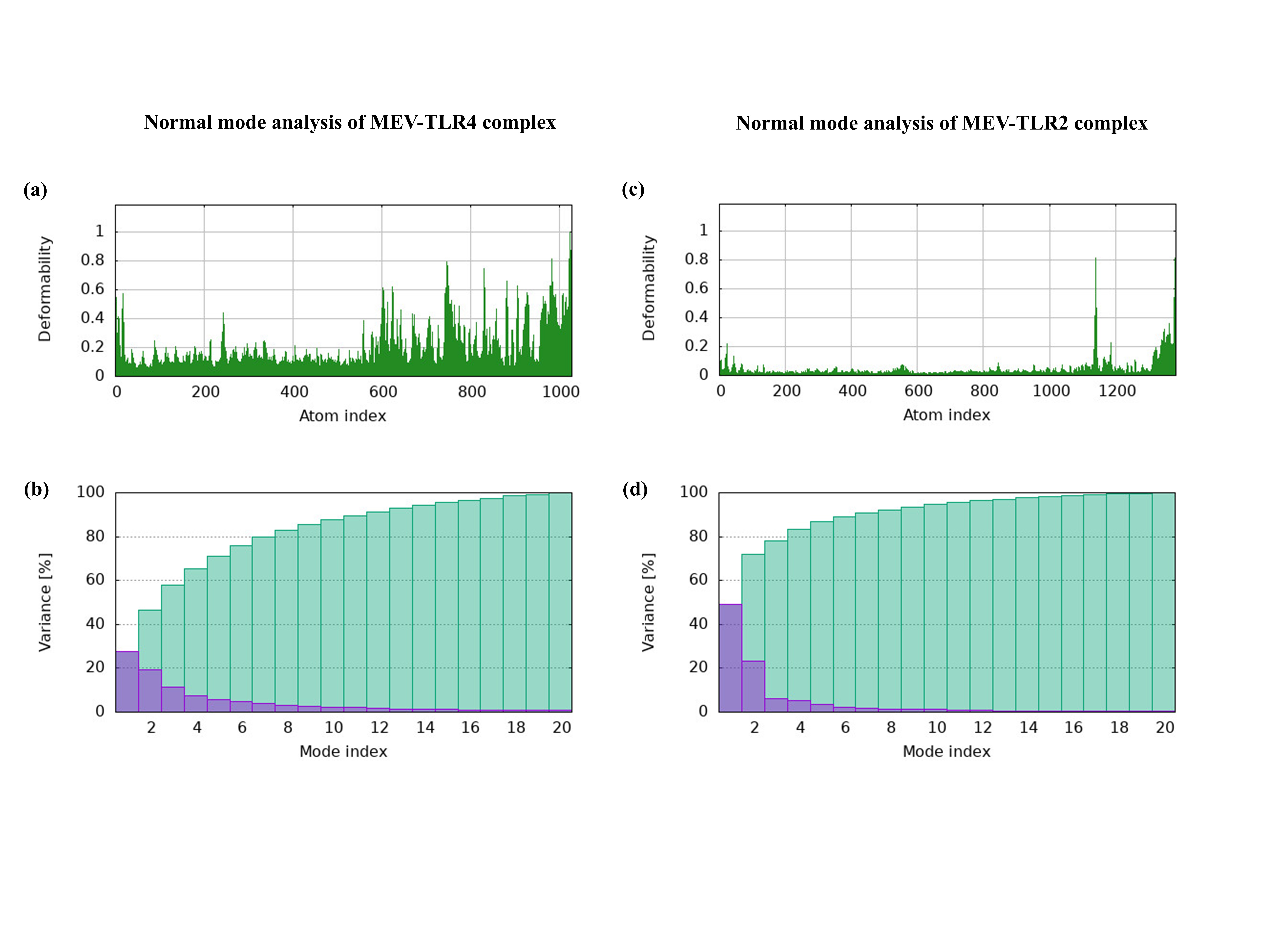

**Figure S5. Normal mode analysis of MEV-TLR4 and MEV-TLR2 complexes.** **(a, c)** Deformability plot of both MEV-TLR4 and MEV-TLR2 shows low deformability with localized flexible peaks. **(b, d)** The variance analysis depicts the first few low-frequency modes contributed to the overall conformational motion and the whole complexes were in coordination by the end of 20 modes rather than random structural movements.

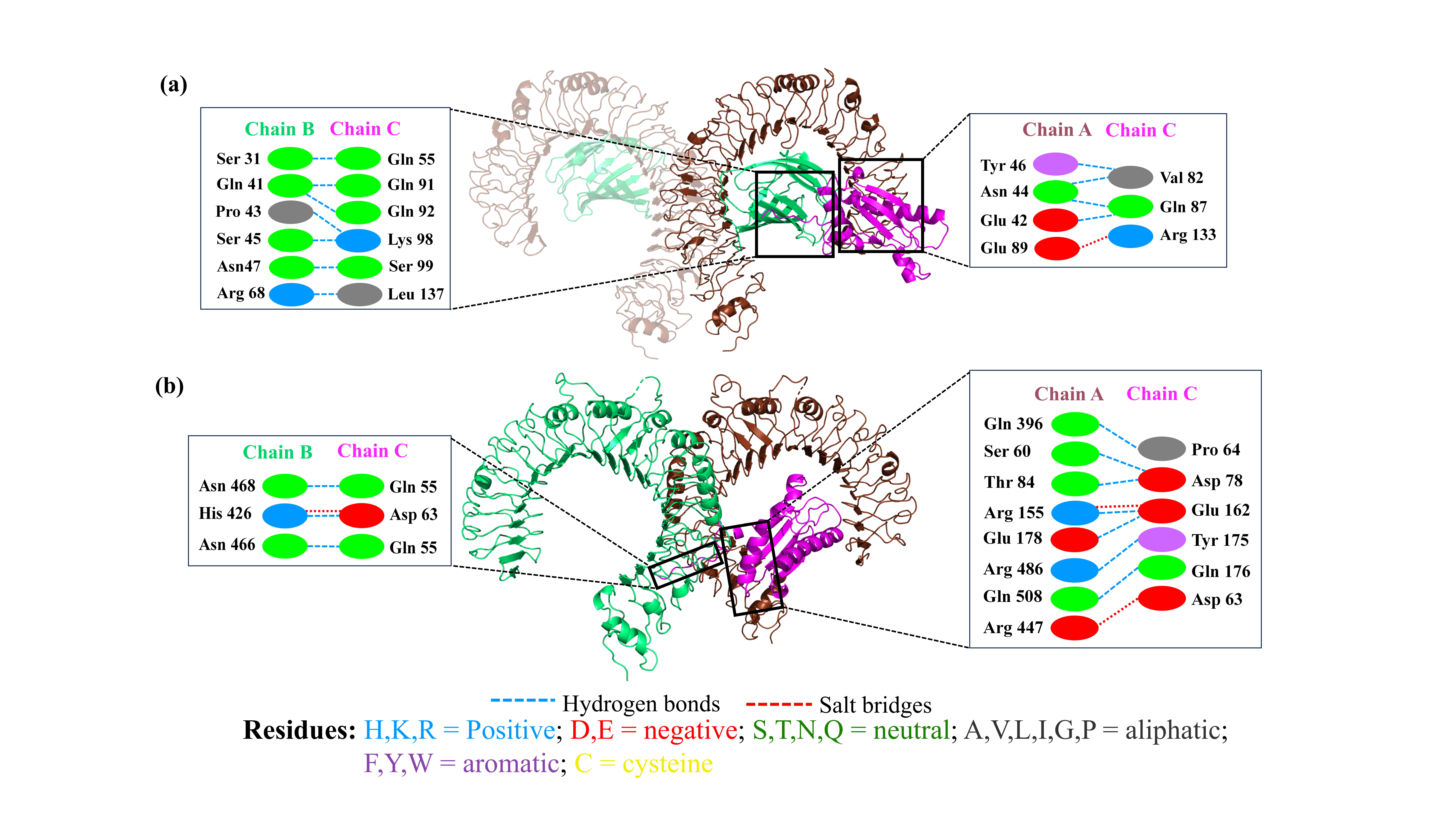

**Figure S6. Interaction analysis of the docked Omp7 with human TLR4 and TLR2.** (a) The docked complex of the Omp7 and TLR4 is shown. The binding interactions between the complex predicted formation of 11 hydrogen bonds and one salt bridge. (b) Omp7-TLR2 complex formed ten hydrogen bonds and three salt bridges at protein-protein interface.

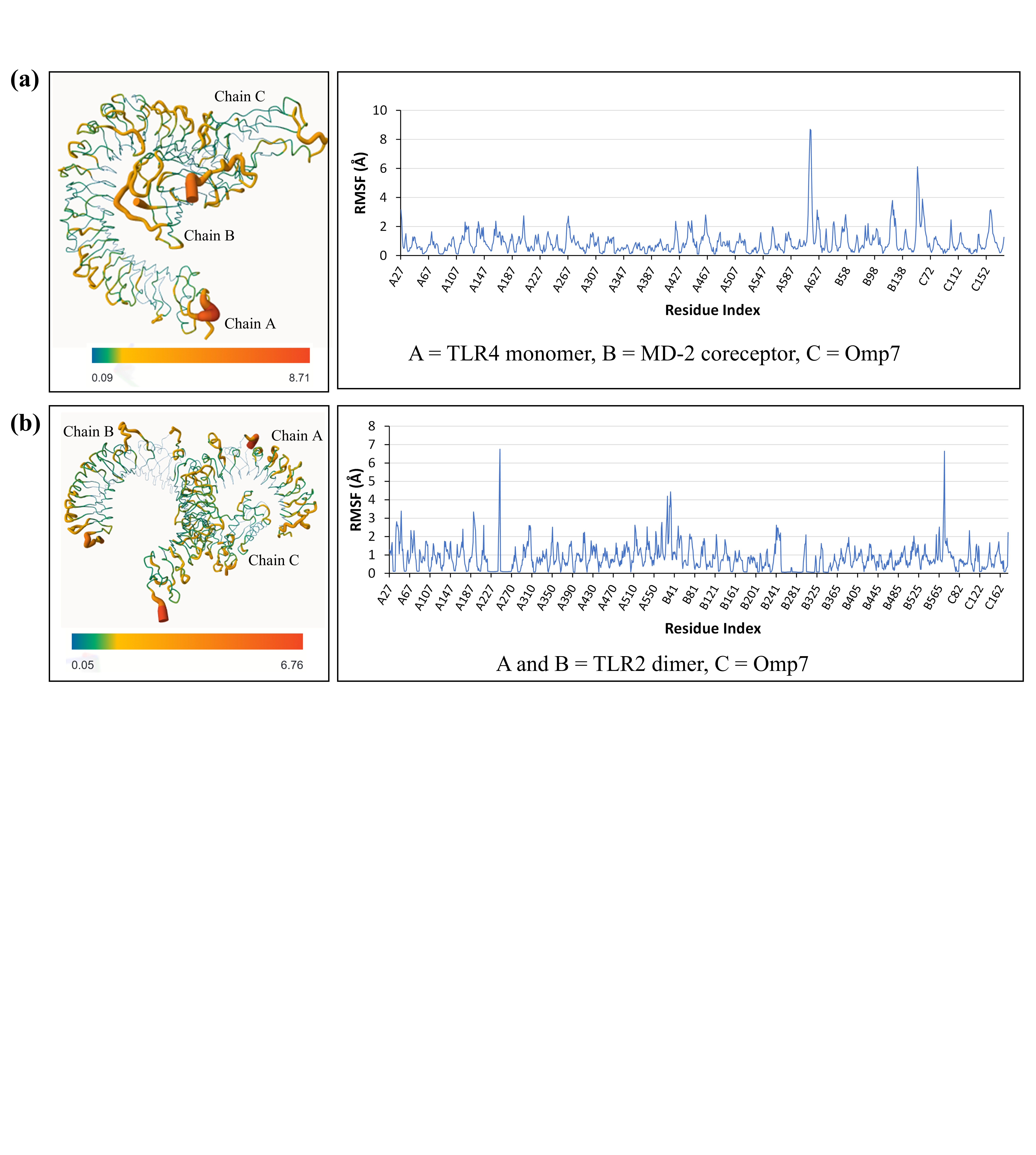

**Figure S7. RMSF profiles of** **Omp7-immune receptor complexes (positive control). (a)** RMSF profile of MEV-TLR4 complex throughout the simulation. **(b)** RMSF-based model and graph showing flexibility distribution in the Omp7-TLR2 complex.

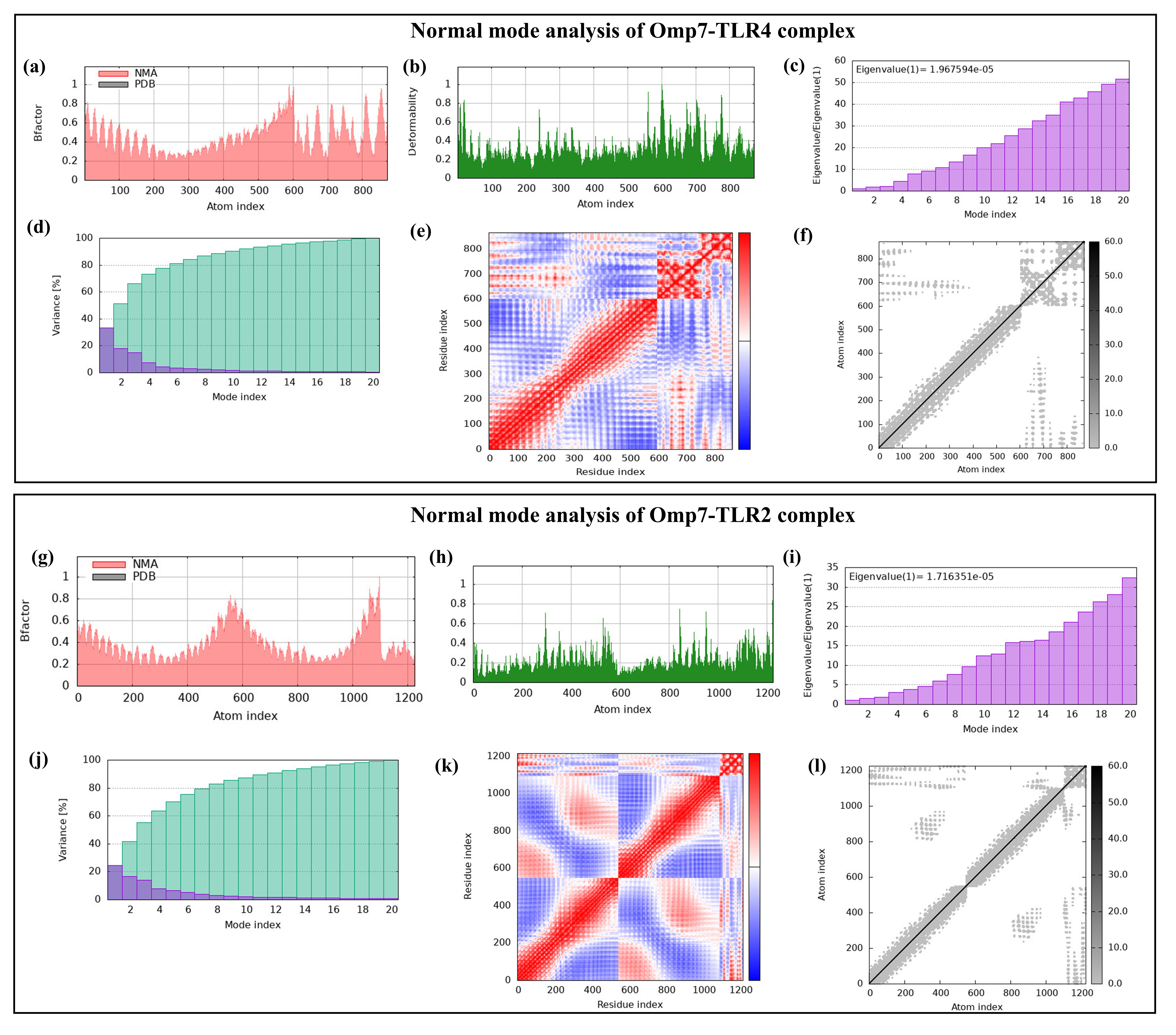

**Figure S8. Normal mode analysis (NMA) of the positive control Omp7-TLR4 and Omp7-TLR2 complexes using iMODS. (a, g)** B-factor profiles, **(b, h)** Deformability profiles, **(c, i)** Eigenvalue spectra, **(d, j)** Variance distribution, **(e, k)** Covariance maps, and **(f, l)** Elastic network models are shown.

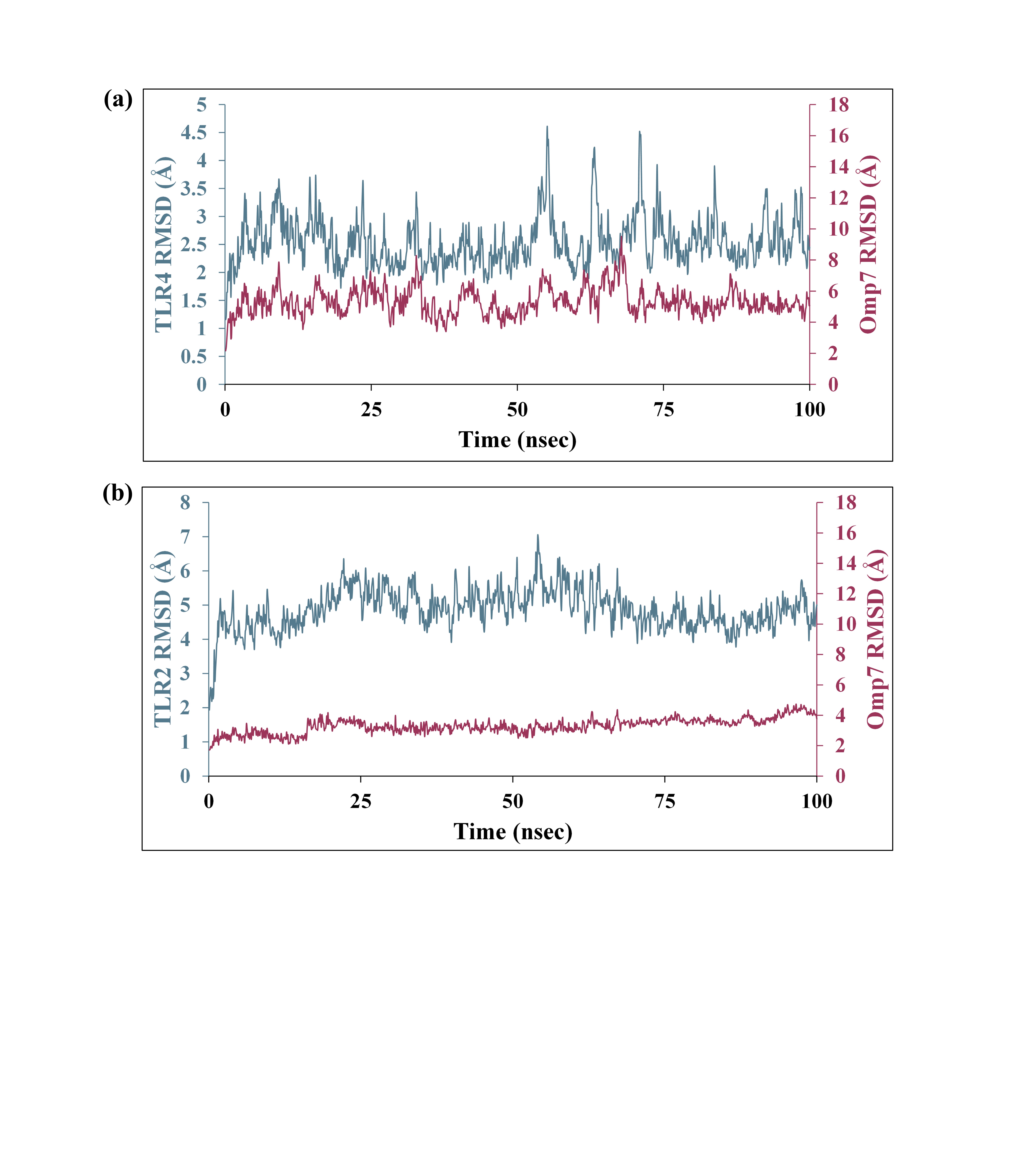

**Figure S9. RMSD profiles of Omp7-TLR4/TLR2 complexes over 100 ns MD simulation. (a)** RMSD profile of TLR4/MD-2 and Omp7. **(b)** RMSD profile of TLR2 dimer and Omp7 protein

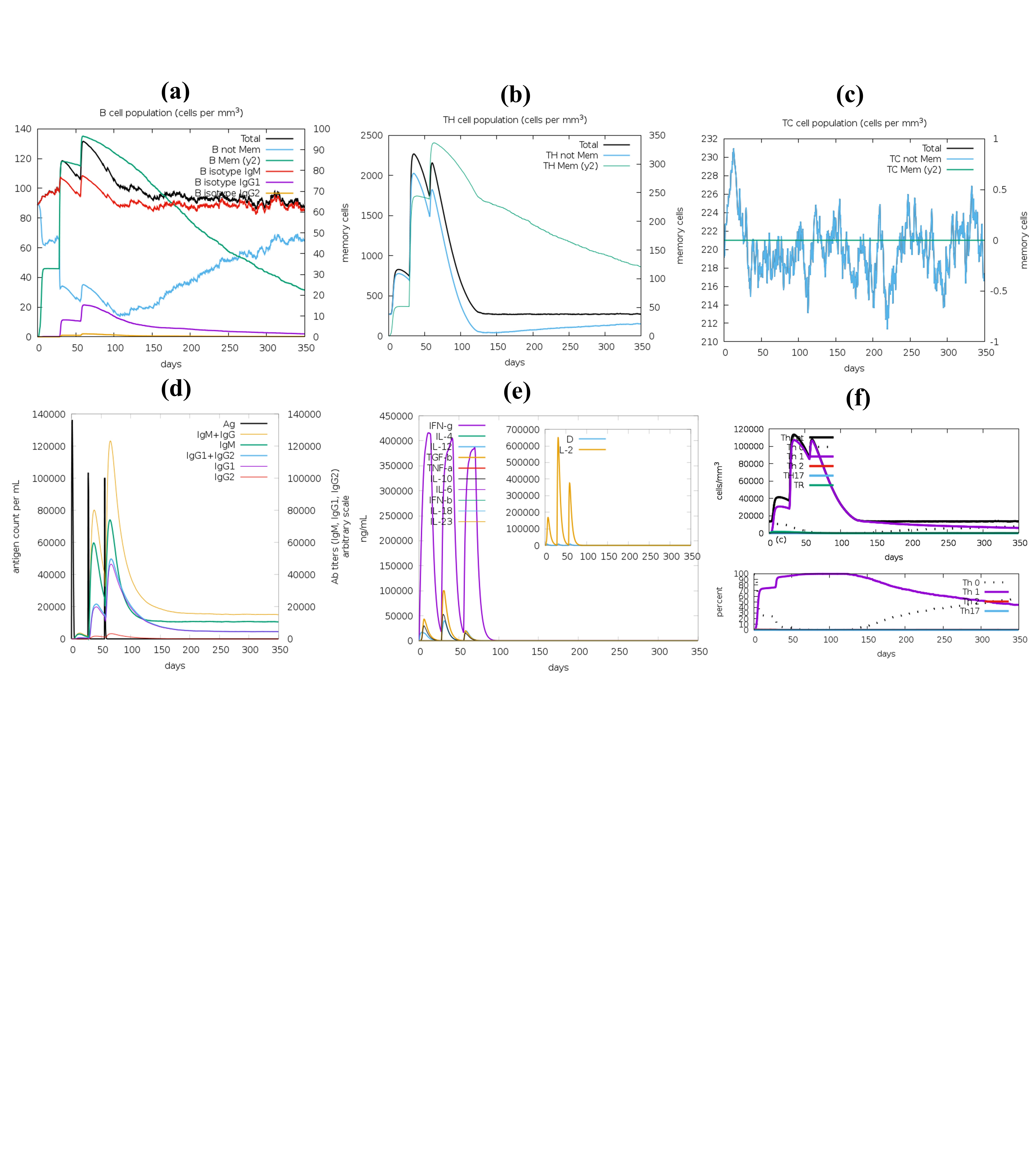

**Figure S10. In-silico immune simulation of Omp7 using C-ImmSim server. (a)** B-cell isotypes in various states. **(b)** Helper T-cell population. **(c)** Cytotoxic T-cell population. **(d)** Antigen and subtypes of immunoglobulin levels. **(e)** Concentration of cytokines and interleukins at three different stages (8 hrs, 28 days, and 56 days). **(f)** Helper T-cell isotypes in various states.
